# BiOmics: A Foundational Agent for Grounded and Autonomous Multi-omics Interpretation

**DOI:** 10.64898/2026.01.17.699830

**Authors:** Lei Cao, Yuntain Li, Hua Qin, Yanbang Shang, Yilin Zhang, Bogdan Jovanovic, Lazar Djokic, Tianyi Xia, Luni Hu, Haiyang Hou, Xingxing Ning, Li’ang Lin, Hao Qiu, Ziqing Deng, Yuxiang Li, Yong Zhang, Shuangsang Fang

## Abstract

While AI has automated bioinformatic workflows, biological interpretation remains fragmented and often disconnected from mechanistic insights. Existing AI is bifurcated between statistical “black-box” models that lack logical grounding and simple agents restricted to shallow knowledge retrieval. To bridge this divide, we introduce BiOmics, a foundational agent that synthesizes multi-omics data with adaptive knowledge for biological interpretation. BiOmics introduces a novel dual-track architecture comprising a harmonized explicit reasoning space for grounded logic and a unified latent embedding space for high-dimensional association mapping. This architecture enables a transformative “Retrieving-Reasoning-Predicting” paradigm for purposeful, cross-scale inference traversing the biological hierarchy, from molecular variants to disease phenotypes. Empirical evaluations demonstrate that BiOmics surpasses state-of-the-art AI agents and specialized algorithms, markedly augmenting the granularity and depth of biological insights. Specifically, BiOmics exhibits unique superiority in uncovering indirect pathogenic variants, achieving reference-free cell annotation, and prioritizing drug repurposing candidates tailored to specific datasets. BiOmics further enriches the interpretive landscape of biological entities, leveraging its reasoning-grounded knowledge graph to uncover deep functional contexts. Ultimately, BiOmics provides a versatile engineering foundation to transition AI for Science from descriptive “data fitting” to autonomous, knowledge-driven interpretation.

## Introduction

The rapid maturation of high-throughput multi-omics technologies has ushered in an era of unprecedented data abundance, yet the fundamental bottleneck of modern biology has shifted from data generation to mechanistic interpretation[1, 2]. The traditional paradigm, characterized by a disjointed “analysis-first, interpretation-later” workflow, creates a formidable epistemic divide between computational outputs and biological insights (**Fig. 1a left**)[3]. While artificial intelligence (AI) has been recruited to bridge this gap, current systems face a critical dichotomy between data-driven representation and knowledge-driven reasoning.

**Fig1.**
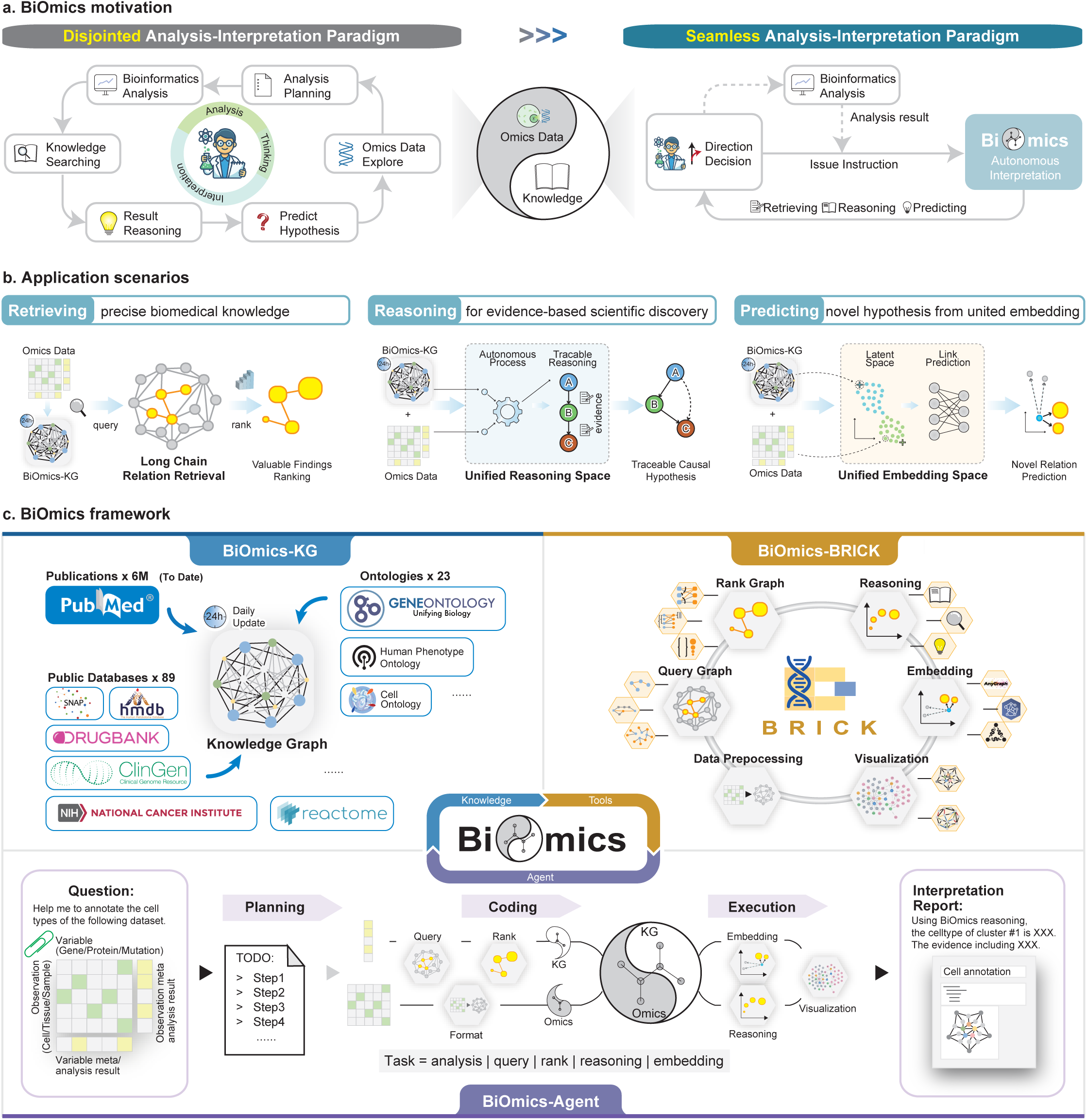
Overview of BiOmics system. a. BiOmics motivation. BiOmics aims to integrate omics data with prior knowledge, simplifying the complex process of traditional omics knowledge discovery into a process achievable solely through human-computer interaction. This enables rapid and automated mining of omics data. **b. Application scenarios.** BiOmics interpret omics data from three aspects, retrieval accurrate bio-medical knowledge, reasoning the evidence-based scientific discovery and predicting new hypothesis from united embedding. **c. BiOmics framework.** The BiOmics framework compose of BiOmics biomedical knowledge graph (left top), BRICK toolkit (right top) and BiOmics multi-agent system (bottom). BiOmics knowledge graph (KG) integrate ontologies, public databases and publications. BRICK, a configurable modular toolkit for using knowledge graph and omics data, contains six modules including data preprocessing, query graph, rank graph, reasoning, united embedding and visualization. BiOmics multi-agent system automatically planing interpretation steps, calling BRICK tools, execute BRICK codes and generate interpretation report based on user’s question and omics data. The integration of knowledge graph, tool calling, and multi-agent system lays the foundation for biological interpretation tasks.

Recent breakthroughs in cell foundational models, most notably GeneFormer[4] and scGPT[5], have demonstrated the transformative power of pre-training on massive single-cell transcriptomes to capture latent biological representations. However, these models predominantly operate as “statistical black boxes”, excelling at pattern recognition and feature extraction but lacking the explicit logical grounding required for mechanistic inference[6]. While they can predict cellular states, they cannot explain the underlying causal chains or integrate real-time scientific knowledge. Furthermore, these foundational models are largely confined to gene-level embeddings, struggling to traverse the multi-scale complexity that connects granular molecular perturbations to systemic clinical phenotypes.

Complementing these representation models, a new generation of biomedical AI agents has emerged to automate data workflows[3]. Systems such as AutoBA[7], BioMaster[8], and CellAgent[9] focus on the execution of bioinformatic pipelines; however, their utility typically terminates at intermediate statistical products without resolving their biological significance. Similarly, While Biomni[10] integrates numerous tools for broad tasks, it relies on shallow tool-result mapping without deep reasoning support. GeneAgent[11] is capable of parsing gene functions but constrained to gene-level entities. Fundamentally, the field lacks a foundational biological reasoner, a system that does not merely fit data but actively reasons through it by coupling high-dimensional embeddings with structured biological laws.

The integration of structured knowledge also faces a crisis of rigidity versus generalizability. Current knowledge graph (KG)-omics fusion frameworks, such as CKG[12] and BioKG[13], offer automated retrieval and knowledge discovery but rely on static workflow. While BioCypher[14] and SPOKE[15] enhance knowledge accessibility, they remain isolated from real-time bioinformatic workflows. Consequently, these approaches predominantly operate in a “static graph and fixed query” mode, failing to establish a closed loop that unifies data-driven path planning, real-time extraction and automatic reasoning.

To resolve these dilemmas, we introduce BiOmics, a foundational agent engineered to align multi-omics data with adaptive knowledge for biological interpretation (**Fig.1a right**). At its heart, the framework operates through a dual-engine core comprising a harmonized explicit reasoning space for formal logical inference and a unified latent embedding space for predictive latent representation. By synergizing these two modules with long-chain knowledge retrieval, BiOmics executes a transformative “Retrieving-Reasoning-Predicting” paradigm (**Fig.1b**). This operational logic is realized through a tripartite framework consisting of Knowledge (BiOmics-KG), Tools (BiOmics-BRICK), and Agents (BiOmics-Agent) (**Fig. 1c**). Specifically, BiOmics-KG provides a foundational memory of 350 million daily-updated relations to ground inference and mitigate stochastic hallucinations; BiOmics-BRICK offers a modular, pluggable toolchain to overcome bioinformatic interoperability bottlenecks; and BiOmics-Agent serves as the logical orchestrator for high-order autonomous path planning and hypothesis generation. Collectively, this infrastructure establishes a foundational capacity for adaptive, cross-scale inference that traverses the biological hierarchy from granular molecular variants and regulatory signals to macroscopic disease phenotypes.

Empirical evaluations demonstrate that BiOmics surpasses state-of-the-art AI agents and specialized algorithms, markedly augmenting the granularity and depth of biological insights. Specifically, BiOmics exhibits unique superiority in uncovering indirect pathogenic variants, achieving reference-free cell annotation with fine-grained subpopulation resolution, and prioritizing drug repurposing candidates tailored to specific disease datasets. BiOmics further expands the interpretive landscape of biological entities through its reasoning-grounded knowledge graph. To illustrate, in gene-centric analysis, it elevates traditional pathway enrichment into comprehensive phenotypic profiling, systematically decoding the intricate links between genes and their roles in tissue specificity, disease pathogenesis, and cellular homeostasis. By establishing a versatile engineering foundation that couples data-driven discovery with knowledge-guided reasoning, BiOmics provides the essential bridge to the next generation of interpretable and precision-driven AI for Science (AI4S).

## Results

### BiOmics system overview

The BiOmics framework leverages the compatibility of graph structures to deeply integrate prior biological knowledge with high-throughput omics data, constructing a explicit reasoning space and a latent embedding space. The harmonized explicit reasoning space provides a logical framework for the “input–fusion–conclusion” pipeline, inferring biologically consistent results while supporting retrospective reasoning-chain validation to rectify contradictions with existing knowledge. Meanwhile, the unified latent embedding space fuses multi-omics data with precisely retrieved knowledge entities to eliminate modality barriers.

The innovation of BiOmics lies in two aspects: integrating prior knowledge into the analysis process to correct results and enhance reliability, and reverse-mining hidden data patterns to enable dynamic updating and rediscovery of prior knowledge. For cell multi-omics data analysis, BiOmics mainly implements three core functions (**Fig. 1b**): (i) Knowledge retrieval: Based on its daily-updated knowledge graph, it supports long-chain association mining of diverse entity relationships. (ii) Causal reasoning: It autonomously integrates omics data information with knowledge graph-derived prior knowledge to generate traceable reasoning results in a harmonized explicit reasoning space. (iii) Association prediction: It embeds the insight of omics data and associated knowledge into a unified latent embedding space, acquires unified embeddings via graph representative learning models or large-model feature fusion, and enables association prediction through diverse downstream tasks. Composed of the daily-updated knowledge graph (BiOmics-KG), pluggable toolkit for interpretation (BiOmics-BRICK), and multi-agent system with reasoning ability (BiOmics-Agent), the BiOmics system provides a foundation for combined reasoning and association prediction of data and knowledge (**Fig 1c**).

**BiOmics-KG** BiOmics-KG provides a high-quality semantic foundation for diverse downstream tasks by systematically integrating multi-source authoritative knowledge. Specifically, BiOmics-KG uses 23 international mainstream biological ontologies[16] (covering GO[17], DOID[18], CL[19], Uberon[20], etc.) as the core skeleton, mapping 89 public databases (UniProt[21], DisGeNet[22], Reactome[23], etc.), and incorporating approximately 6 million PubMed articles with an impact factor ≥4 from 2004 to the present (**Fig. 1c** left top, **Supplementary Fig. 3**). BiOmics-KG includes three modules: entity standardization, relationship normalization, and timely updates. Compared to existing knowledge graphs, it features: (i) The largest knowledge graph volume: As of the manuscript drafting date, BiOmics-KG has 10,882,055 nodes and 356,017,954 relationships (**Supplementary Fig. 4**). (ii) High data quality: Through node merging and de-redundancy operations, approximately 140k nodes were merged, and 376k core biological entity nodes were matched with semantic vectors, improving data quality and hybrid retrieval recall (**Supplementary Fig. 5, 6, 7, Methods**). (iii) Real-time updates: We built a fully automated literature collecting-cleaning-extraction pipeline triggered by PubMed daily updates, combined with incremental entity extraction and normalization, achieving zero-human-intervention daily knowledge updates (**Methods**). Compared to existing biomedical knowledge graphs, BiOmics-KG is larger, covers wider data, has high timeliness, and ensures high quality and reliability through strict quality control (**Supplementary Table 1**)[12, 15, 24–29]. **BiOmics-BRICK** BiOmics-BRICK offers a modular, pluggable toolchain to overcome bioinformatic interoperability bottlenecks. BiOmics-BRICK is composed of six pluggable tool modules: Data Preprocessing (Standardization), Querying, Ranking, Reasoning, Graph Representation Learning (Embedding), and Visualization (**Fig. 1c** right top, **Supplementary Fig. 2b, Methods**). The Agent assembles tools from each module on demand according to user needs, forming an end-to-end pipeline that couples omics analysis results with prior knowledge in the knowledge graph in real-time, efficiently distilling key information and potential biological discoveries. The specific modules are:

- Data Processing Module: Implements format conversion between omics data and queried knowledge results, data preprocessing, construct unified sub-graph, and autonomous bioinformatic execution.
- Query Module: Responsible for diversified retrieval of associated entities, relations, neighbor nodes, and paths in the graph using differential expressed genes, variant sites, or protein nodes as anchors.
- Ranking Module: Filters and recommends candidate results based on statistical significance or prior weights.
- Reasoning Module: Transforms prediction tasks into classification or enrichment problems and construct inference trajectory to recommend potential high-value knowledge.
- Representation Learning Module: Jointly embeds omics data sub-graph and knowledge sub-graph via graph neural networks, achieving association prediction by completing unknown relationships in the joint graph.
- Visualization Module: Reconstructs key visualizations of analysis results, achieving intuitive and simultaneous presentation of data itself and knowledge interpretation results.

**BiOmics-Agent** BiOmics-KG and BiOmics-BRICK consolidate the semantic foundation and scalable framework for knowledge interpretation; BiOmics-Agent relies on the universality and generalization capability of large language models (LLMs) to achieve autonomous interpretation and discovery of biological knowledge. Its main core functions can be summarized as six items (**Fig. 1c** bottom, **Supplementary Fig. 1, Methods**):

- Requirement Parsing: Maps user requirements in natural language and multi-omics data to a shared multi-modal reasoning space to complete intent and omics data alignment.
- Scheme Generation: Retrieves and orchestrates BiOmics-BRICK components via retrieval-augmented generation (RAG) to build an end-to-end reasoning pipeline, fusing knowledge graph and omics evidence[30].
- Planning and Execution: The agent constructs an analytical workflow predicated on data meta-information and prior successful heuristics. Within a dockerized sandbox environment integrated with the Jupyter kernel gateway[31], the agent generates and executes code in a modular, step-wise fashion. This approach facilitates real-time tracking, precise debugging, and computational efficiency by bypassing redundant execution of validated code blocks.
- Result Compilation: Automatically generates complete, comprehensive, and traceable text reports, highlighting key biological discoveries.
- Human-Agent Interaction: Supports multi-turn dialogue for intent clarification, task refinement, and intermediate result confirmation.
- Memory Retention: Adopts persistent session memory to record decision history and analysis context, ensuring session consistency and continuity[32].

Through innovative mechanism design, BiOmics achieves the foundation for deep coupling of omics data and prior biological knowledge based on these three core components, building a critical bridge connecting data-driven discovery and knowledge-driven reasoning.

### BiOmics sets new benchmarks for knowledge-grounded discovery and multi-scale omics interpretation

To evaluate the information retrieval performance of BiOmics, we benchmarked its accuracy using the Biomix dataset across True/False Questions (TFQ) and Multiple-choice Questions (MCQ)[33]. BiOmics achieved a 91.32% accuracy on TFQs, representing a 6.43% improvement over the vanilla LLM. In MCQ tasks, it reached an accuracy of 66.01%, which further escalated to 76.44% for queries where relevant information was indexed within the knowledge graph. This demonstrates a significant performance gain over base models and Biomni, surpassing GPT-4o by approximately 6% and Biomni by 13% (**Fig. 2a**). Furthermore, to assess response granularity, we constructed a test set of 100 questions (25 each for cells, diseases, genes, and variants, e.g., “What is BARC1?”). BiOmics exhibited superior semantic richness, yielding significantly more biological entities than base LLMs (≈92% increase) and Biomni (≈208% increase), with all retrieved entities fully traceable to verified references or databases (**Fig. 2b**).

**Fig2.**
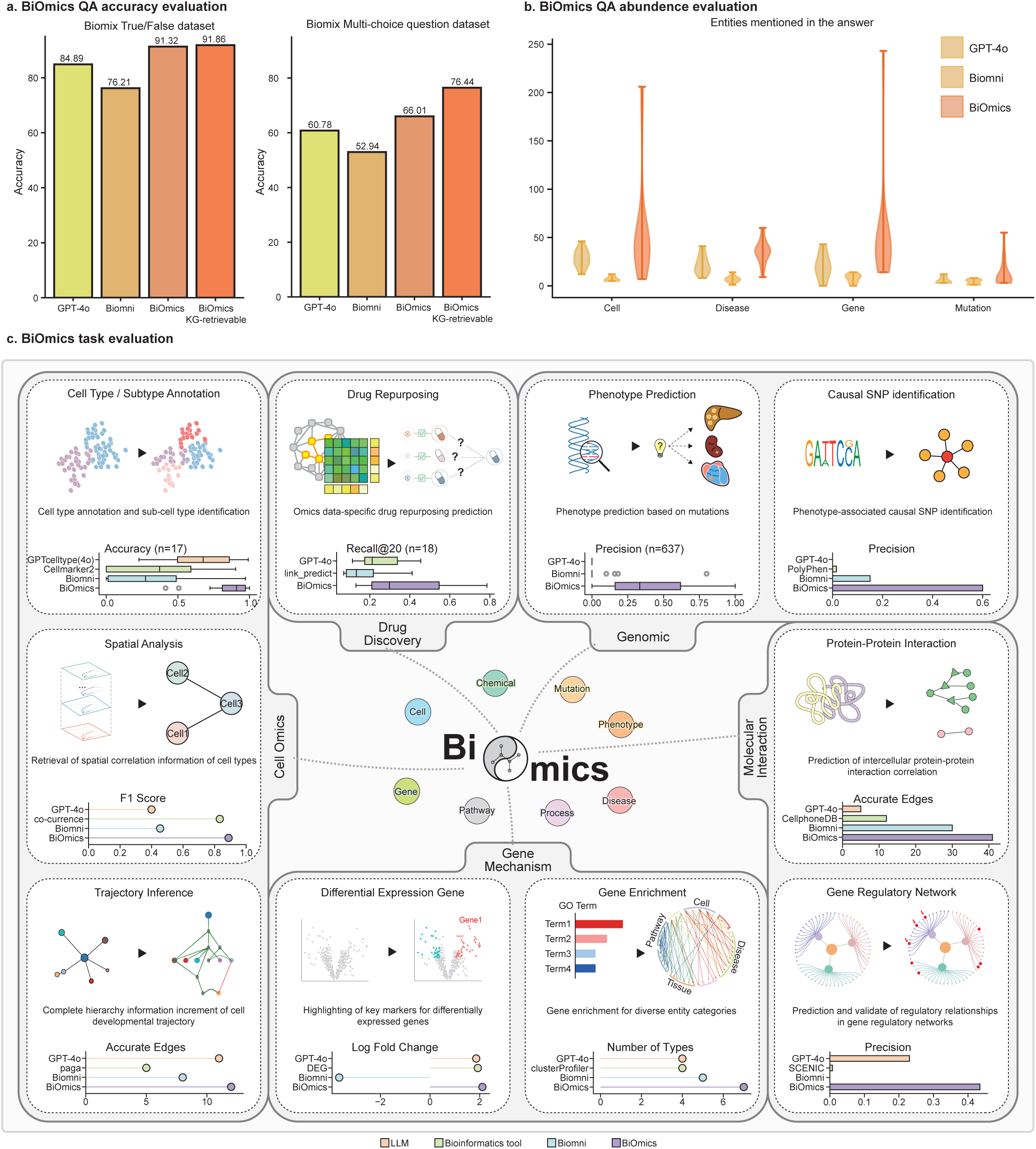
Benchmark of BiOmics. a. BiOmics QA accuracy evaluation. Using Biomix QA dataset, we evaluate Accuracy of questions (left: True or False dataset; right: multi-choice question dataset).Alternative methods including GPT-4o and Biomni, the last column indicate to accuracy of BiOmics within the questions that BiOmics queried information from knowledge graph. **b. BiOmics QA abundency evaluation.** y-axis: the entities reported in answers. **c. BiOmics bioinformatics task evaluation.** 10 bioinformatics task arraged in each box. In each box, There is a diagram to indicate the main difference of BiOmics and traditional analysis, key points to highlight BiOmics innovation and the evaluation comparing to LLM method, traditional bioinformatics methods and Biomni. See methods for more details.

Simultaneously, we validated the integrated query, reasoning, and predictive capabilities of BiOmics in complex multi-omics interpretation tasks across domains including variants, genes, proteins, cells, and drugs. We employed GPT-4o, task-specific bioinformatics algorithms, and the specialized agent Biomni[10] as comparative baselines. Benchmarked against ground-truth labels, BiOmics consistently outperformed all established methods (**Fig. 2c**). Key findings include:

- Phenotype prediction: In predicting phenotypes from 639 TCGA SNP samples, BiOmics correctly identified 151 samples, achieving a 13-22% relative performance gain over Biomni and GPT-4o.
- Cell annotation: Across 17 distinct tissue types (e.g., heart, liver, spleen, lung, kidney) in human and murine models[34], BiOmics attained a mean accuracy of 0.856 ± 0.165 (n=17), surpassing representative baselines by 17–54% (GPTcelltype: ≈17%; Cellmarker2: ≈54%; Biomni: ≈53%) (**Supplementary Fig. 8**).
- Drug repurposing: The average top-20 hit rate reached 0.772 ± 0.136 (n=18), exceeding traditional link prediction tools and vanilla LLMs by 42% and 21%, respectively.
- Pathogenic variant identification: A significantly higher proportion of pathogenic variants identified by BiOmics were validated in the GWAS Catalog[35, 36], with precision exceeding Biomni by approximately 40%.
- Network analysis (Gene regulatory network (GRN) / Protein-protein interaction (PPI)): For gene-regulatory and protein-protein interaction tasks, pairs identified by BiOmics showed higher validation rates in TRRUST[37] and StringDB[38]; accuracy improved by 20% (GRN task compared to vanilla LLMs) and 36% (PPI task compared to Biomni).

Beyond predictive accuracy, we evaluated BiOmics’ capacity to enhance information richness and completeness in complex reasoning tasks (**Fig. 2c**):

- Cellular trajectory inference: BiOmics successfully supplement valuable intercellular developmental relationships, increasing the edge count by 140% compared to PAGA[39] while maintaining higher edge-level accuracy than other tools (Biomni, GPT-4o, etc.).
- Differential expression analysis: The framework suggests marker genes reported in the knowledge graph that correlate with higher log-fold changes in the empirical data.
- Enrichment analysis: BiOmics facilitates simultaneous enrichment across seven biological conceptual categories, providing a more holistic interpretation than Biomni.
- Spatial co-localization analysis: BiOmics enables the high-resolution interpretation of spatial co-localization and functional relationships between distinct cell types.

### BiOmics enables precise bidirectional inference from mutations to phenotypes

To systematically evaluate BiOmics’ “genotype-to-phenotype” reasoning capability, we applied BiOmics to Type 2 Diabetes (T2D) GWAS data, which identified 290 variants significantly associated with T2D (P≤1×10DD)[35]. BiOmics performed automated retrieval and integration of variant information directly and indirectly associated with T2D (**Fig. 3a**). Specifically, BiOmics retrieved 37 variants previously reported to have direct links to T2D (**Fig. 3b, c**) and constructed a a multi-level relationship network centered on GWAS-identified variants (**Fig. 3d**). Notably, T2D was reasoned to be the most relevant phenotype for this GWAS, demonstrating the robust unsupervised disease prioritization capability of BiOmics. Entities clustering within the same subgraph as T2D also included critical clinical features such as “Glucose Tolerance Test”, “coronary artery disease”, and “Body Mass Index”.

**Fig3.**
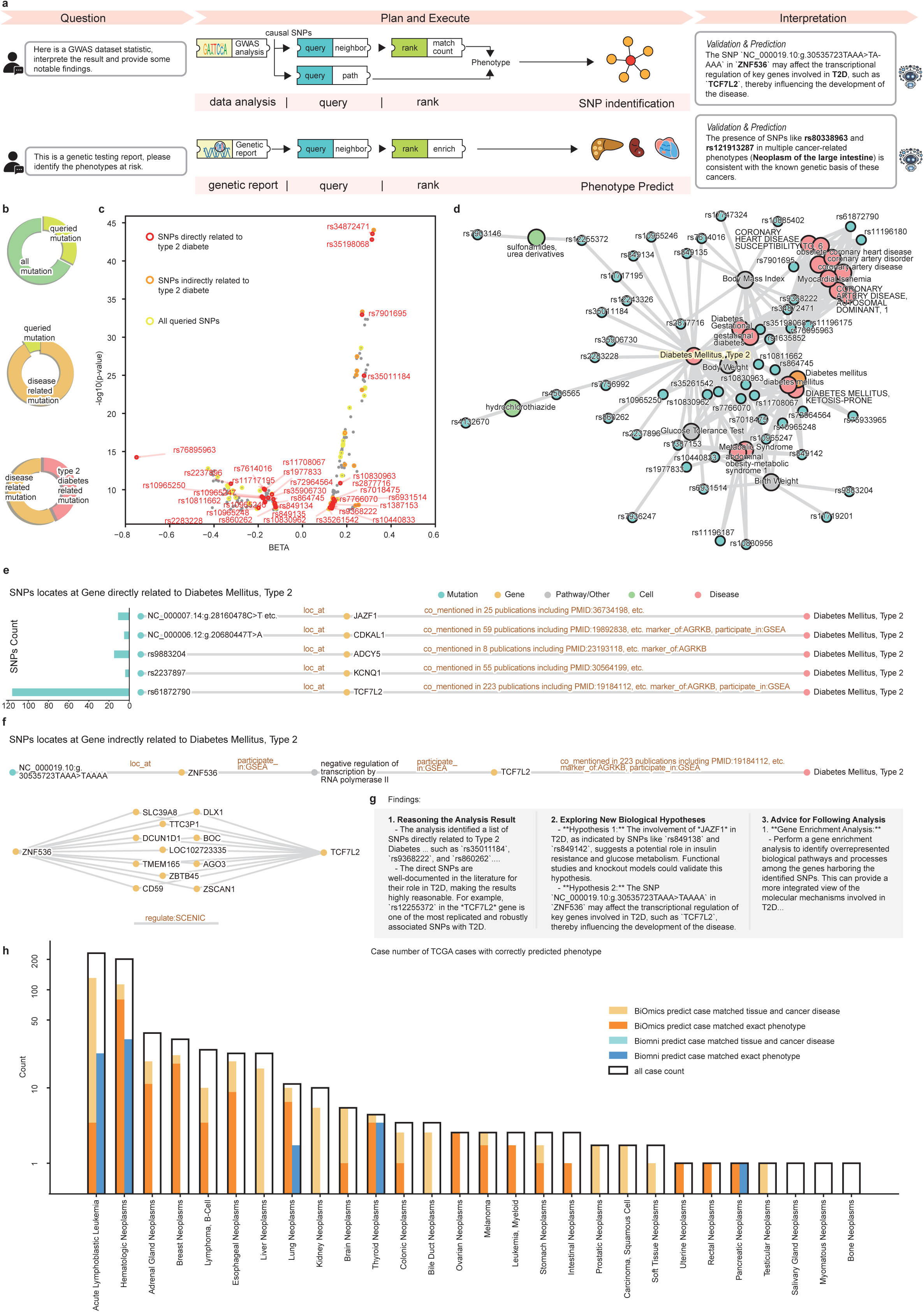
BiOmics supports precise bidirectional inference from genotype to phenotype. **a.** BiOmics pipeline for enhancing traditional omics analysis in case of causal SNP identification. **b.** Pie chart of causal SNPs among all SNPs. **c.** scatter of causal SNPs. x-axis indicate the BETA of GWAS analysis; y-axis indicate −log10(p-value); yellow, orange and red circles indicate all queried SNPs, SNPs indirectly related to type 2 diabetes and SNPs directly related to type 2 diabetes, respectively. **d.** visualization of queried graph. **e.** SNPs that locates at Gene related to T2D, the bar plot on the left indicate the SNPs count locate at corresponding gene in each row. **f.** SNPs locates at gene indirectly related to T2D. upper: the queried path from *ZNF536* gene to T2D, lower: the regulate genes between *ZNF536* and *TCG7L2*. **g.** abstract of BiOmics interpretation report. h. barplot phenotype predict on TCGA cases. x-axis: all kinds of disease type; y-axis: count, which is logarithmically compressed.

Beyond direct evidence, BiOmics mapped 151 additional variants to genes essential for T2D. Their functional roles were verified by integrating MSigDB-derived[40] gene sets and PubMed literature with evidence from the Alliance of Genome Resources Knowledge Base (AGRKB)[41]. These findings covered classic susceptibility loci, including *JAZF1, CDKAL1, ADCY5, KCNQ1*, and *TCF7L2* (**Fig. 3e**). More significantly, BiOmics identified an insertion variant in *ZNF536* which, despite not being directly linked to T2D in existing reports, participates in the “negative regulation of transcription by RNA polymerase II” alongside *TCF7L2* (**Fig. 3f**). BiOmics reasoning suggests this insertion may indirectly modulate T2D progression by perturbing *TCF7L2*-mediated transcriptional regulation (**Fig. 3g**). Additionally, BiOmics interpreted key pathogenic genes at the variant-associated gene level (**Supplementary Fig. 9**).

By integrating multi-level associations, BiOmics precisely locates potential pathogenic variants and generates mechanistic hypotheses, enabling end-to-end prediction of phenotypes or disease risks based on individual mutational profiles. To evaluate generalizability, we analyzed somatic mutation data from 639 tumor samples in the TCGA cohort. BiOmics accurately classified the cancer type for 151 samples, while an additional 232 samples were correctly associated with related malignancies of the same tissue origin. In comparison, Biomni correctly predicted only 62 cases; BiOmics achieved an approximately 2.4-fold improvement in accuracy (**Fig. 3h**). In a representative case study, BiOmics not only successfully identified the colorectal neoplasm phenotype but also pinpointed key variants categorized as highly pathogenic by both PolyPhen-2[42] and SIFT[43] (**Supplementary Fig. 10**).

### BiOmics elucidates multi-scale gene and cellular landscapes in transcriptomics

To demonstrate the capability of BiOmics in deepening the interpretation of quantitative cellular and genetic data, we systematically evaluated the framework using a mouse pancreas single-cell transcriptome dataset[44]. In a reference-free cell annotation scenario, BiOmics leveraged its autonomous reasoning engine to intelligently discriminate between cell clusters based on pre-clustered data. While generating standardized cell type labels, BiOmics provided a traceable chain of evidence for each annotation (**Fig. 4a**). Quantitative evaluation revealed that BiOmics’ annotation accuracy surpassed existing reference-free methods, such as GPTcelltype[45] and CellMarker2[46], by 19-35% (**Fig. 4b, c, d**).

**Fig4.**
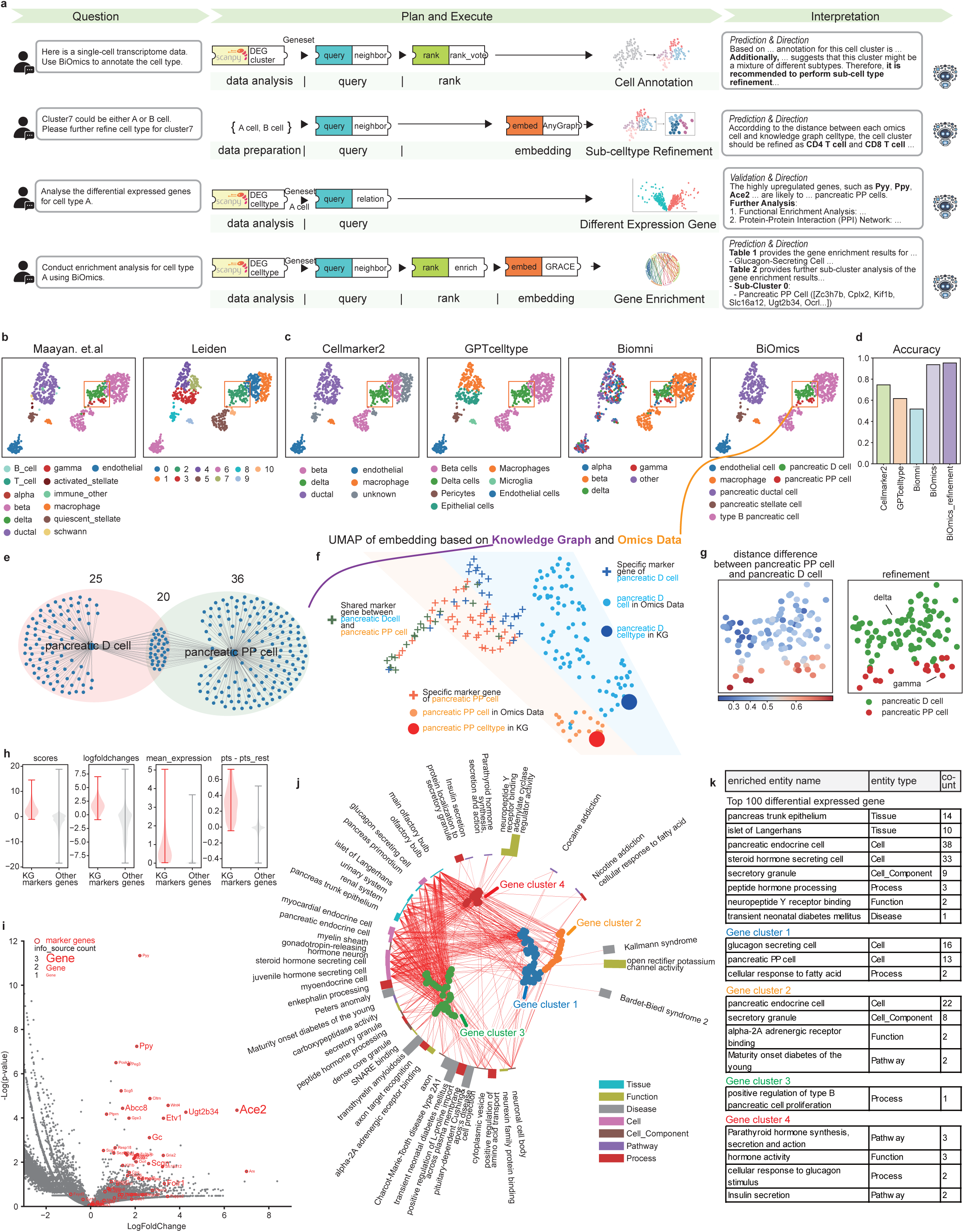
BiOmics enhances the depth of omics analysis. **a.** BiOmics pipeline for enhancing traditional omics analysis in case of cell annotation (first row), celltype refinement (second row), differential expression gene (third row) and gene enrichment (forth row). **b.** UMAP of ground truth annotation and clustering labels of mouse pancreatic islets. **c.** UMAP of BiOmics and alternative method’s annotation result. From left to the right are CellMarker2, GPTcelltype, Biomni and BiOmics result. **d.** Bar plot of accuracy of BiOmics and alternative cell annotation metrics. the last columns is BiOmics refinement result. **e.** The venn plot of queried marker of pancreatic D cell and pancreatic PP cell. Scatter plot directly display the queried graph and Venn plot visualize the marker count of each cell type. **f.** The UMAP based on unite embedding of omics dataset and KG query result. **g.** UMAP of refinement result between pancreatic D cell and pancreatic PP cell. left. the difference of euclidean distance between pancreatic PP cell and pancreatic D cell, right. refinement result. **h.** Violin plot of differential expression gene indicators for KG markers and other differential expression genes, respectively. Red: KG markers, Grey: Other genes. The indicator is scores, logfoldchanges, mean expression and difference of expression cell percentage between specific cluster and other cell clusters. **i.** Volcano plot of differential expression gene of pancreatic PP cell with known marker highlighted. x-axis: log-fold-change, y-axis: −log(p-value), each point represent a gene, known marker are circled in red and label gene name, the font size of gene name represent information source count of marker of relation between gene and pancreatic PP cell. **j.** Radial enrich plot of marker of pancreatic PP cell. The colors of the bars distinguish the types of enriched entities, and the heights of the bars represent the number of genes in the enriched entities. Meanwhile, gene sets associated with adjacent bars have higher similarity. In UMAP scatter plot in the center of the circular bar chart, each point represent a gene. The colors of the genes represent the grouping of genes. Genes that are close to each other in the UMAP also indicate a higher functional correlation. Red lines connect each gene to its corresponding enriched term. **k.** A table list enriched term of all DEG of pancreatic PP cell and each gene sub-cluster of DEG of pancreatic PP cell.

Notably, BiOmics’ reasoning logic suggested that Cluster 2 might be a mixture of pancreatic D cells (delta cells) and pancreatic PP cells (gamma cells) (**Supplementary Fig. 11**). Subsequently, BiOmics initiated an association prediction workflow, constructing a graph network integrated with omics data, relevant cell type nodes, and associated marker genes to build a unified latent embedding space (**Fig. 4e, f**). Through this approach, BiOmics successfully resolved the cluster into two distinct subgroups, each showing a high correlation with their respective pancreatic D cell and PP cell type nodes (**Fig. 4g**, **Supplementary Fig. 12a,b)**. Quantitative evaluation indicated that this BiOmics-refined approach improved accuracy by an additional 1.5% compared to the baseline version (**Fig. 4d**). Notably, the inherent physiological proximity of pancreatic D and PP cells, stemming from their common endocrine lineage, creates a transcriptomic similarity that makes them indistinguishable to traditional unsupervised tools like Leiden[47] (**Supplementary Fig. 12c**). BiOmics overcomes this bottleneck by providing a knowledge-guided subdivision axis, effectively mapping clusters to their true biological types where data-only methods fail. This achievement effectively reproduces expert-level biological reasoning under zero-reference conditions and establishes an interpretable, scalable framework for identifying novel or potential subtypes.

Next, to investigate BiOmics’ enhancement of gene-level interpretation, we applied BiOmics to perform a comprehensive differential expression analysis for pancreatic PP cell (**Fig. 4h, i**, **Supplementary Fig. 13, 14**). We then utilized BiOmics to perform multi-variate enrichment reasoning on differential genes across multiple hierarchical levels, including cellular, tissue, pathway, and disease ontologies(**Fig. 4j**). BiOmics not only precisely recovered canonical pancreatic PP cell pathways like “insulin secretion” and “peptide hormone processing”, but also captured broader contexts like “pancreas trunk epithelium” (anatomical) and “glucagon-secreting cell” (cellular), suggesting potential roles of differential genes in developmental origin and paracrine regulation.

Of clinical value, the “transient neonatal diabetes mellitus” phenotype was significantly enriched, indicating that the dysregulation of these genes may be intrinsically linked to the disease’s pathogenesis. Furthermore, BiOmics projected differential genes into a unified embedding space to quantify their functional and expression heterogeneity. By fusing omics signals with knowledge graph relationships, correlated genes autonomously organized into several “semantic clusters” in the embedding space (**Fig. 4j, Supplementary Fig. 15, 16, 17**). For instance, the green cluster was highly coupled with tissue localization and cell identity, while the blue cluster primarily associated with cellular identity features (**Fig. 4j, k**). This “hypothesis-free” clustering not only validated the biological consistency of traditional differential expression analysis but also provided an interpretable, scalable gene coordinate system for subsequent mechanism mining.

### BiOmics enhances rational prediction and verification of cellular trajectories

To demonstrate BiOmics’ reasoning verification and reflection capabilities empowered by high-confidence knowledge, we applied BiOmics to a Mus musculus leukocyte dataset to explore the differentiation pathways of the myeloid lineages[48] (**Fig. 5a**). We specifically assessed the reliability of single-cell differentiation trajectories predicted by PAGA[39] (**Fig. 5c**), which initially partitioned the dataset into three distinct cell-type clusters (**Supplementary Fig. 18**). Utilizing BiOmics to query relationships across seven primary cell types, we generated an expanded developmental lineage tree encompassing 23 cell types (**Supplementary Fig. 19**). BiOmics enhanced the trajectory analysis in three critical dimensions: (i) Root Node Inference: By determining the directionality of developmental relationships, BiOmics accurately identified the root node, a prerequisite for precise pseudotime analysis[49] (**Fig. 5b**). (ii) Lineage Reconstruction: BiOmics reconstructed complete developmental paths, effectively capturing transient intermediate states (**Fig. 5d**). (iii)Knowledge-based Validation: The framework distinguishes between established developmental paths indexed in databases (green paths) and unrecorded or divergent paths (red paths) (**Fig. 5d, Supplementary Fig. 20**).

**Fig5.**
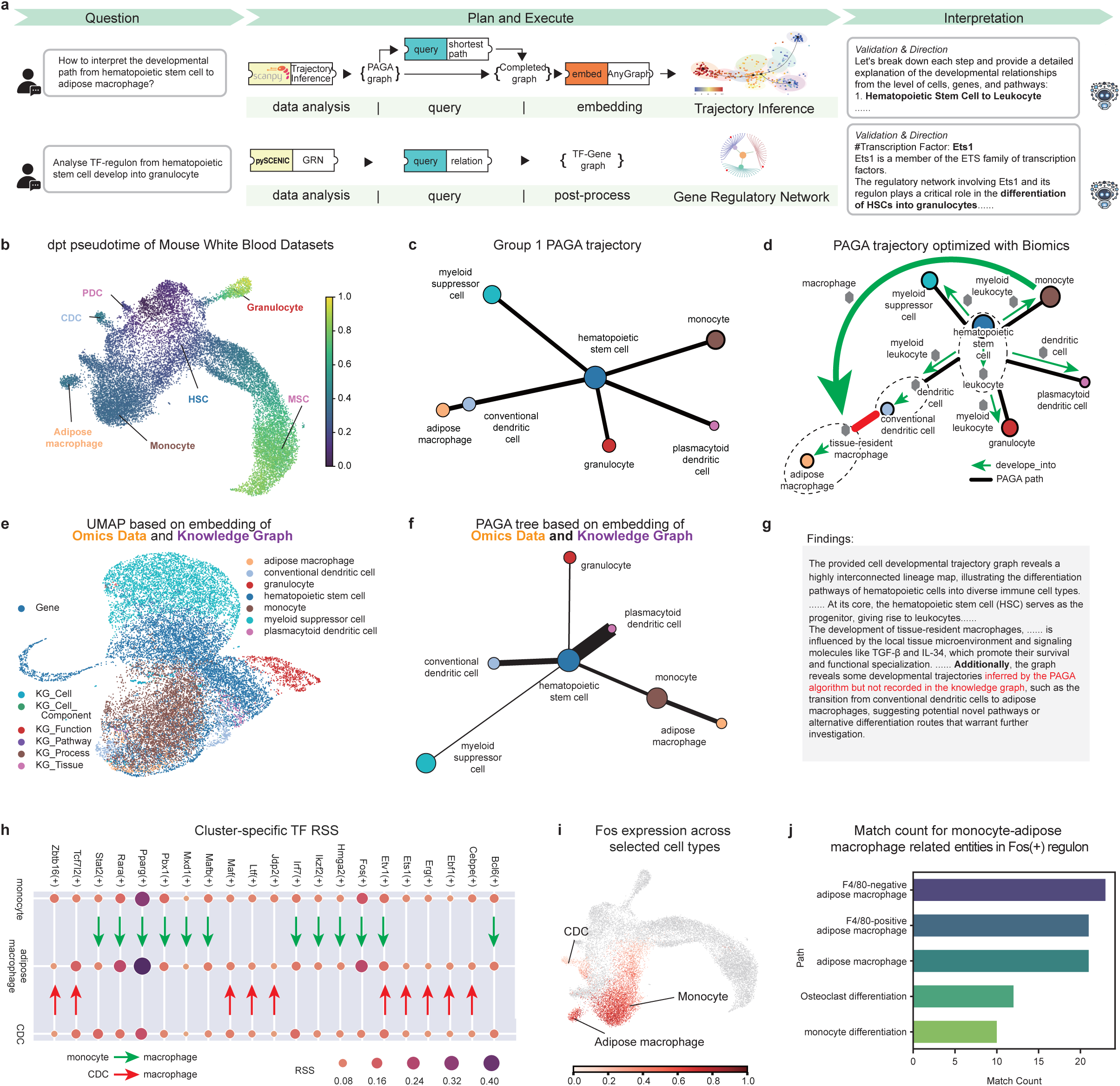
BiOmics navigates the path of omics analysis. **a.** BiOmics pipeline for enhancing traditional omics analysis in case of trajectory analysis (first row) and gene regulatory network (second row). **b.** UMAP visualization of Dpt-Pseudotime analysis result of cell group 1 with root node HSC. **c.** PAGA trajectory of WBCs dataset. **d.** The simplified trajectory graph about cell group in bone marrow (Granulocyte, HSC, MSC, PDC, CDC, Monocyte and Adipose macrophage) after BiOmics completion the PAGA graph with Knowledge graph. The colorful circle nodes represents the cell types from dataset and the other gray hexagon nodes represents the cell types from knowledge graph. The green path represents verified path and the red path represents unverified path. **e.** The UMAP based on embedding of knowledge graph and omics data based on BiOmics unite embedding. **f.** PAGA trajectory of cell group 1 based on BiOmics unite embedding of knowledge graph and omics data. **g.** The interpretation context about cell group 1 trajectory based on BiOmics. **h.** The Bubble chart of cluster-specific transcription factor regulon specificity score(RSS) among three cell types (monocyte, CDC and adipose macrophage). Arrow indicate the celltypes with more similiar RSS, green: monocyte and macrophage; red: CDC and macrophage. **i.** The UMAP of Fos(+) regulon intensity among three cell types (monocyte, CDC and adipose macrophage). **j.** Selected terms based on gene enrichment of Fos(+) regulon.

On this dataset, BiOmics identified a trajectory in the PAGA output that contradicted established biological consensus: a predicted path between adipose tissue macrophages (ATMs) and conventional dendritic cells (cDCs) (**Fig. 5d, g, Supplementary Fig. 21**). Conversely, BiOmics’ reasoning suggested a more probable developmental link between ATMs and monocytes.

To resolve this discrepancy, we utilized the representation learning module of BiOmics to re-evaluate the trajectories. By embedding cellular omics data, prior knowledge, and cell-type metadata into a unified embedding space (**Fig. 5e**), we re-ran the PAGA inference based on these integrated features. The revised results were significantly more consistent with biological knowledge, confirming that ATMs differentiate from monocytes rather than cDCs (**Fig. 5f, Supplementary Fig. 22**). To further validate the relationship between ATMs, cDCs, and monocytes, we performed GRN analysis using pySCENIC[50]. The resulting regulon activity profiles revealed that monocytes and ATMs share highly similar transcriptional regulatory signatures (**Fig. 5h, Supplementary Fig. 23, 24**). BiOmics identified that the *Fos* transcription factor as potential master regulator in the monocyte-to-ATM differentiation axis (**Supplementary Fig. 25**). Visualization confirmed that *Fos* expression and transcriptional activity were highly specific to monocytes and ATMs, while remaining minimal in cDCs (**Fig. 5i**). Functional enrichment of *Fos* target genes further corroborated their roles in ATM differentiation, reinforcing the monocyte-origin hypothesis (**Fig. 5j, Supplementary Fig. 26**). Collectively, BiOmics not only corrected the differentiation trajectory but also identified the key transcriptional drivers of the process.

### BiOmics advances omics-driven drug repurposing via dynamic knowledge integration

Traditional drug repurposing paradigms based on knowledge graphs typically formalize the task as a link prediction problem on a static network. Once constructed, these graphs remain stationary and cannot dynamically assimilate real-time omics signals, thereby failing to uncover novel biological mechanisms beyond the pre-defined network structure. To validate the capability of BiOmics in an “omics-knowledge complementary” scenario within a unified latent embedding space (**Fig. 6a**), we applied the framework to single-cell transcriptome data from COVID-19 peripheral blood samples (6 healthy controls and 8 patients, including 4 ventilated individuals) to infer drug candidates driven jointly by data and knowledge[51] (**Fig. 6b**). Given that pharmacological effects vary across different cellular contexts, we conducted cell-type-specific drug repurposing analysis (**Fig. 6c**).

**Fig6.**
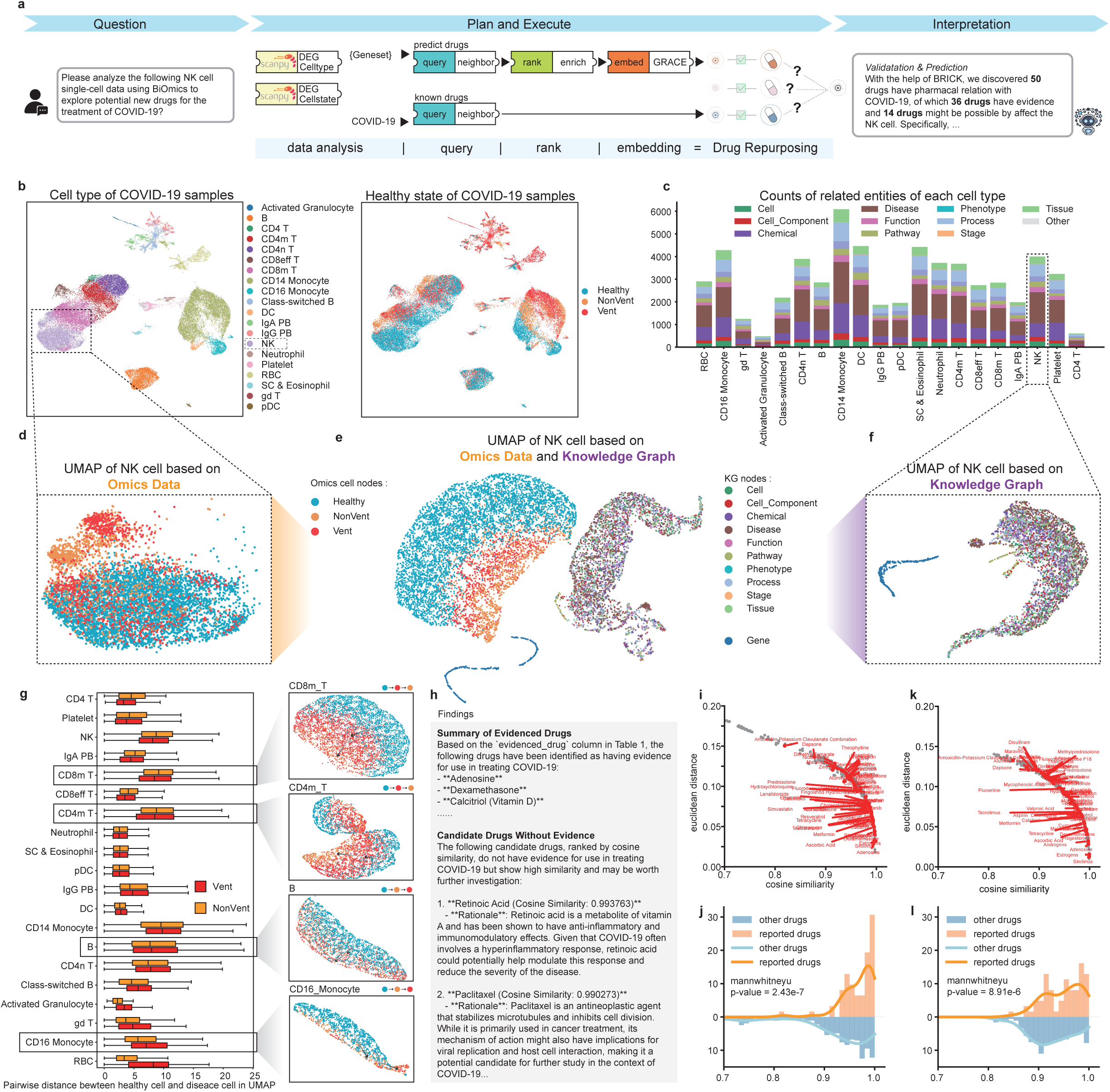
BiOmics catalyzing biomedical discovery. **a.** BiOmics pipeline for enhancing traditional omics analysis in case of drug repurposing. **b.** UMAP of COVID-19 PBMC dataset. left: colored by celltype, right: colored by health state. **c.** The stacked bar plot of queried entity type for each cell type. **d.** The UMAP of NK cell based on expression of Omics data. **e.** The UMAP of NK cell based on embedding of omics dataset and KG query result after representative learning. **f.** The UMAP based on embedding of KG query result after representative learning. **g.** The embedding of omics cell based on unite embedding of knowledge graph and omics data. left: Boxplot of embedding distances between Vent cells and healthy cells, as well as between NonVent cells and healthy cells for each celltype. right: examples of embedding for some of celltypes. **h.** BiOmics interpretation for drug repurposing task. **i.** The scatter plot of distance and similiarity between each chemical to COVID-19 based on knowledge graph and omics data. x-axis: cosine similiarity, y-axis: euclidean distance, reported COVID-19 chemicals are marked and labeled in red. **j.** The hist gram of cosine similarity of reported chemicals and other chemicals based on knowledge graph and omics data. the p-value of mannwhitneyu test equals 2.43e-7. **k.** The scatter plot of distance and similiarity between each chemical to COVID-19 based on only knowledge graph without omics data. x-axis: cosine similiarity, y-axis: euclidean distance, reported COVID-19 chemicals are marked and labeled in red. **l.** The hist gram of cosine similarity of reported chemicals and other chemicals based on only knowledge graph without omics data. the p-value of mannwhitneyu test equals 1.58e-6.

Taking NK cells as a narrative example, BiOmics integrated cell profiles, differentially expressed genes (DEGs) across patient states (Healthy, NonVent, Vent), and relevant biological entities from the knowledge graph into a unified representation learning space (**Supplementary Fig. 27**). The resulting embeddings effectively distinguished between various biological entities and cellular states (**Fig. 6d, e left, Supplementary Fig. 28**). Notably, most cell types (e.g., B cells, CD4n T cells, CD16 monocytes and CD14 monocytes) exhibited a linear Healthy→NonVent→Vent progression, reflecting standard disease advancement. However, NK cells displayed a distinct Healthy→Vent→NonVent transition. This non-linear pattern is likely attributable to NK cell exhaustion in ventilated patients, where reduced cellular vitality causes the omics signatures of Vent-state cells to cluster closer to those of healthy controls[52] (**Fig. 6g, Supplementary Fig. 29, 30, 31**).

The integrated embedding space of BiOmics not only captures the biological states of cells but also facilitates the convergence of functionally related biological entities. Following the injection of omics data, we ranked compounds associated with COVID-19 based on Euclidean distance and cosine similarity within the embedding space (**Fig. 6e right, f**). Our analysis demonstrated that compared to traditional link prediction on static KGs, established COVID-19 therapeutics such as Adenosine and Dexamethasone exhibited significantly higher cosine similarity and closer proximity to COVID-19 disease entities in the BiOmics space (**Fig. 6e, i, k**). The similarity distribution of reported drugs was highly concentrated within the 0.95-1.00 range, validating the framework’s ability to learn potential associations by embedding key genes, drugs, and phenotypes into a cohesive latent space (**Fig. 6e, j, l**). Furthermore, BiOmics identified several unreported compounds with high cosine similarity (≈0.98), such as Retinoic Acid and Paclitaxel, suggesting their potential as candidates for COVID-19 treatment through the modulation of NK cell activity (**Fig. 6h, Supplementary Fig. 34**). Moreover, by synergizing individual-specific omics signals with its extensive knowledge base, BiOmics provides a promising framework for personalized and precision drug recommendation, offering the potential to tailor therapeutic interventions to the unique molecular landscape of each patient.

### BiOmics demonstrates robust generalizability across multi-omics and proteomics landscapes

To further validate the extensive generalizability and scalability of the BiOmics framework, we applied BiOmics to a proteomics task (**Fig. 7a**). BiOmics analyzed quantitative proteomics of 10 placental malaria samples (5 healthy, 5 infected)[53]. Initially, BiOmics performed a systematic screening of differentially expressed proteins (**Fig. 7b**), then constructed a differential protein interaction network and marked phosphorylation and glycosylation modifications on the nodes (**Fig. 7c, d**). In the interaction network, node shapes represent protein modifications, and colors reflect the degree of difference between healthy and disease states. Starting from raw quantitative data, BiOmics retrieved a high-fidelity interaction network and successfully mapped distinct modification states. Functionalenrichment analysis of the differential proteins elucidated critical malaria-associated biological processes (**Fig. 7e**). For instance, *IGKC, ELANE, IGHG2* were related to Severe infection, among which *IGKC* and *IGHG2* were related to antigen binding processes. Notably, BiOmics identified *CLIC1, DSP, HSPB1, and VIM* as potential targets of *Artenimol*, a potent anti-malarial agent, providing significant pharmacological insights (**Fig. 7e**). Furthermore, the graph representation learning module of BiOmics was employed to predict novel protein-protein interactions. By generating joint embeddings, the framework transformed each protein into a high-dimensional vector representation (**Fig. 7f**). Through similarity calculations in the latent space, BiOmics screened for high-confidence interacting protein pairs (**Fig. 7g, h**). Among the Top 20 predicted interactions, 17 were successfully validated in the StringDB database[38], underscoring the system’s capacity for high-precision knowledge mining. These results collectively validate the superior generalizability and scalability of BiOmics in handling complex proteomics data, highlighting its utility as a rapid guidance tool for deciphering complicated disease mechanisms.

**Fig7.**
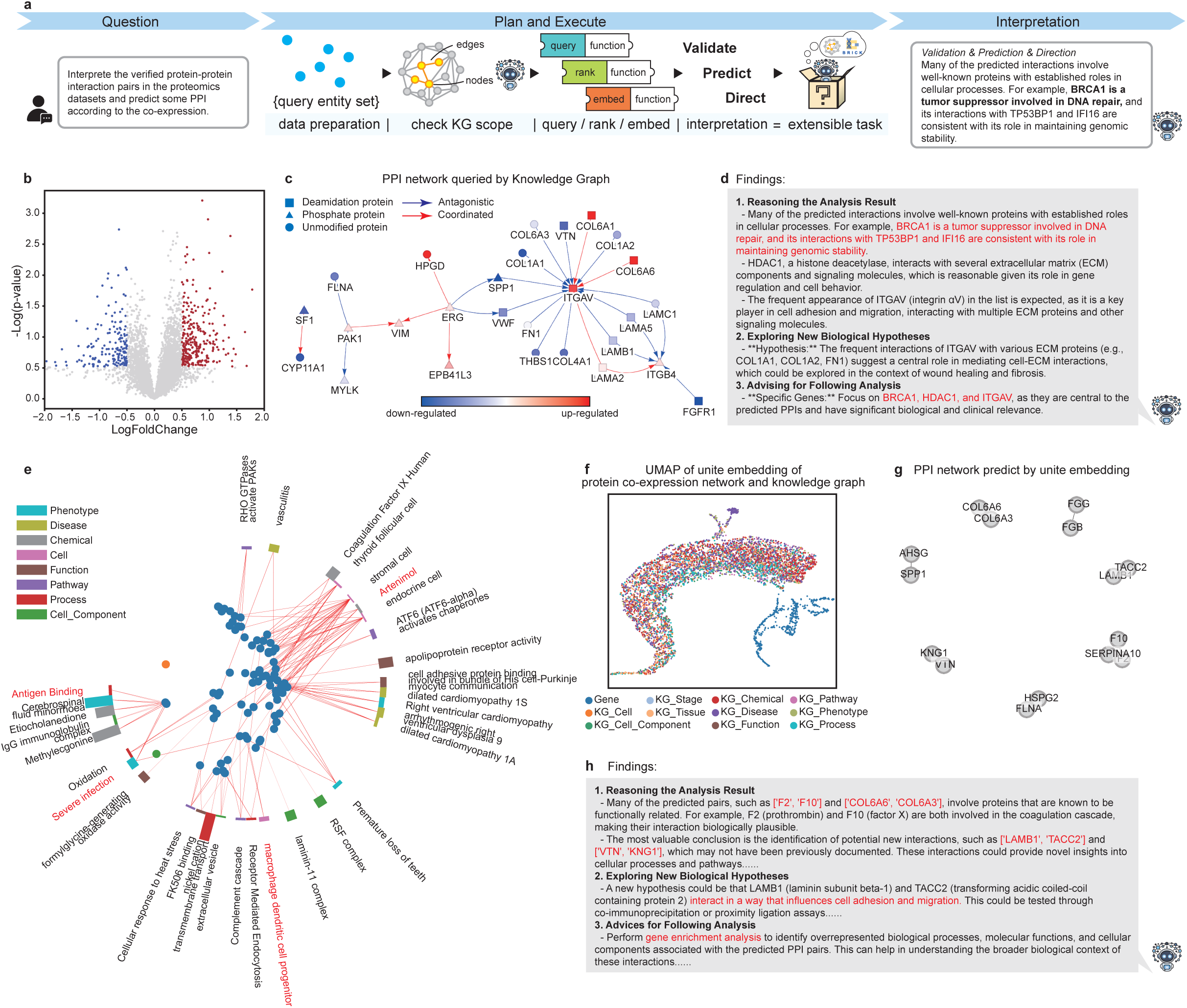
BiOmics facilitates multi-omics adaptability. **a.** BiOmics pipeline for extensible application on other omics analysis. **b.** Volcano plot of differential expression protein of infected samples. x-axis: log-fold-change, y-axis: −log(p-value), blue and red marked positive markers and negative markers, respectively. **c.** Visualization of protein-protein interaction(PPI) network. Points in the shape of circles, triangles, and squares represent normal proteins, phosphorylated proteins, and glycosylated proteins respectively. Red and blue color represent log-fold-change of differetial expressed proteins. **d.** The interpretation context about PPI task based on BiOmics queried result. **e.** Radial enrich plot of marker of differential expression protein. The colors of the bars distinguish the types of enriched entities, and the heights of the bars represent the number of genes in the enriched entities. f. The UMAP based on unite embedding of proteomics dataset and KG query result after representative learning. **g.** The predicted PPI network based on unite embedding of proteomics dataset. **h.** The interpretation context about PPI task based on BiOmics unite embedding result.

## Discussion

In the evolving AI4S landscape, research has predominantly focused on deploying autonomous agents for automated bioinformatics execution. However, a critical gap persists: the translation of high-confidence analytical outputs into interpretable and verifiable biological discoveries. BiOmics addresses this challenge through a tripartite “Retrieving-Reasoning-Predicting” paradigm, characterized by hallucination-free retrieval of prior knowledge, long-chain traceable causal reasoning, and adaptive data-knowledge fusion. This framework generates foundational entity representations that accelerate the data-to-discovery translation. To establish an engineering-level closed loop, BiOmics-KG (a daily-updated knowledge graph) standardizes schemas to mitigate LLM hallucinations, while BiOmics-BRICK (a multi-omics and knowledge graph integration toolkit) maps heterogeneous data into a shared inference space. Finally, BiOmics-Agent orchestrates these components into a streamlined pipeline, effectively bridging the divide between raw analysis and mechanistic discovery. Real-world multi-omics validation confirms its capacity to enhance knowledge mining depth and hypothesis quality, serving as an out-of-the-box foundation for data-driven biological research.

The core value of the BiOmics architecture resides in its seamless synthesis of analysis and interpretation, effectively collapsing the traditional divide between computational processing and biological reasoning. Guided by robust biological knowledge, analysis transcends simple data-pattern dependency to generate credible biological outcomes, while associative insights beyond established boundaries enable the mining of novel mechanisms. This capability relies on two pivotal designs. First, the harmonized explicit reasoning space provides a logical framework for the “input–fusion–conclusion” pipeline, inferring biologically consistent results while supporting retrospective reasoning-chain validation to rectify contradictions with existing knowledge. Second, the unified latent embedding space fuses multi-omics data with precisely retrieved knowledge entities to eliminate modality barriers. This facilitates customized data-driven fusion and enables biological knowledge to inform entity correlation analysis, aligning feature associations with both statistical rigor and biological significance.

BiOmics’ innovative application drives a transformative research paradigm. Its built-in agents automate end-to-end workflows for standard omics tasks (e.g., association screening, mechanism validation, causal reasoning), freeing researchers from repetitive work to focus on refining core biological questions, This automated paradigm addresses a critical AI4S bottleneck: while current tools are often confined to isolated links like data collection or model building, BiOmics’ fusion architecture bridges the entire intelligent research chain.

As specialized multi-agent systems (e.g., SRAgent[54] for data collection, CellAgent[9] for analysis) permeate the omics ecosystem, BiOmics complements rather than competes with them by filling the unique “result to interpretation” gap. We propose a collaborative vision: encapsulating these tools as pluggable components within a federated network that covers the full “collection-analysis-interpretation” loop. Such a network could mine the “dark matter” in historical sequencing data while expanding global knowledge repositories. BiOmics provides RESTful APIs, MCP Servers, and Web interfaces for standardized integration; its knowledge-driven engine enables cross-modal, long-chain reasoning, advancing bioinformatics mining from simple correlation to deep mechanism.

Limitations and Future Directions: While multi-omics experiments confirm BiOmics’ cross-omics transferability, certain gaps remain. The framework currently lacks systematic evaluations for specialized fields such as metabolomics and epigenomics. As a cell-omics-focused platform, its broader generalization requires collaboration with domain experts to expand KG coverage and accumulate diverse cross-omics cases. Furthermore, data processing currently relies on LLM-generated code (e.g., Scanpy[55], Stereopy[56]), which, while effective for standard tasks, may struggle in highly complex scenarios. This necessitates optimization through more specialized analytical agents. Future upgrades will focus on developing an “active research assistant” that autonomously identifies knowledge gaps to provide proactive support, thereby fully complementing the burgeoning omics multi-agent ecosystem.

## Methods

### Logical framework and paradigms of BiOmics

BiOmics serves as a foundational agent designed to bridge the gap between high-dimensional omics data and biological insight. It offers a “plug-and-play” capability for complex interpretation tasks, executed through three core logical paradigms: Knowledge Retrieval, Causal Reasoning, and Association Prediction. This framework encompasses the complete cognitive continuum, from “accessing established facts” and “inferring implicit mechanisms” to “forecasting latent associations”, forming a graph-driven closed loop that unifies data analysis with biological interpretation. Upon receiving natural language queries and multi-omics files, BiOmics functions as a logical orchestrator. It performs intent recognition to categorize requirements and subsequently decomposes tasks into modular execution workflows, dynamically invoking the following specialized logic:

#### (1) Knowledge retrieval logic (accessing established facts)

Objective: To precisely extract structured relationships from the Knowledge Graph (KG) to provide factual grounding for biological queries.

##### Step 1: Task parsing and parameterization

The agent extracts query parameters from natural language inputs:

i. Entity alignment: Identifies source and target entities by mapping natural language descriptors to canonical Knowledge Graph (KG) names and types.
ii. Relational constraints: Determines if specific biological edge types (e.g., regulation, co-expression, co-mention) are required to filter the search.
iii. Topological depth: Sets the number of query hops (default 1–3) based on the search scope or user-defined requirements.

##### Step 2: Automated query generation & execution

The agent populates specialized Cypher templates with the extracted parameters and executes the query to perform targeted interrogation of the knowledge graph.

##### Step 3: Knowledge synthesis and output representation

The results are compiled into a structured evidence suite, providing relational path tables, NetworkX-compatible subgraphs[57], and interactive visualizations.

#### (2) Reasoning logic (inferring implicit mechanisms)

Objective: To infer high-order biological entities (e.g., cell identities, pathological phenotypes) associated with omics profiles by navigating the explicit reasoning space.

##### Step 1: Task parsing and path mapping

i. Reasoning trajectory identification: Maps the logical path from input data objects to potential biological targets (e.g., Cluster → Differential Genes → Cell Identity).
ii. Dependency assessment: Determines if computational preprocessing (e.g., differential expression analysis) is required to prime the reasoning chain.

##### Step 2: Autonomous data processing

Evaluates task-specific requirements and invokes the agent to generate and execute bioinformatic code (via the BRICK toolchain) to extract core biological signals.

##### Step 3: Explicit reasoning space construction

i. Entity extraction: Identifies high-confidence biological entities from omics data.
ii. Bridge entity extraction: Extracts key omics features and identifies “bridge” entity categories to link data with the Knowledge Graph (KG).
iii. Knowledge grounding: Retrieves contextually relevant subgraphs from the KG via 1–3 hop neighborhood expansion.
iv. Sub-graph construction: Constructs a unified graph by bridging the omics-derived observation-feature sub-graph and adaptive knowledge sub-graph based on BiOmics-KG connected by features.
v. Reasoning space formulation: Establishes the directional reasoning trajectory based on valuable information from unified graph for LLM context.

##### Step 4: Causal path inference

Constructs logical reasoning chains within the harmonized reasoning space, employing ranking and pruning algorithms to isolate high-confidence biological connections from data to targets.

##### (i) Step 5: Mechanistic interpretation and reporting

The LLM synthesizes the inferred paths into a tripartite evidentiary report, including concise overview of identified associations, mechanistic insights derived from the reasoning chain and actionable suggestions for downstream validation.

#### (3) Prediction logic (forecasting latent associations)

Objective: To leverage the unified latent embedding space and graph representation learning to forecast unknown associations where direct KG evidence is absent.

##### Step 1: Task parsing and trajectory mapping

i. Prediction trajectory identification: Defines the logical path from input omics data to potential biological targets (e.g., Cell State → Key Genes → Therapeutic Candidates).
ii. Dependency assessment: Identifies necessary computational preprocessing (e.g., differential expression analysis) to prepare the data for feature extraction.

##### Step 2: Autonomous data processing

Evaluates task requirements and invokes the agent to generate and execute bioinformatic code (via the BRICK toolchain), extracting core biological features from the datasets.

##### Step 3: Unified feature space construction

(ii) Entity extraction: Identifies high-confidence biological entities from the processed omics results.
(iii) Bridge entity extraction: Extracts key omics features and identifies “bridge” entity categories to link data with the Knowledge Graph (KG).
(iv) Knowledge grounding: Retrieves contextually relevant subgraphs from the KG via 1–3 hop neighborhood expansion.
(v) Sub-graph construction: Constructs a unified graph by bridging the omics-derived observation-feature sub-graph and adaptive knowledge sub-graph based on BiOmics-KG connected by features.
(vi) Embedding space mapping : Projects the unified sub-graph into a unified embedding space using graph representation learning (e.g., AnyGraph[58] or GRACE[59]) to synthesize high-dimensional node features that bridge omics data and biological knowledge.

##### Step 4: Latent association quantification

Calculates relational proximity scores (e.g., cosine similarity) within the latent embedding space to prioritize the most probable associations between candidate and query entities.

##### Step 5: Mechanistic interpretation and reporting

The LLM synthesizes the inferred paths into a tripartite evidentiary report, including concise overview of identified associations, mechanistic insights derived from the reasoning chain and actionable suggestions for downstream validation.

### BiOmics-KG construction

BiOmics-KG serves as the foundational knowledge engine for the BiOmics platform, integrating heterogeneous biological data into a structured, scalable, and temporally sensitive framework. Through unified schema design, rigorous data normalization, and an automated update pipeline, BiOmics-KG facilitates high-fidelity knowledge discovery and intelligent reasoning.

#### (1) BiOmics-KG schema structure

##### Nodes

BiOmics-KG employs a hierarchical schema comprising six primary and 13 secondary classes, including anatomical concepts (Species, Tissue, Cell, Cell Component); temporal concepts (Stage); molecular entities (Gene, Protein, Mutation, Chemical); biological processes / functions (Pathway, Function, Process); and bibliographic evidence (Publication) (**Supplementary Fig. 3**). Each node is characterized by a standardized attribute set, including Node Type, Database ID, Name, Definition, and Synonyms.

##### Relationships

We adopted the Relation Ontology (RO)[60] as our gold standard, Curating 28 fundamental biological relations. To maintain provenance and granularity, we implemented *original_relation* to preserve source-specific descriptions and *info_source* to document evidence origins (**Supplementary Table 2**).

#### (2) Data integration and automated normalization

We collected data from ontologies, curated databases, and literatures.

##### Integration of curated ontologies and Databases

Heterogeneous data from source ontologies and databases were converted to align with our predefined schema. Core attributes—including ID, name, definition, and synonyms—were directly mapped from primary sources, with missing fields designated as null. Given that node-type categorization is pivotal for Knowledge Graph (KG) retrieval performance, we developed an *LLM-based taxonomic agent* to classify entities lacking categorical metadata. Benchmarking against the MeSH database demonstrated that this agent achieved a taxonomic classification accuracy of 98.3%. Subsequently, relational data were manually curated and mapped onto the 28 standardized relationship classes within our schema. Comprehensive details regarding data sources and processing protocols are provided in Supplementary Table 2.

##### Literature mining

Real-time knowledge acquisition from primary literature is managed by the *AutoPubMedKG builder*, a specialized agent for automated extraction and graph adaptation (see Section (5).

#### (3) Graph refinement and deduplication

To enhance graph quality and retrieval efficiency, we performed a two-stage refinement process:

##### Node consolidation

We designated primary databases for each entity class (**Supplementary Table 3**). Entities from secondary sources were merged into primary nodes through naming overlap analysis and cross-reference validation (via BioPortal[16]). Post-merging, all relational links were re-routed to the primary entity, and redundant metadata were consolidated.

##### Relation deduplication

Identical triplets (Head, Relation, Tail) were merged into unique edges. Crucially, the length of the *info_source* list is utilized as a confidence metric (*info_source_length*), providing a quantitative weight for downstream reasoning tasks within the BRICK toolchain.

#### (4) Automatic update of literature knowledge

To ensure coverage and timeliness, we designed an automated update pipeline encompassing literature acquisition, entity recognition, and graph updating.

##### Daily literature acquisition

The system automatically retrieves the latest publications via the PubMed API.

##### Hybrid entity recognition

We utilized BERN2[61] for standard entities (genes, diseases). To overcome the limitations of pre-trained models for specialized categories (e.g., phenotypes, pathways), we implemented a dictionary-driven approach using the Aho-Corasick suffix tree algorithm, based on comprehensive ontological vocabularies (Supplementary Table 4).

##### Traceable relationship extraction

To address the inherent complexities of open-domain relation extraction, we implemented a sentence-level co-occurrence paradigm. When two entities co-occur within a single sentence, a “co-mentioned” relationship is instantiated. Crucially, the source sentence is preserved as a provenance attribute within the *original_relation* field, ensuring the traceability and mechanistic interpretability of all inferred biological links.

##### Literature node construction

Each PubMed article was constructed as a Publication node and connected to the mentioned entities through edges. Meanwhile, *cited_by* edges were constructed based on citation relationships to form a literature-based sub-network.

##### Systematic automated update

The complete workflow, from data retrieval to graph integration, is executed as a nightly cron-job. This facilitates incremental daily updates, ensuring the BiOmics-KG remains a temporally sensitive and current resource for multi-omics discovery.

#### (5) Implementation and infrastructure

The KG was built on Neo4j[62] (v5.14.0 Community Edition). The cleaned ontology and public knowledge base data were imported into the knowledge graph, where node-related information was stored as node attributes and relationships as edges.

### BiOmics-BRICK implementation

BiOmics-BRICK serves as the operational orchestration layer of the framework, providing a standardized suite of functional modules that bridge the gap between raw bioinformatic processing and high-order biological reasoning.

#### (1) Multi-modal data processing engine (data processing module)

This module facilitates the seamless transition between tabular omics data and graph-structured topologies:

##### Autonomous bioinformatic execution

Leverages LLM-driven code generation to perform standardized omics workflows, including preprocessing, clustering, and trajectory inference, primarily utilizing the Scanpy[55] ecosystem.

##### Graph-relational transformation

Standardizes the conversion of single-cell objects (AnnData) into bipartite/KNN graph structures and transforms analytical results (DataFrames) into relational triplets (e.g., Gene-Pathway-Disease).

##### Unified graph integration

Orchestrates the fusion of omics-derived sub-graphs with adaptive knowledge sub-graph derived from BiOmics-KG to construct a unified substrate for downstream representation learning. For omics-derived sub-graphs, the edges between observation and feature (e.g. cell and gene) established from quantitative matrix (e.g. expression matrix), meanwhile the edges among observations established based on K-Nearest Neighbor (KNN) similarity. The adaptive knowledge sub-graph is constructed according to the queried and ranked result based on key feature from omics data.

#### (2) High-performance knowledge interrogation (query module)

Built upon a Neo4j backbone, this module translates natural language intents into optimized Cypher queries:

##### Standardized interfacing

Architected a universal query interface that returns multi-modal formats (e.g., Pandas DataFrames for statistics, NetworkX[57] MultiDiGraphs for topology).

##### Parametric templates

Provides a library of high-frequency templates for complex biological exploration, including multi-hop neighborhood expansion, shortest-path identification, and relational sub-network extraction between disparate entity sets.

#### (3) Multi-dimensional evidence prioritization (ranking module)

To distill actionable insights from dense KG retrievals, BRICK employs a prioritization engine featuring six statistical strategies: support-source count, database-density scores, matching frequency probability, enrichment significance (hypergeometric tests), and multi-index ensemble voting. These strategies enhance the biological relevance and interpretability of inferred associations.

#### (4) Mechanistic reasoning engine (reasoning module)

This module operates within the reasoning space, integrating filtered prior knowledge with data-driven features to synthesize biological findings. It follows a tripartite interpretive paradigm:

##### Evidence Validation

Cross-referencing analytical results against established biological laws.

##### Hypothesis Generation

Identifying discrepancies between data and prior knowledge to propose novel mechanisms.

##### Strategic Guidance

Heuristically planning subsequent analytical tasks based on inferred biological context.

#### (5) Latent representation learning (Embedding Module)

To achieve deep predictive interpretation, BRICK facilitates the mapping of biological entities into the embedding space. It supports a unified interface for state-of-the-art graph models: (i) GRACE: A graph contrastive learning framework for robust node representation[59, 63]. (ii) AnyGraph: A foundational graph model capable of zero-shot cross-domain inference, enabling tasks such as reference-free cell annotation and drug repurposing without the need for task-specific re-training[58].

#### (6) Semantic visualization and interactivity (Visualization module)

BRICK provides an expansive suite of visualization tools to enhance interpretability, ranging from interactive relational graphs[64] and TF-target regulatory networks to knowledge-annotated volcano plots and hierarchical functional enrichment maps[65].

Through unified interface design, BiOmics-BRICK highly modularizes data processing, query, ranking, reasoning, representive learning, and visualization functions, supporting a complete closed loop from original query to biological interpretation. Its flexibility and extensibility enable seamless integration into various omics scenarios such as single-cell analysis, drug discovery, and causal SNP identification, providing a powerful and intuitive graph-driven toolchain for biological research.

### BiOmics-Agent architecture

BiOmics-Agent functions as the central intelligence of the framework, responsible for autonomous workflow orchestration and high-order interpretation of multi-omics data. By synergizing multi-agent collaboration with a specialized long-short-term memory (LSTM) mechanism[32], the Agent transforms natural language queries into traceable, high-value biological insights through a collaborative, reasoning-centric paradigm (**Supplementary Fig. 1**).

#### (1) Hierarchical multi-agent orchestration

The multi-agent system comprises 10 sub-agent systems (**Supplementary Table 5, Supplementary Fig. 1c**). Among them, *env_checker* first parses the complete information of the uploaded data, which is then summarized by *data_analyzer* and stored in short-term memory. Subsequently, *analyze_planner* decomposes the overall task into multiple subtasks based on data characteristics, user needs, successfully executed cases, and other memory-related information. These subtasks are then forwarded to *planner*, which retrieves relevant BRICK package source code via Retrieval-Augmented Generation (RAG)[30]. A comprehensive subtask list which including subtask serial numbers, content, BRICK functions utilized, and function source code, is encapsulated into short-term memory. *plan_executor* provides uncompleted subtasks from the subtask list to a sub-agent group (*coder, code_debugger, code_runner*) designed based on the ReACT paradigm[66] for collaborative completion. Specifically, *coder* first generates code based on subtask information, data content, BRICK source code retrieved by planner, and successfully run cases stored in long-term memory. *code_runner* executes the generated code and checks for successful operation. If the code runs successfully, indicating the completion of the current subtask, *plan_executor* is invoked to obtain a new subtask. If an error occurs, *code_debugger* is called to diagnose the error cause based on error information and detailed data, generate revised code, and then invoke *code_runner* for re-execution. Upon completion of all subtasks, the system saves all intermediate graph files and analytical results. Responder then interprets the final analytical results using a set of reusable interpretation prompts. Additionally, if the user’s query is unrelated to bioinformatics tasks, the aforementioned workflow is not employed; instead, *general_responder* is directly activated to generate general responses leveraging the capabilities of LLMs.

#### (2) Human-in-the-loop collaborative intelligence

BiOmics-Agent implements a sophisticated human-machine interaction mechanism powered by LangGraph[67]. This ensures that the discovery process is not a “black box” but an intervenable workflow. Users are guided through three critical synchronization checkpoints: Data Inspection, Initial Report Generation, and Analysis Plan Formulation. Feedback at these stages is dynamically integrated into the Short-term memory to re-align the workflow, ensuring the final results are strictly concordant with the user’s research objectives (**Supplementary Fig. 1a**).

#### (3) Synergistic long-short-term memory mechanism

The agent’s capacity for “thinking” and “interpreting” is anchored by its unique dual-memory architecture:

##### Long-term memory

Serves as a permanent experiential repository, housing the vectorized BiOmics-KG, historical successful analysis cases, and the standardized BiOmics-BRICK toolchain. LTM provides the domain-specific expertise required for high-fidelity code generation and reasoning (**Supplementary Fig. 2**).

##### Short-term memory

Functions as a dynamic task context, dedicated to storing current data features, user preferences, and intermediate analytical states. This synergy allows specialized agents (e.g., planner, coder, and responder) to combine foundational biological laws with task-specific nuances, ensuring both the robustness and flexibility of the interpretation.

Through the convergence of multi-agent orchestration, collaborative interaction, and memory-driven reasoning, BiOmics-Agent transcends traditional automated pipelines. It functions as an autonomous, reasoning-centric scientific colleague, one that is not only automated but also interpretable, iterable, and capable of evolving through scientific dialogue.

### Benchmarking and evaluation

We performed an extensive benchmarking of BiOmics across eleven distinct biological categories to evaluate its performance in knowledge retrieval, reasoning-driven analysis, and association prediction. BiOmics was compared against LLMs (GPT-4o)[68], biological agents (Biomni)[10], and specialized bioinformatic toolkits.

#### QA task

For the QA task, we assessed BiOmics in terms of QA accuracy and answer richness. Regarding accuracy, we used the multiple-choice and true/false question datasets from Biomix to evaluate the answer accuracy of BiOmics, Biomni, and GPT-4o. Additionally, we further evaluated BiOmics’ accuracy in cases where information was retrievable. For the assessment of answer richness, we constructed a dataset consisting of 25 examples each for cells, diseases, genes, and mutations, with the query “What is XXX?”. We then counted the number of entities mentioned in the answers from BiOmics, Biomni, and GPT-4o respectively.

#### Cell type annotation

Leveraging 17 heterogeneous single-cell datasets from the CellxGene database[34], we assessed the annotation fidelity of BiOmics across diverse tissue types. Comparative baselines included GPTcelltype (driven by GPT-4o), Cellmarker2 (database-centric), and Biomni[10, 45, 46].

#### Drug repurposing

We performed drug repurposing analysis across various cell types using human COVID-19 peripheral blood datasets[51]. Using therapeutics documented in ClinicalTrials as the gold standard, we calculated the Recall@20 of reported compounds for each cell-type-specific case. For BiOmics, the top 20 candidates were extracted from the autonomous reasoning discourse; for the link-prediction baseline, we quantified latent proximity by calculating the cosine similarity between jointly embedded drug and COVID-19 disease nodes. For GPT-4o, a list of retrievable compounds was provided, and the model was tasked with prioritizing the most promising therapeutic leads.

#### Phenotype prediction

Using 637 mutation profiles (dbSNP IDs) from TCGA, we evaluated the capacity of BiOmics, Biomni, and GPT-4o to infer disease phenotypes. Evaluation was conducted at two granularities: stringent match (identical to the ground truth) and semantic match (phenotypes occurring within the same physiological system or oncological domain). All models received mutation lists as input to predict associated clinical phenotypes.

#### Causal SNPs identification

Utilizing a colon cancer case from TCGA (case_id: d34894bf-a0bf-468b-9b99-e821d781409f), we measured the precision of each method in prioritizing pathogenic variants, cross-referencing results with the ClinVAR database. For BiOmics, Biomni, and GPT-4o, all variants identified in the final interpretations were extracted for evaluation; for PolyPhen[42], we applied a standard pathogenicity threshold of 0.9.

#### Trajectory inference

Based on mouse embryo white blood cell (WBC) data[48], we evaluated the topological consistency of developmental lineages against expert-curated developmental relationships. BiOmics and Biomni were provided with raw single-cell transcriptomic data, while GPT-4o was provided with the full list of constituent cell types. The PAGA algorithm (within the Scanpy ecosystem) served as the traditional bioinformatic baseline, utilizing a connectivity threshold of 0.05.

#### Gene enrichment analysis

Using pancreatic PP cells from a mouse islet dataset[44], we quantified the volume of biologically significant entities enriched by each method. BiOmics, Biomni, and clusterProfiler[69] directly ingested differential gene lists, whereas GPT-4o was queried to extract the most relevant biological concepts based on the same gene lists.

#### Protein-protein interaction (PPI) analysis

We assessed the precision of inferred PPIs using human placenta mass spectrometry data[53], with StringDB[38] serving as the ground truth. BiOmics and Biomni processed the raw proteomics dataset; for GPT-4o, the differential protein list was provided as a textual summary; and for CellphoneDB[70], the identified protein pairs among differential features were directly quantified.

#### Gene regulatory network (GRN) analysis

Focusing on the *Pparg* transcription factor in mouse embryo WBC data[48], we calculated the precision of predicted regulatory pairs against the TRRUST database[37]. BiOmics processed the target gene list; Biomni and GPT-4o were queried directly for *Pparg* targets; and for SCENIC[50], regulatory associations were quantified based on the raw expression matrix.

#### Spatial analysis

Using a mouse testis spatial transcriptomics dataset[71], we evaluated each model’s ability to infer anatomical spatial relationships between cell types. Performance was measured via the F1-score relative to expert-curated spatial annotations. BiOmics and Biomni were provided with the spatial dataset, while GPT-4o was provided with the cell type list. The co-occurrence algorithm in Stereopy[56] (threshold=0.7) served as the quantitative baseline.

#### Differential analysis

We evaluated the biological significance of marker genes identified by each method for pancreatic PP cells[44]. Performance was quantified by the average log-fold change (logFC) of the identified marker genes within the dataset. BiOmics and Biomni processed the raw transcriptomic data, while GPT-4o was directly queried for known PP cell markers. The rank_genes_groups function in Scanpy (p < 0.05, score > 0) was utilized as the standard bioinformatic baseline.

### Data preprocessing and analysis

#### Data preprocessing

For mouse leukocyte single-cell data used in trajectory inference[48], we first performed PAGA trajectory inference, dividing the dataset into three lineages using a threshold of 0.05 (**Supplementary Fig. 18**); subsequent analysis focused on the myeloid cell group. For human placental malaria proteomics data[53], we converted it to the AnnData structure, where obs corresponds to 10 samples and var corresponds to protein names.

#### pySCENIC analysis

For the mouse embryo WBC dataset[48], pySCENIC was executed prior to BiOmics interpretation, allowing the agent to directly access regulatory inference results.

#### Trajectory validation

To validate the stronger association of the Monocyte-to-Adipose Macrophage trajectory (vs. cDC-to-Adipose Macrophage) in the unified embedding, we calculated the Jaccard similarity and Pearson correlation of TF activities along these lineages.

#### Drug similarity significance

For COVID-19 drug repurposing[51], a Mann-Whitney U test was performed to verify that known COVID-19 drugs exhibited significantly higher cosine similarity to the COVID-19 node in the unified embedding space compared to unreported drugs.

#### Differential expressed protein

For the human placental malaria proteomics data[53], after we converted it to the AnnData structure, we ask BiOmics to preform a differential expressed gene using scanpy with an exploratory threshold (|logFC|>0.5 and p<0.3).

## Data availability

The omics data integrated into BiOmics is categorized into two distinct subsets: evaluation data for benchmark(**Supplementary Table 6**) and analysis case data for case study (**Supplementary Table 7**). For the evaluation subset: Biomix mcq and tf datasets are retrieved from the Hugging Face platform (BiomixQA, https://huggingface.co/datasets/kg-rag/BiomixQA)[33]; an in-house dataset is employed for the QA abundance task (detailed in **Supplementary Table 8**); 17 tissue-specific cellxgene datasets are utilized for cell annotation (**Supplementary Table 9**)[34]; the GSE150728 dataset supports drug repurposing analyses[51]; 639 open-access TCGA cases with corresponding identifiers are used for phenotype prediction; the All of Us 2024 T2D WG dataset is adopted for causal SNP identification[35]; the GSM2230761 dataset is applied for differential expression gene and gene enrichment analyses[44]; the GSE228590 dataset facilitates trajectory inference and gene regulatory network (GRN) investigations[48]; the PRJNA668433 dataset is used for spatial analysis[71]; and the PXD008079 dataset underpins protein-protein interaction studies[53].

The analysis case subset including: the All of Us 2024 T2D WGS dataset (human/Type 2 diabetes); the GSM2230761 dataset (mouse/pancreatic islets); the GSE228590 dataset (mouse/embryo WBC); the GSE150728 dataset (human/PBMC); the PRJNA668433 dataset (mouse/testis); and the PXD008079 dataset (human/placentas)[35, 44, 48, 51, 53, 71].

## Code availability

The code used to develop the model, conduct the analyses, and generate the results reported in this study is publicly available and deposited in the GitHub repository.

## Acknowledgments

This work is supported by National Natural Science Foundation of China (32300526).

## Author contributions

S.S.F. and L.C. conceived the study and proposed the primary conceptual framework. L.C., H.Q., Y.T.L., and Y.B.S. were responsible for algorithm development and implementation. Data analysis was performed by L.C. and Y.B.S. C.L. and Y.T.L. contributed to the implementation of the agent. L.C., W.Y.D., B.J., and L.D. constructed the knowledge graph. Y.L.Z. was responsible for figure visualization and typesetting. T.Y.X., L.N.H., and H.Y.H. provided key insights and technical advice. The study was supervised by S.S.F., Y.Z., and Y.X.L.

