## Supplemental Figures for "BiOmics: A Foundational Agent for Grounded and Autonomous Multi-omics Interpretation"

FigureS1

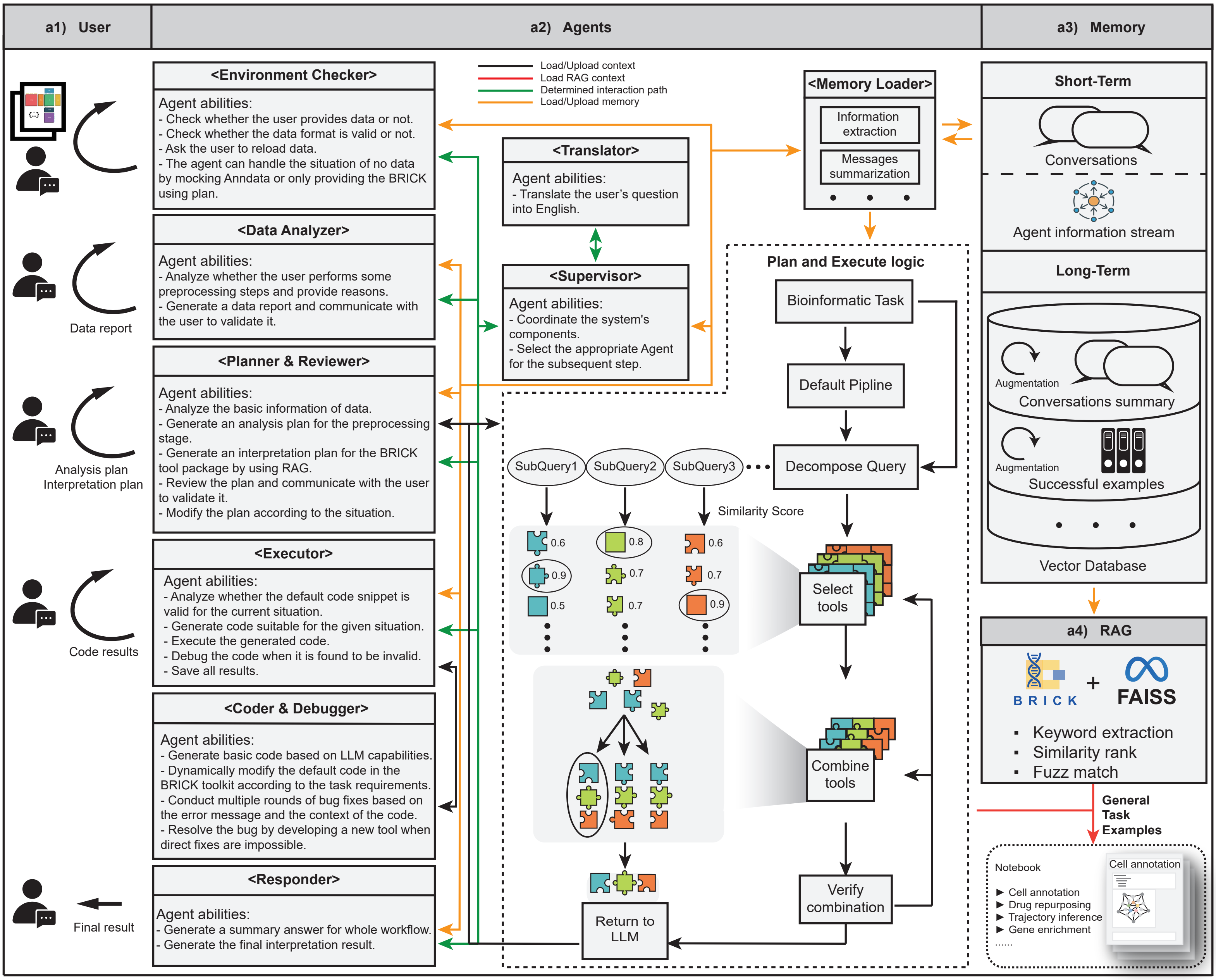


**Supplementary Fig S1. BiOmics Agent schema. a1.** The user interface. Users can submit their omics data along with specific requirements to the BRICK agent system and engage in interaction with the designated agent at every critical stage of the system's operation. Furthermore, users are also able to retrieve the intermediate operational results from each step of the process. **a2.** The agents module. This module illustrates the interaction flow and the agent's capabilities. The black line indicates the logic for loading or uploading context, the green line represents the determined interaction pathway logic, the green dotted line denotes the logic for loading RAG context, and the yellow line signifies the logic for loading or uploading memory context. **a3.** The memory module. BRICK's memory module comprises both short-term and long-term memory components. When BRICK stores its memory, it summarizes the chat history and contextual messages from each agent. When BRICK retrieves its memory, it searches and extracts relevant information from this memory module. **a4.** The RAG module. BRICK's agent through the RAG module to access the initial BRICK tools and follow the given instructions. outlines the various steps involved in retrieving the appropriate tool and generating the processing pipeline.

FigureS2

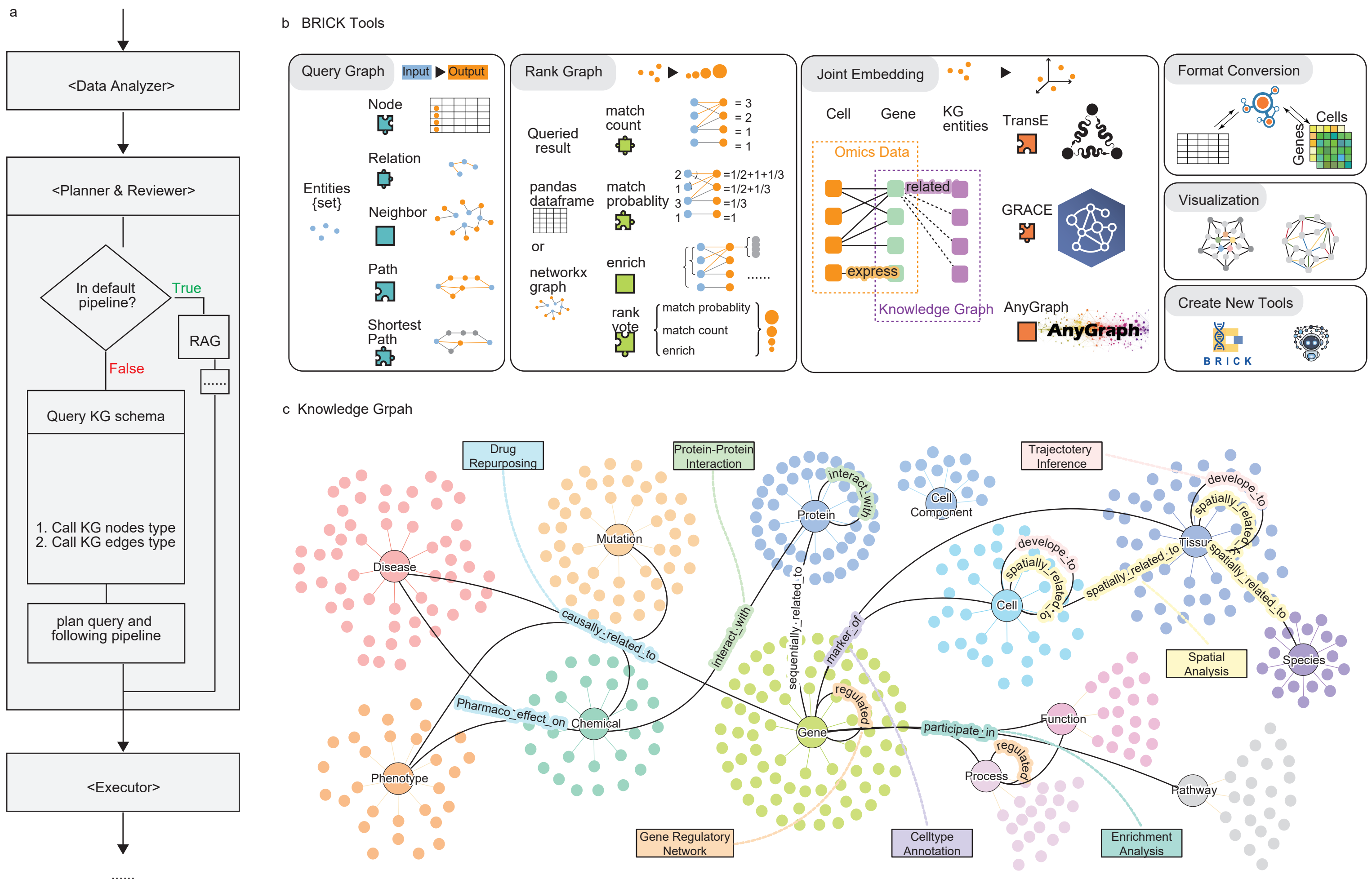


**Supplementary Fig S2. Diagram of how BiOmics agent interact with BiOmics-BRICK and BiOmics-KG. a.** BiOmics Agent call knowledge graph schema when meeting tasks on other omics. **b.** The BRICK tools. BRICK tools contains following modules: query graph to get related knowledge according to omics data analysis result; rank graph to filter and sort valuable knowledge; unite embedding module embed knowledge and omics data in a unite space; format conversion make sure the all modular function can connected with each other. visualization module highlight knowledge in analysis result. **c.** The schematic diagram of the knowledge graph data. The edge color of each relationship corresponds to the fill color of its associated bioinformatics task, maintaining visual consistency.

FigureS3

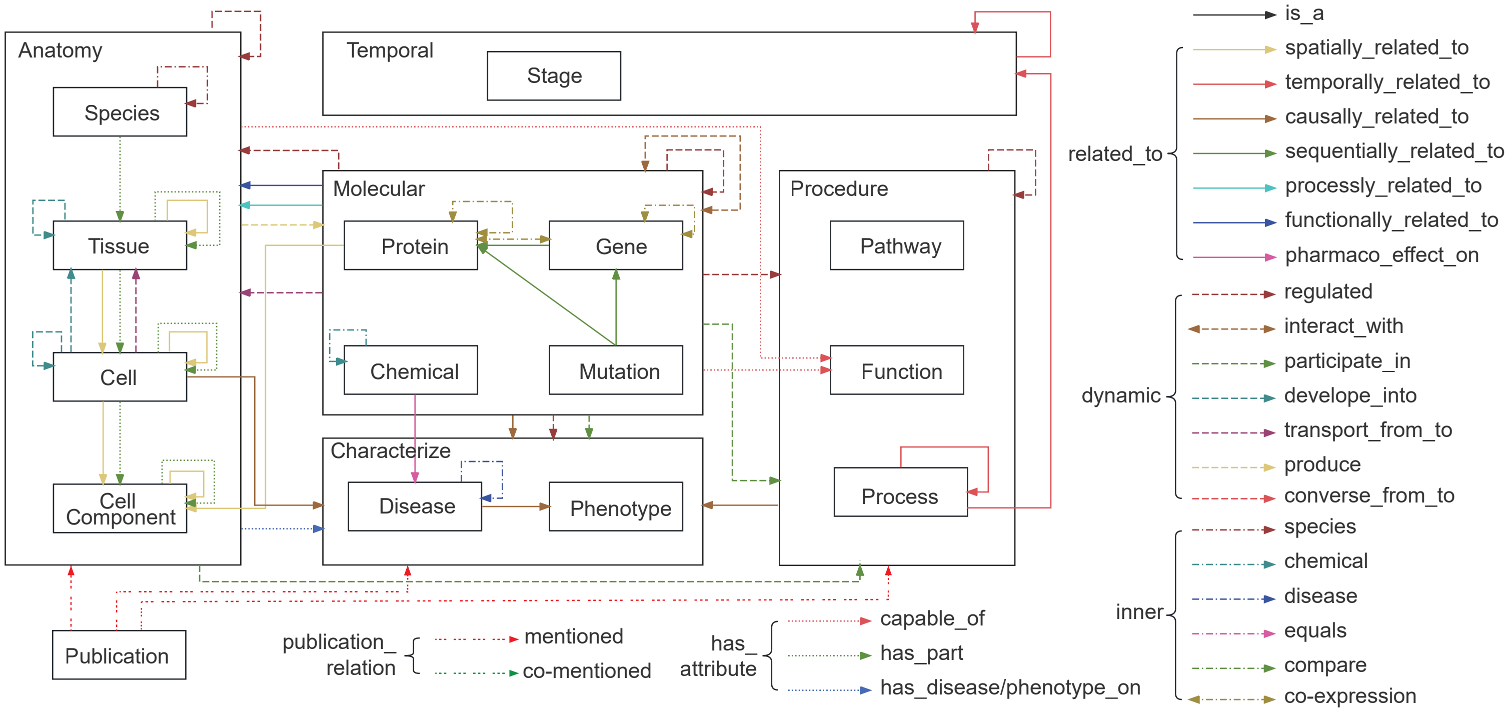


**Supplementary Fig S3. BRICK knowledge graph schema. Box represent entity type and lines represent the different relationship types.**

FigureS4

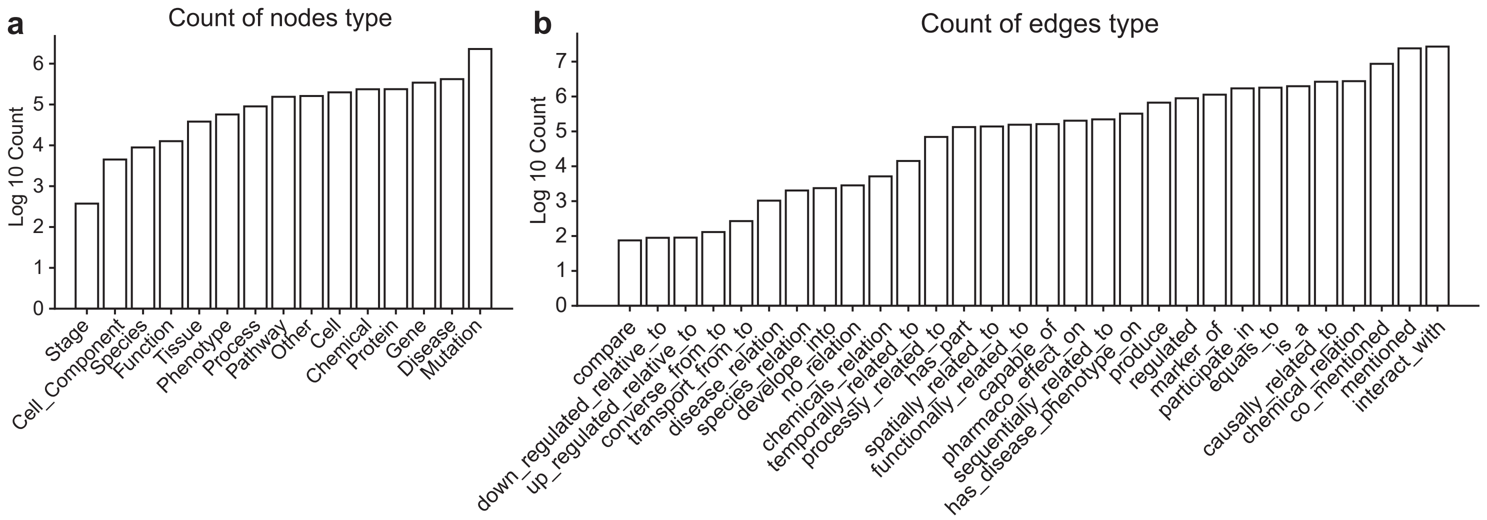


**Supplementary Fig S4. Statistics of the data amount of the BRICK knowledge graph. a.** Barplot of node type amount. y-axis: log10(nodes count). **b.** Barplot of edge type amount. y-axis: log10(edges count).

FigureS5

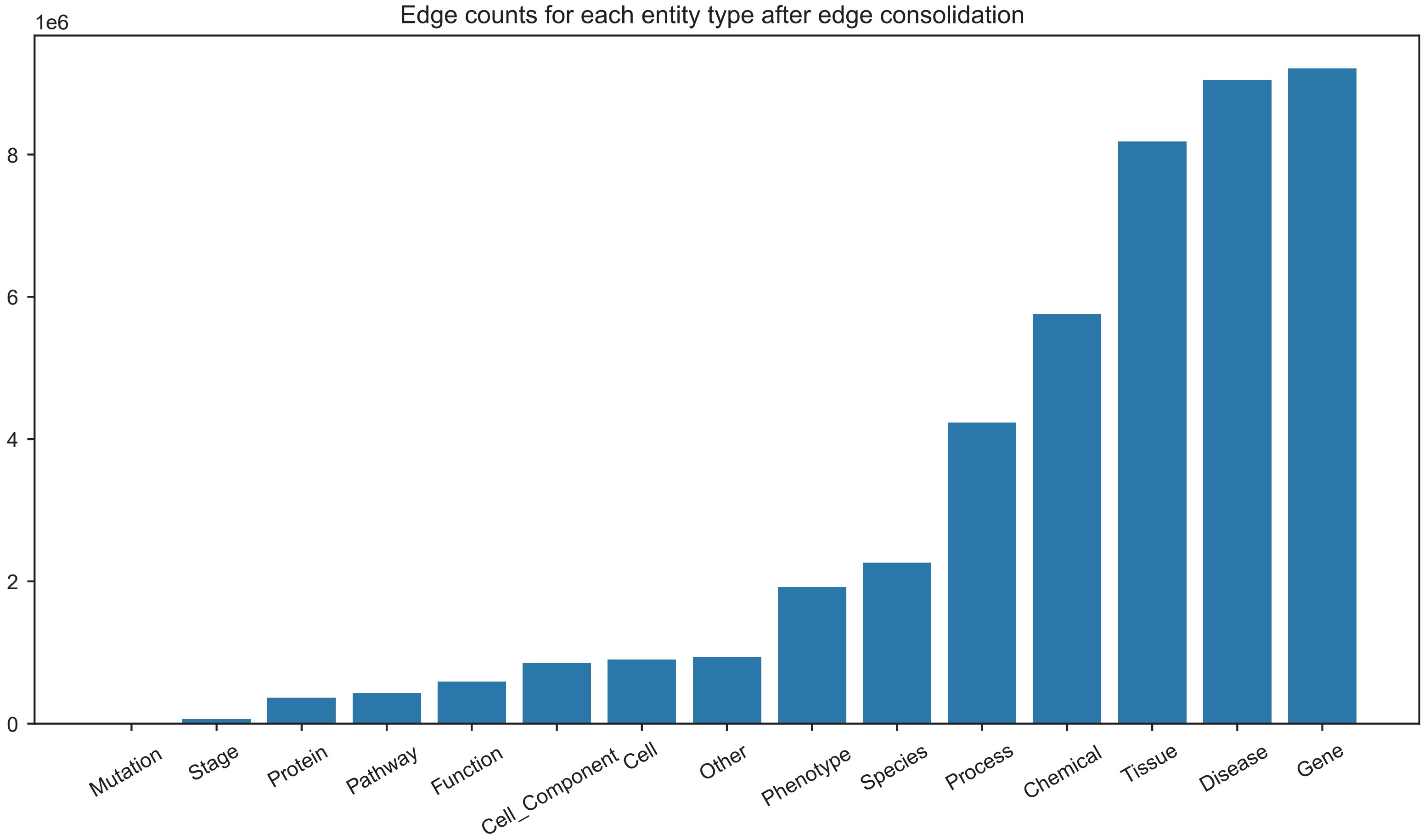


**Supplementary Fig S5. Edge count related to each entity type.**

FigureS6

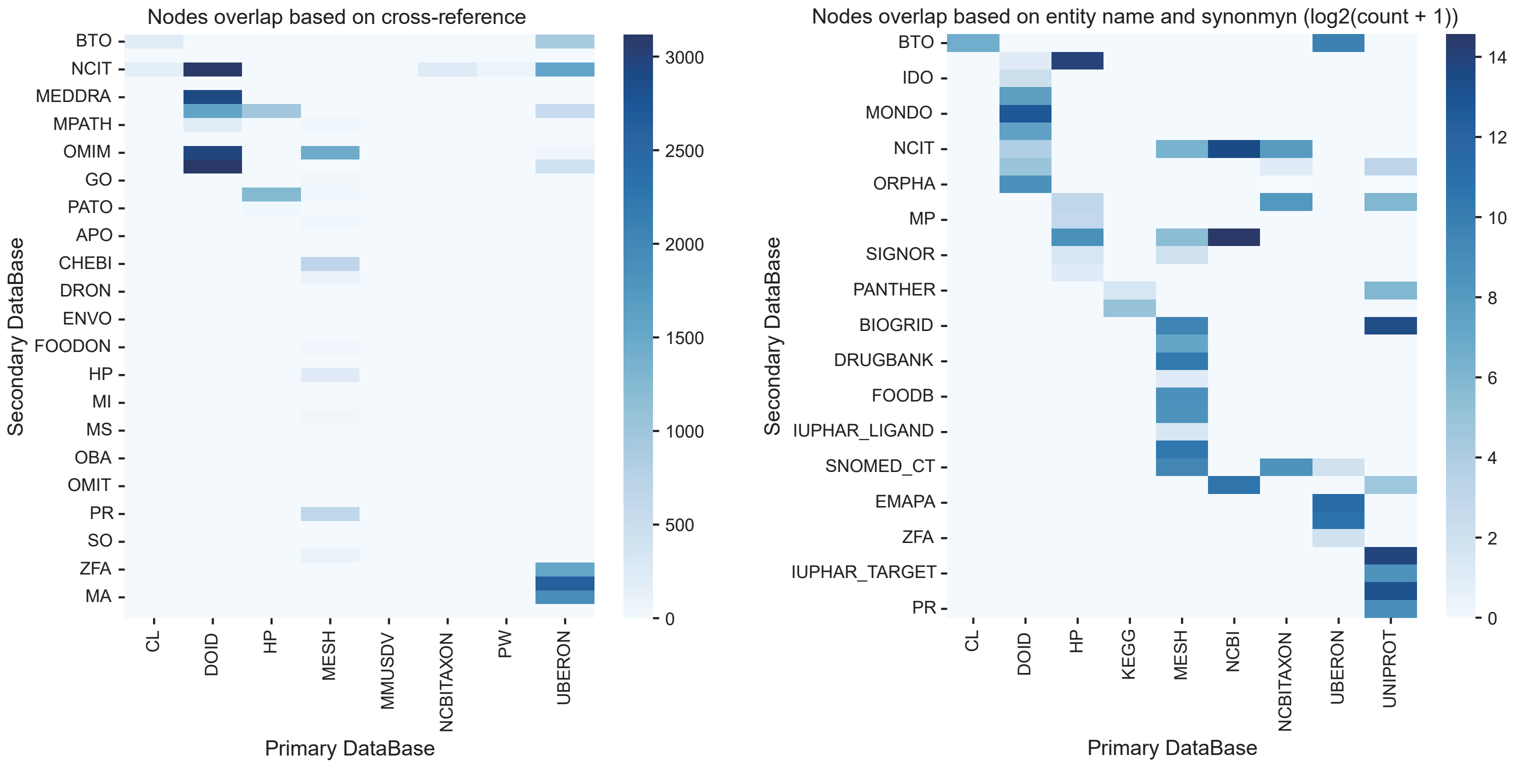


**Supplementary Fig S6. Heatmap of nodes overlap between different databases. left:** nodes overlap based on cross-reference, **right:** nodes overlap based on name and synonym

FigureS7

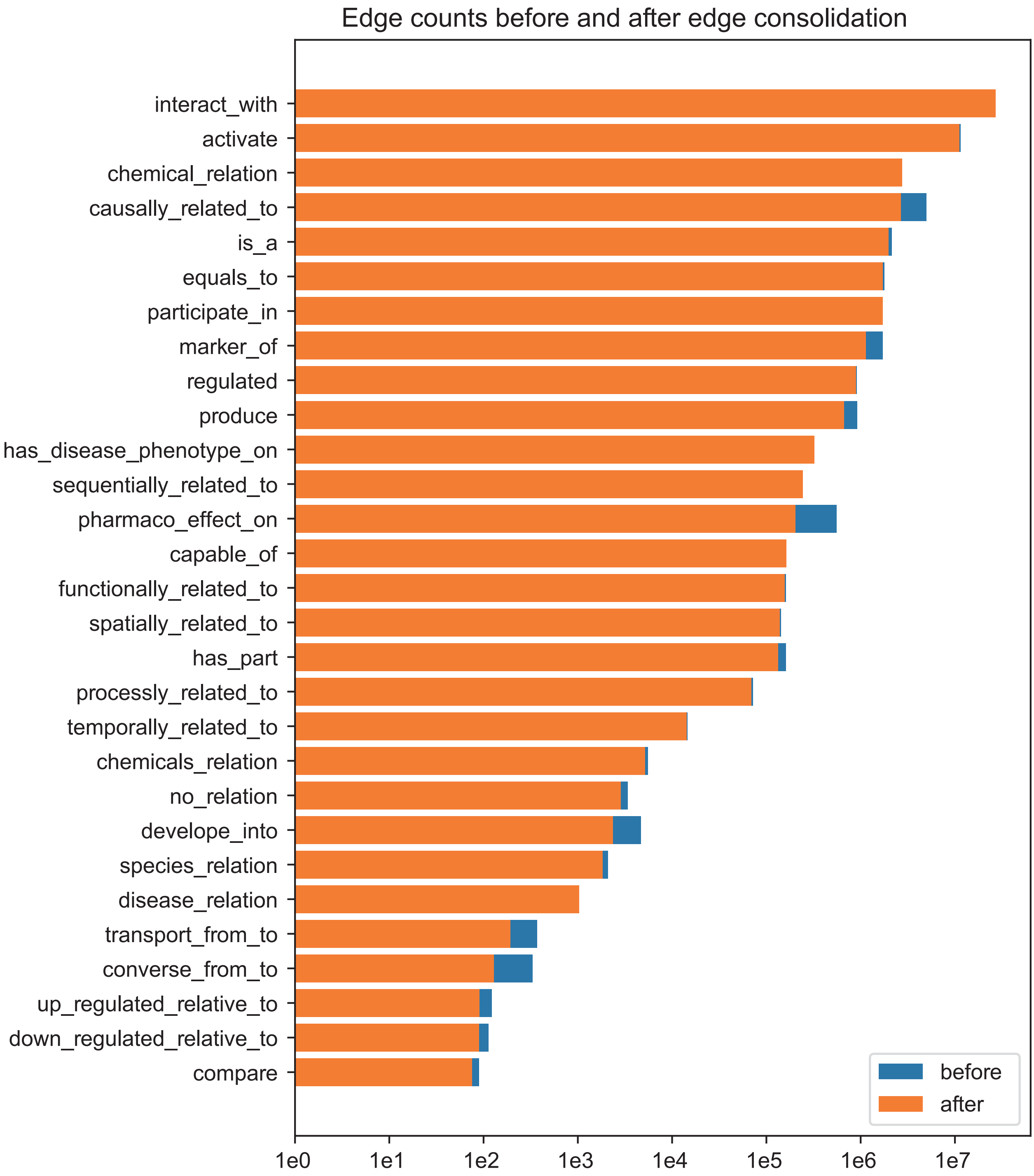


**Supplementary Fig S7. Edge counts before and after edge consolidation. blue: before, orange: after**

FigureS8

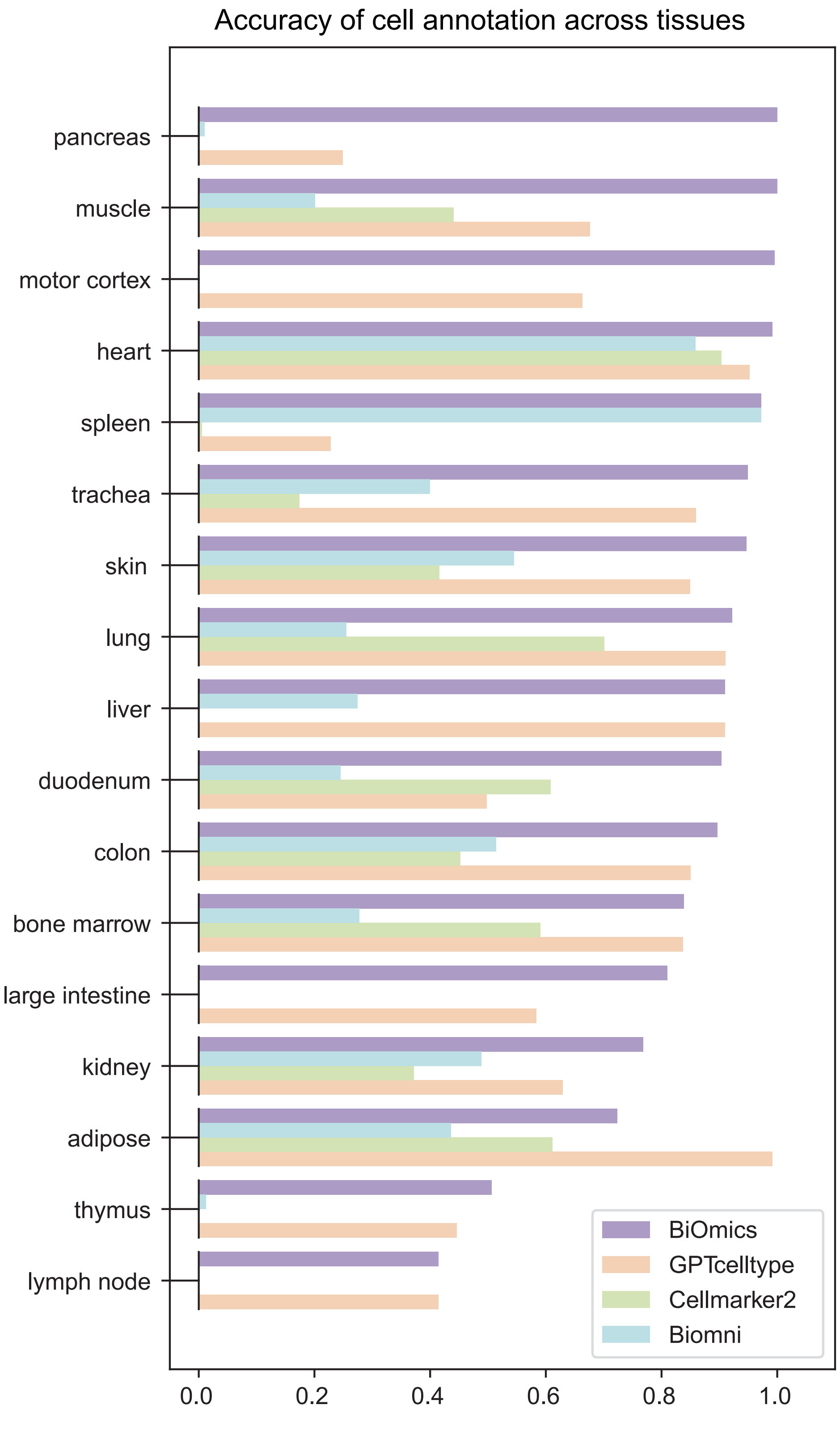


**Supplementary Fig S8. Benchmark of BiOmics annotation accuracy over 17 tissues. alternative methods including GPTcelltype, Cellmarker2 and Biomni.**

FigureS9

a

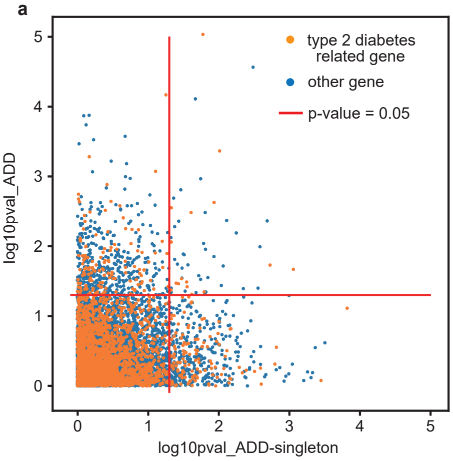


b

### 1. Reasoning the Analysis Result

- **Key Evidence Chain and Statistical Significance:**
  - The analysis identified several genes (e.g., LPAR6, SLC19A2, FYN, ZNF331, CHD7) that are significantly associated with Type 2 Diabetes Mellitus (T2DM). These genes were found to be enriched in various diseases, including genetic diseases, neoplasms, and specific types of cancer.
  - The match count for "type 2 diabetes mellitus" is 16, which is a relatively high number, indicating a strong association between these genes and T2DM.
  - The statistical significance is not explicitly provided, but the high match count suggests that the association is likely to be significant.

- **Reasonability of the Analysis Result:**
  - The identified genes are involved in various biological processes, such as cell signaling, metabolism, and immune response, which are known to play a role in the pathogenesis of T2DM.
  - For example, FYN is a tyrosine kinase that regulates insulin signaling, and its dysregulation has been linked to T2DM. Similarly, KEAP1 is involved in the regulation of oxidative stress, which is a key factor in T2DM.
  - The presence of these genes in multiple disease pathways (e.g., cancer, obesity) also supports the idea that they have pleiotropic effects, which is consistent with the multifactorial nature of T2DM.

- **Most Valuable Conclusion:**
  - The most valuable conclusion from this analysis is the identification of a set of genes (e.g., LPAR6, SLC19A2, FYN, ZNF331, CHD7) that are significantly associated with T2DM and are also implicated in other diseases. This suggests that these genes may be key players in the development and progression of T2DM and could serve as potential therapeutic targets.

### 2. Exploring New Biological Hypotheses

**Supplementary Fig S9. BiOmics analysis based on T2D GWAS analysis at gene level. a.** the scatter plot of GWAS gene related to T2D. x-axis: log 10 pvalue of ADD singleton; y-axis: log 10 pvalue of ADD; red line: p-value = 0.05; oranger point: T2D related genes, blue point: other genes. **b.** BiOmics interpretation context of T2D GWAS analysis at gene level.

FigureS10

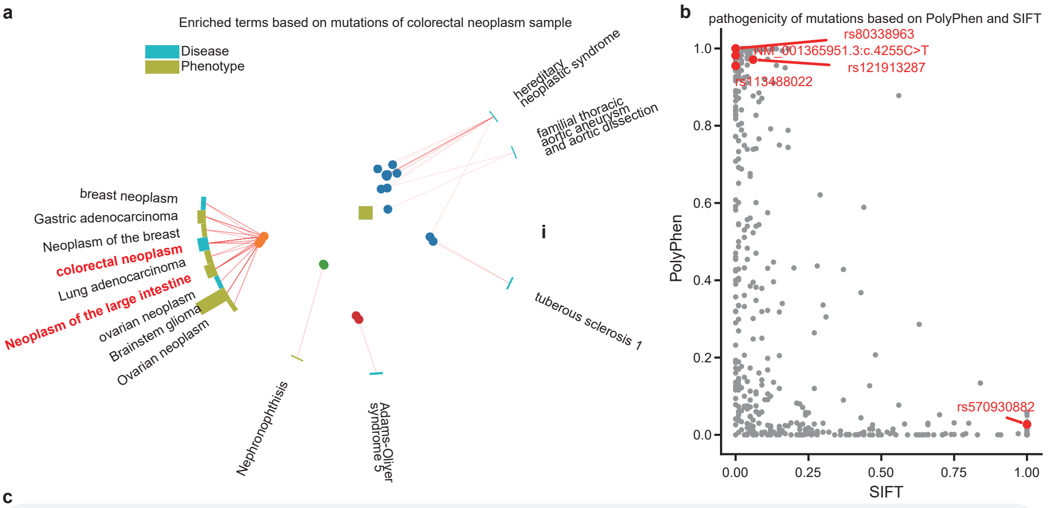


c

**### 1. Reasoning the Analysis Result**
- The presence of SNPs like rs80338963 and rs121913287 in multiple cancer-related phenotypes (e.g., Gastric adenocarcinoma, Neoplasm of the large intestine) is consistent with the known genetic basis of these cancers.
- The association of SNPs with specific diseases (e.g., Colorectal neoplasm, Hereditary neoplastic syndrome) is supported by existing literature, making the results reasonable.
- The most valuable conclusion is the identification of SNPs rs80338963 and rs121913287, which are consistently associated with multiple cancer types, particularly colorectal neoplasm. This suggests they may be key drivers or markers for these conditions.

**### 2. Exploring New Biological Hypotheses**
- Hypothesis 1: SNPs rs80338963 and rs121913287 may play a role in the development of multiple cancer types through shared signaling pathways.
- Hypothesis 2: These SNPs may also influence the progression and metastasis of tumors, as they are associated with different stages and types of neoplasms.

- **Clinical/Translational Medical Significance:**
  - Identifying these SNPs could lead to the development of targeted therapies for multiple cancer types, potentially improving patient outcomes.
  - These SNPs could serve as biomarkers for early detection and risk stratification in clinical settings, aiding in personalized medicine approaches.

**### 3. Advice for Following Analysis**
- **Gene Enrichment Analysis:**
  - Perform gene enrichment analysis to identify the biological pathways and functions associated with the identified SNPs. This will help in understanding the molecular mechanisms underlying the observed phenotypes.
- **Drug Repurposing Analysis:**
  - Conduct drug repurposing analysis to find existing drugs that target the pathways affected by these SNPs. This could lead to the discovery of new therapeutic options for the associated diseases.

**Supplementary Fig S10. BiOmics analysis and interpretation on TCGA colon cancer sample dataset. a.** Radial enrich plot of SNPs by BiOmics. The colors of the bars distinguish the types of enriched entities, and the heights of the bars represent the number of genes in the enriched entities. Meanwhile, mutations sets associated with adjacent bars have higher similarity. In UMAP scatter plot in the center of the circular bar chart, each point represent a mutation. The colors of the mutations represent the grouping of mutations. mutations that are close to each other in the UMAP also indicate a higher functional correlation. Red lines connect each mutation to its corresponding enriched term. **b.** Scatter plot of causal SNPs. x-axis: SIFT index, y-axis: PolyPhen index. BiOmics highlight the SNPs related to colon cancer. c. BiOmics interpretation context of analysis on TCGA colon cancer sample dataset.

FigureS11

Based on the queried table, the most reasonable annotation for this cell cluster is **type D enteroendocrine cell**. Here's why:

1. **Rank and Match Probability**: The "type D enteroendocrine cell" has the highest rank (rank 1) and the highest match probability (0.044860) among the candidate cell types.
2. **Subset-Superset Relations**: "Type D enteroendocrine cell" is a more specific subtype of the broader "enteroendocrine cell" category, which also appears in the list but with a lower rank and match probability.

Additionally, the presence of other enteroendocrine subtypes (e.g., "type A enteroendocrine cell") and pancreatic endocrine cells (e.g., "pancreatic PP cell" and "pancreatic endocrine cell") suggests that this cluster might be a mixture of different subtypes. Therefore, it is recommended to perform sub-cell type refinement in the next analysis step to further resolve these potential sub-populations.

**Conclusion**: This cell cluster is most likely annotated as **type D enteroendocrine cell**, but it may also contain a mixture of other enteroendocrine and pancreatic endocrine subtypes.

**Supplementary Fig S11. BiOmics interpretation context of annotation for #cluster2.**

FigureS12

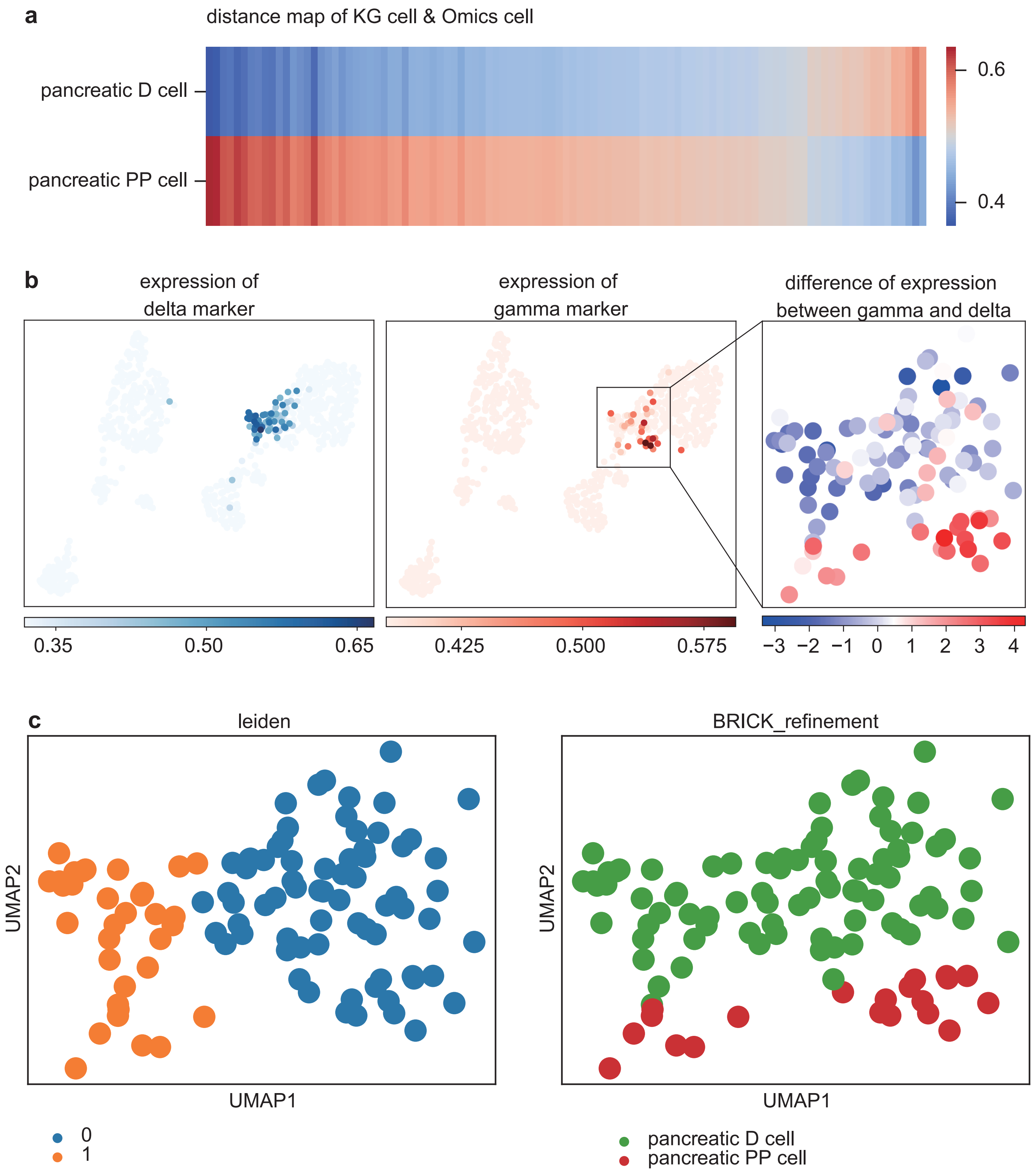


**Supplementary Fig S12. a.** Heatmap of distance map between KG cell and Omics cell. **b.** Expression pattern support refinement result. left. the UMAP of expression of delta cell markers. middle. the UMAP of expression of gamma cell markers. right. the UMAP of differential expression between gamma and delta cell markers. **c.** UMAP of sub-clustering of leiden cluster 2. left: leiden, right: BiOmics refinement result.

FigureS13

### Interpretation of Differential Expression Gene (DEG) Analysis in Pancreatic PP Cells

#### 1. Summary of Valuable Findings and Conclusions

**Overview:**
The DEG analysis of pancreatic PP cells has identified several genes that are differentially expressed, with significant p-values and log fold changes. The top differentially expressed genes (DEGs) include *Pcsk2*, *Pyy*, *Ppy*, *Peg3*, and *Pcsk1n*. These genes show high scores, low p-values, and substantial log fold changes, indicating their significant differential expression.

**Key Findings:**

- **Highly Upregulated Genes:**
  - *Pyy* (logFC: 6.055765, p-value: 4.579082e-12)
  - *Ppy* (logFC: 6.044227, p-value: 5.788614e-08)
  - *Ace2* (logFC: 7.329956, p-value: 4.464613e-05)
  - *Wnt4* (logFC: 4.289319, p-value: 2.697137e-05)
  - *Ugt2b34* (logFC: 4.896807, p-value: 5.487855e-05)

- **Genes with High Scores and Low P-values:**
  - *Pcsk2* (score: 18.791946, p-value: 5.048756e-19)
  - *Pyy* (score: 14.531603, p-value: 4.579082e-12)
  - *Ppy* (score: 8.546126, p-value: 5.788614e-08)
  - *Peg3* (score: 7.407349, p-value: 3.735858e-07)
  - *Pcsk1n* (score: 7.395367, p-value: 3.162213e-07)

- **Markers of Pancreatic PP Cells:**
  - From Table 2, several of the DEGs are known markers of pancreatic PP cells, including *Zc3h7b*, *Cplx2*, *Kif1b*, *Ugt2b34*, *Ocrl*, *Sez6l2*, *Cltrn*, *Scg2*, *Etv1*, *Scg5*, *Ptprn*, *Elapor1*, *Pnisr*, *Pyy*, *Peg3*, *Ppy*, *Pcsk1n*, *Rbp4*, *Gpx3*, *Ace2*, *Abcc8*, *Resp18*, *Wnt4*.

**Conclusions:**
- The highly upregulated genes, such as *Pyy*, *Ppy*, *Ace2*, *Wnt4*, and *Ugt2b34*, are likely to play crucial roles in the function and regulation of pancreatic PP cells.
- The significant differential expression of known PP cell markers (*Pyy*, *Ppy*, *Peg3*, *Pcsk1n*, etc.) confirms the specificity of the DEG analysis and highlights the importance of these genes in the cellular identity and function of PP cells.
- The identification of *Pcsk2* and *Pcsk1n* as highly differentially expressed genes suggests a potential role in the processing and maturation of prohormones and neuropeptides, which is a critical function of PP cells.

#### 2. Advice for Following Analysis and Research Objectives

**Further Analysis:**

1. **Functional Enrichment Analysis:**
   - Perform Gene Ontology (GO) and Kyoto Encyclopedia of Genes and Genomes (KEGG) pathway enrichment analysis to identify the biological processes, molecular functions, and pathways associated with the DEGs.
   - This will help in understanding the functional implications of the differentially expressed genes in PP cells.

2. **Protein-Protein Interaction (PPI) Network:**
   - Construct a PPI network using tools like STRING or Cytoscape to explore the interactions between the DEGs.
   - Identify key hub genes and subnetworks that may be involved in the regulation and function of PP cells.

3. **Validation of DEGs:**
   - Validate the expression levels of the top DEGs using qPCR or Western blotting to confirm the results from the RNA-seq data.
   - This will provide additional confidence in the DEG findings and help in prioritizing genes for further study.

4. **Single-Cell RNA-Seq Analysis:**
   - If possible, perform single-cell RNA-seq analysis to gain a more detailed understanding of the heterogeneity within the PP cell population.
   - This can help in identifying subpopulations of PP cells and their specific gene expression profiles.

**Research Objectives:**

1. **Mechanistic Studies:**
   - Investigate the molecular mechanisms by which the top DEGs, such as *Pyy*, *Ppy*, *Ace2*, and *Wnt4*, contribute to the function and regulation of PP cells.
   - Use CRISPR/Cas9 or siRNA knockdown experiments to study the effects of these genes on PP cell function and hormone secretion.

2. **Regulatory Networks:**
   - Explore the transcriptional and post-transcriptional regulatory networks that control the expression of the DEGs.
   - Identify key transcription factors and microRNAs that regulate the expression of these genes.

3. **Disease Relevance:**
   - Investigate the relevance of the DEGs in the context of pancreatic diseases, such as diabetes and pancreatitis.
   - Determine if the dysregulation of these genes contributes to the pathogenesis of these diseases and whether they can serve as potential therapeutic

**Supplementary Fig S13. BiOmics interpretation context of DEG task on pancreatics PP cell.**

FigureS14

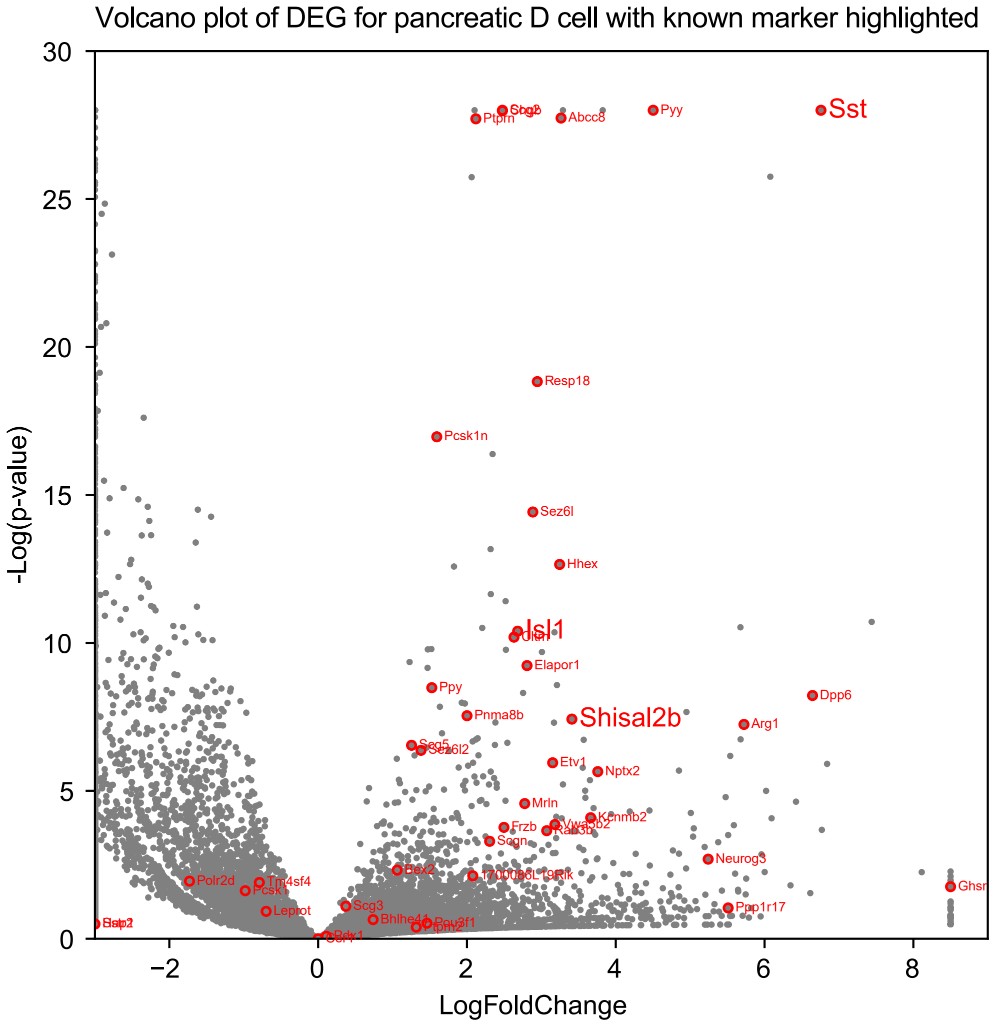


**Supplementary Fig S14. Volcano plot of differential expression gene of pancreatic D cell with known marker highlighted. x-axis: log-fold-change, y-axis: -log(p-value), each point represent a gene, known marker are circled in red and label gene name, the font size of gene name represent information source count of marker of relation between gene and pancreatic D cell.**

FigureS15

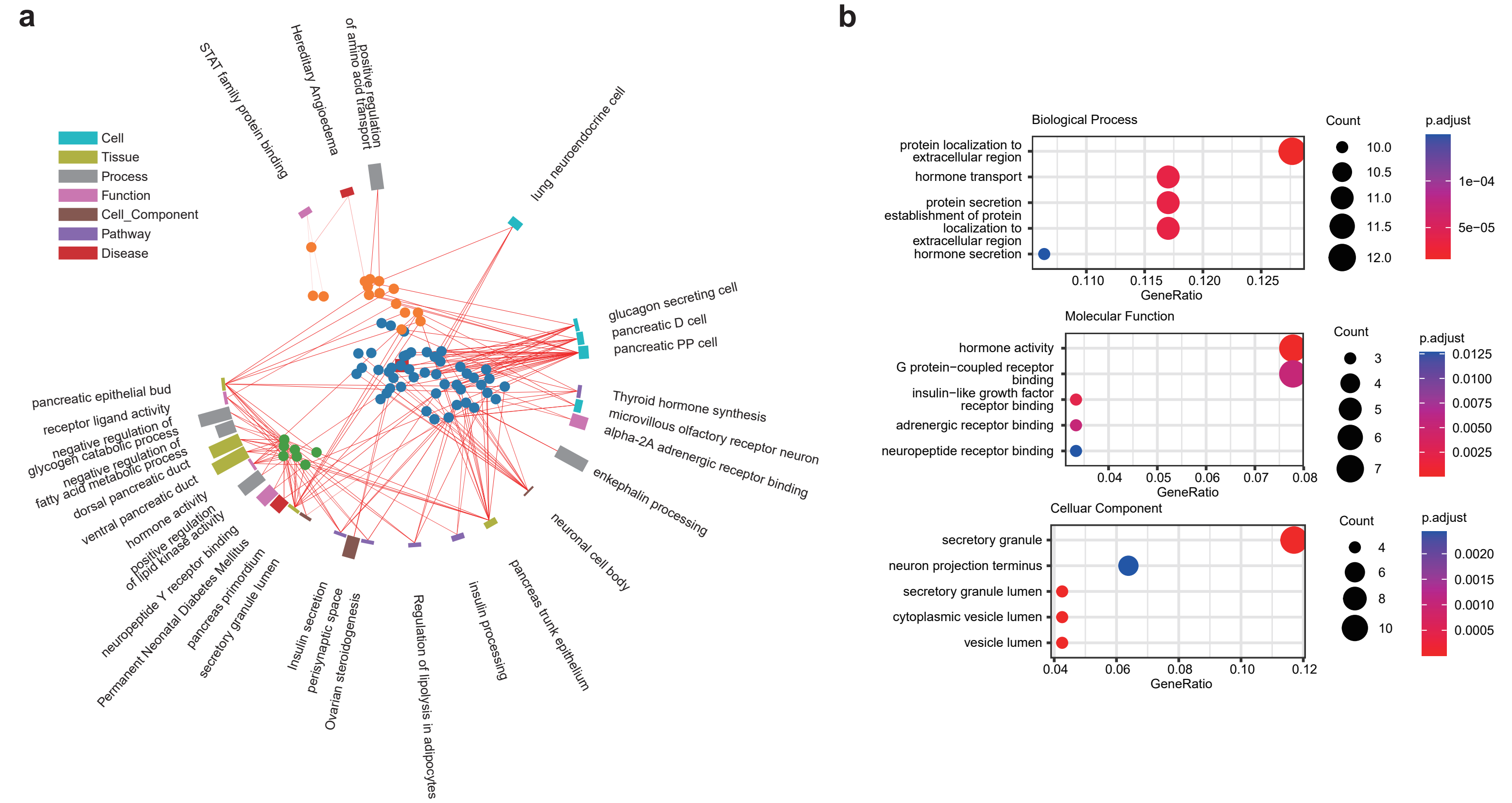


**Supplementary Fig S15. Gene enrichment of pancreatic D cell. a.** Radial enrich plot of marker of pancreatic D cell by BiOmics. The colors of the bars distinguish the types of enriched entities, and the heights of the bars represent the number of genes in the enriched entities. Meanwhile, gene sets associated with adjacent bars have higher similarity. In UMAP scatter plot in the center of the circular bar chart, each point represent a gene. The colors of the genes represent the grouping of genes. Genes that are close to each other in the UMAP also indicate a higher functional correlation. Red lines connect each gene to its corresponding enriched term. **b.** Traditional gene enrichment dotplot by clusterProfiler. The size of each dot represent enrich count, x-axis: gene ratio, color: p-value.

FigureS16

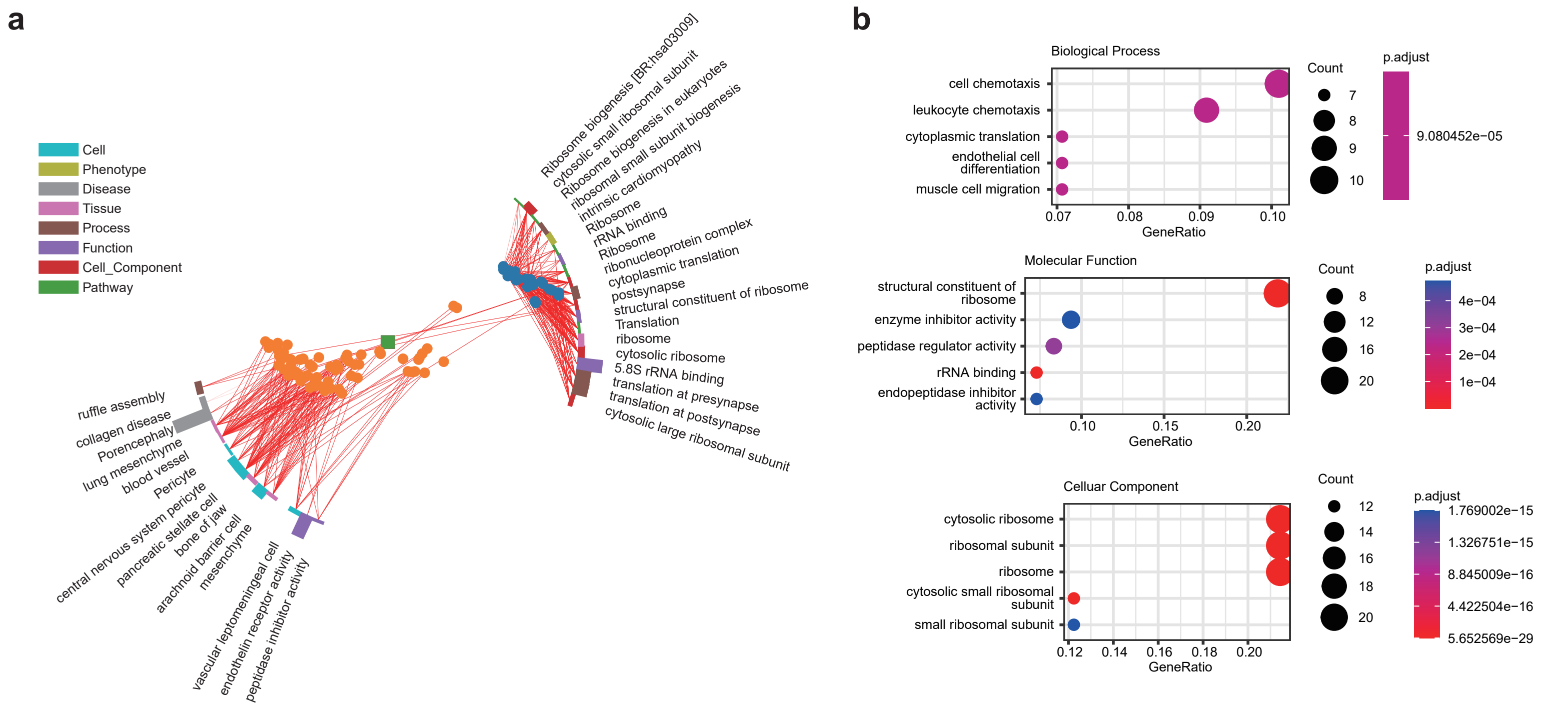


**Supplementary Fig S16. Gene enrichment of macrophage. a.** Radial enrich plot of marker of macrophage by BiOmics. The colors of the bars distinguish the types of enriched entities, and the heights of the bars represent the number of genes in the enriched entities. Meanwhile, gene sets associated with adjacent bars have higher similarity. In UMAP scatter plot in the center of the circular bar chart, each point represent a gene. The colors of the genes represent the grouping of genes. Genes that are close to each other in the UMAP also indicate a higher functional correlation. Red lines connect each gene to its corresponding enriched term. **b.** Traditional gene enrichment dotplot by clusterProfiler. The size of each dot represent enrich count, x-axis: gene ratio, color: p-value.

FigureS17

### Interpretation of Gene Enrichment Results for Pancreatic PP Cells

#### 1. Summary and Valuable Findings from Table 1

**Table 1** provides the gene enrichment results for pancreatic PP cells, focusing on pathways and cell types. Here are the key findings:

- **Glucagon-Secreting Cell (path.0.name: [Ttr, Serpina10, Pyy, Scgn, Abcc8, Scg5, Scg2, ...])**:
  - This pathway is significantly enriched with a p-value of \(8.09 \times 10^{-43}\).
  - It includes genes such as Ttr, Serpina10, Pyy, Scgn, Abcc8, Scg5, and Scg2, which are known to be involved in the function and regulation of glucagon-secreting cells.
  - The high match count (39) indicates a strong association between these genes and the glucagon-secreting cell type.

- **Steroid Hormone Secreting Cell (path.0.name: [Meg3, Isl1, Foxa2, Gria2, Gcg, Cplx2, Cpe, Ch...])**:
  - This pathway is also significantly enriched with a p-value of \(8.71 \times 10^{-36}\).
  - Genes like Meg3, Isl1, Foxa2, Gria2, Gcg, Cplx2, Cpe, and Chga are associated with steroid hormone secretion.
  - The match count (33) suggests a substantial overlap between the genes and the steroid hormone secreting cell type.

- **Myoendocrine Cell (path.0.name: [Isl1, Foxa2, Gria2, Gcg, Cplx2, Cpe, Chga, Sl...])**:
  - This pathway has a p-value of \(1.36 \times 10^{-35}\).
  - Genes such as Isl1, Foxa2, Gria2, Gcg, Cplx2, Cpe, Chga, and Slc7a2 are involved in myoendocrine cell function.
  - The match count (34) indicates a strong association with this cell type.

- **Pancreatic Endocrine Cell (path.0.name: [Cplx2, Cpe, Chga, Serping1, Slc7a2, Arx, Aplp...])**:
  - This pathway is significantly enriched with a p-value of \(2.74 \times 10^{-35}\).
  - Genes like Cplx2, Cpe, Chga, Serping1, Slc7a2, Arx, and Aplp2 are associated with pancreatic endocrine cell function.
  - The match count (38) suggests a strong association with this cell type.

- **Myocardial Endocrine Cell (path.0.name: [Scg5, Scg2, Resp18, Ptprn, Ppy, Peg3, Pcsk2, ...])**:
  - This pathway has a p-value of \(1.02 \times 10^{-34}\).
  - Genes such as Scg5, Scg2, Resp18, Ptprn, Ppy, Peg3, Pcsk2, and others are involved in myocardial endocrine cell function.
  - The match count (29) indicates a significant association with this cell type.

- **Other Pathways**:
  - There are several other pathways with lower match counts and higher p-values, indicating less significant associations. For example, perisynaptic space, non-syndromic X-linked intellectual disability, dendrite cytoplasm, chromaffin granule, and X chromosome.

#### 2. Interpretation of Gene Sub-Clusters from Table 2

**Table 2** provides further sub-cluster analysis of the gene enrichment results. Here are the key findings for each sub-cluster:

- **Sub-Cluster 0**:
  - **Pathway: Glucagon-Secreting Cell (path.0.name: [Ttr, Serpina10, Pyy, Scgn, Abcc8, Scg5, Scg2, ...])**
    - This sub-cluster is strongly associated with the glucagon-secreting cell type, with a match count of 16.
    - The p-value is \(8.09 \times 10^{-43}\), indicating a highly significant enrichment.
    - Key genes include Ttr, Serpina10, Pyy, Scgn, Abcc8, Scg5, and Scg2, which are crucial for glucagon secretion and regulation.

  - **Pathway: Pancreatic PP Cell (path.0.name: [Zc3h7b, Cplx2, Kif1b, Slc16a12, Ugt2b34, Ocrl...])**
    - This sub-cluster is specifically associated with the pancreatic PP cell type, with a match count of 13.
    - The p-value is not provided, but the high match count suggests a significant association.
    - Key genes include Zc3h7b, Cplx2, Kif1b, Slc16a12, Ugt2b34, and Ocrl, which are involved in the specific functions of PP cells.

- **Sub-Cluster 1**:
  - **Pathway: Metanephros (path.0.name: [Kif1b, Etv1, Tshz3, Bcl3, Hdac6, Pgrmc1, Cnot...])**
    - This sub-cluster is associated with the metanephros, with a match count of 12.
    - The p-value is not provided, but the match count suggests a significant association.
    - Key genes include Kif1b, Etv1, Tshz3, Bcl3, Hdac6, Pgrmc1, and Cnot, which are involved in kidney development and function.

  - **Pathway: Kidney (path.0.name: [Meis2, Serpina10, Phip, Irx1, Khdrbs3, Klf10,...])**
    - This sub-cluster is associated with the kidney, with a match count of 12.
    - The p-value is not provided, but the match count suggests a significant association.
    - Key genes include Meis2, Serpina10, Phip, Irx1, Khdrbs3, Klf10, and others, which are involved in kidney function and development.

- **Sub-Cluster 2**:
  - **Pathway: Myoendocrine Cell (path.0.name: [Isl1, Foxa2, Gria2, Gcg, Cplx2, Cpe, Chga, Sl...])**
    - This sub-cluster is associated with the myoendocrine cell type, with a match count of 11.
    - The p-value is \(1.36 \times 10^{-35}\), indicating a highly significant enrichment.
    - Key genes include Isl1, Foxa2, Gria2, Gcg, Cplx2, Cpe, Chga, and Slc7a2, which are involved in myoendocrine cell function.

- **Sub-Cluster 3**:
  - **Pathway: Pseudohypoparathyroidism (path.0.name: [Gnas])**
    - This sub-cluster is associated with pseudohypoparathyroidism, with a match count of 1.
    - The p-value is not provided, but the low match count suggests a less significant association.
    - Key gene: Gnas, which is involved in parathyroid hormone resistance.

  - **Pathway: Albright's Hereditary Osteodystrophy (path.0.name: [Gnas])**
    - This sub-cluster is associated with Albright's hereditary osteodystrophy, with a match count of 1.
    - The p-value is not provided, but the low match count suggests a less significant association.
    - Key gene: Gnas, which is involved in this genetic disorder.

  - **Pathway: ITM2B-Related Cerebral Amyloid Angiopathy 1 (path.0.name: [Itm2b])**
    - This sub-cluster is associated with ITM2B-related cerebral amyloid angiopathy 1, with a match count of 1.
    - The p-value is not provided, but the low match count suggests a less significant association.
    - Key gene: Itm2b, which is involved in this neurological condition.

  - **Pathway: Maturity Onset Diabetes of the Young (path.0.name: [Foxa2, Pax6])**
    - This sub-cluster is associated with maturity onset diabetes of the young, with a match count of 0.
    - The p-value is not provided, and the zero match count suggests no significant association.
    - Key genes: Foxa2 and Pax6, which are involved in pancreatic development and function.

  - **Pathway: Nicotine Addiction (path.0.name: [Gria2, Gprin1])**
    - This sub-cluster is associated with nicotine addiction, with a match count of 0.
    - The p-value is not provided, and the zero match count suggests no significant association.
    - Key genes: Gria2 and Gprin1, which are involved in neuronal function and addiction.

### Conclusions

- **Primary Associations**: The primary associations for pancreatic PP cells are with glucagon-secreting cells, pancreatic endocrine cells, and myoendocrine cells. These pathways are highly enriched and have strong p-values, indicating their importance in the function and regulation of PP cells.
- **Secondary Associations**: There are secondary associations with steroid hormone-secreting cells and myocardial endocrine cells, which also show significant enrichment.
- **Gene Sub-Clusters**: The sub-cluster analysis further refines the associations, highlighting specific pathways and cell types. Sub-cluster 0, which includes glucagon-secreting cells and pancreatic PP cells, is particularly significant. Other sub-clusters, such as those related to kidney function and myoendocrine cells, also show notable associations.
- **Less Significant Associations**: Some sub-clusters, such as those related to pseudohypoparathyroidism, Albright's hereditary osteodystrophy, and ITM2B-related cerebral amyloid angiopathy, show less significant associations, with low match counts and no provided p-values.

This analysis provides a comprehensive understanding of the gene enrichment in pancreatic PP cells and highlights the key pathways and cell types involved in their function.

**Supplementary Fig S17. BiOmics interpretation context of gene enrichment task on pancreatics PP cell.**

FigureS18

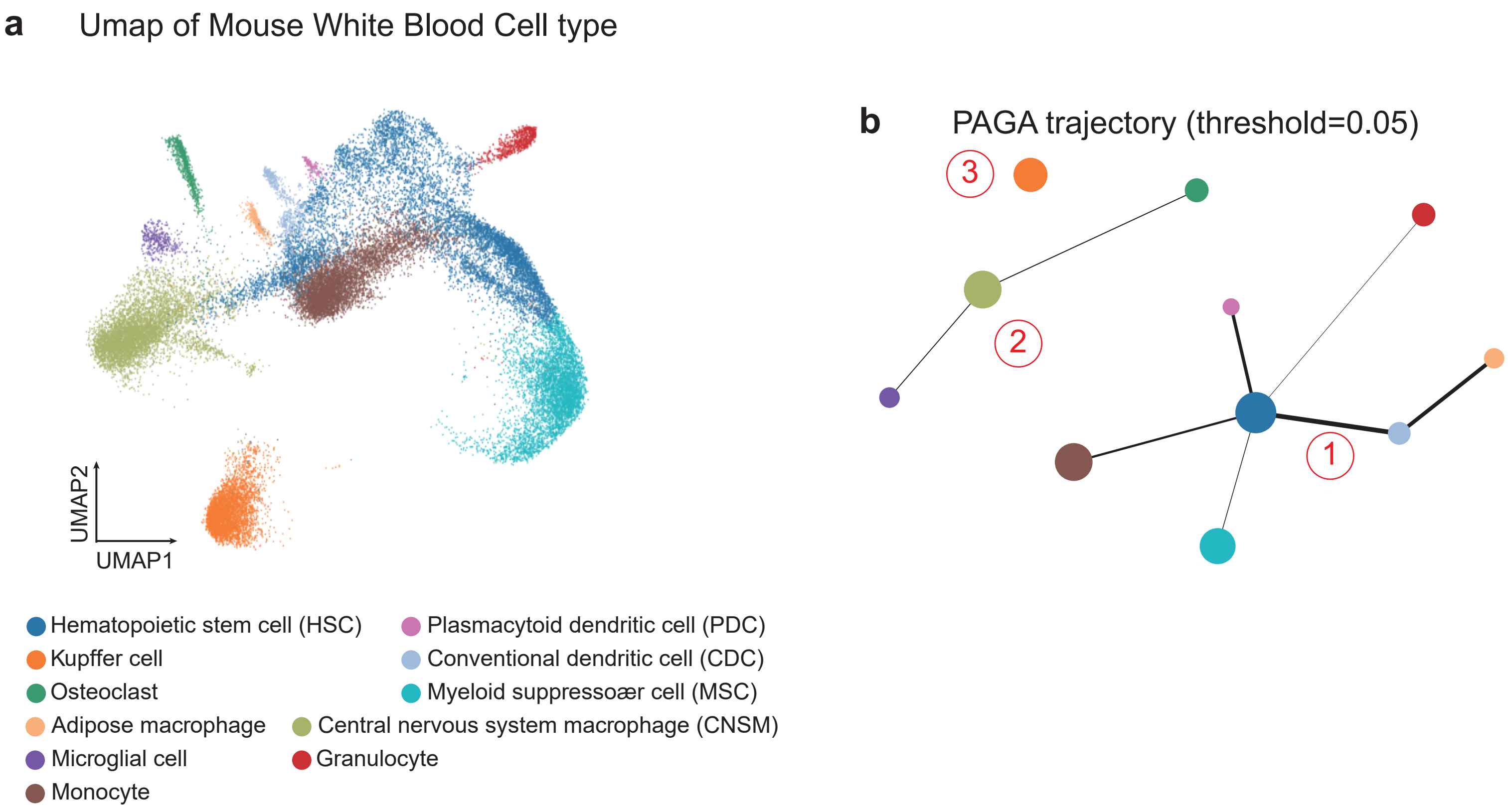


**Supplementary Fig S18. Preprocess on Mouse White Blood cell dataset. a.** UMAP projections of whole WBCs dataset with eleven cell type clusters (Central nervous system macrophage (CNSM), Hematopoietic stem cell (HSC), Monocyte, Kupffer cell, Myeloid suppressor cell (MSC), Osteoclast, Granulocyte, Conventional dendritic cell (CDC), Plasmacytoid dendritic cell (PDC), Microglial cell and Adipose macrophage). **b.** PAGA trajectory of WBCs dataset, which was split into 3 groups under threshold=0.05.

FigureS19

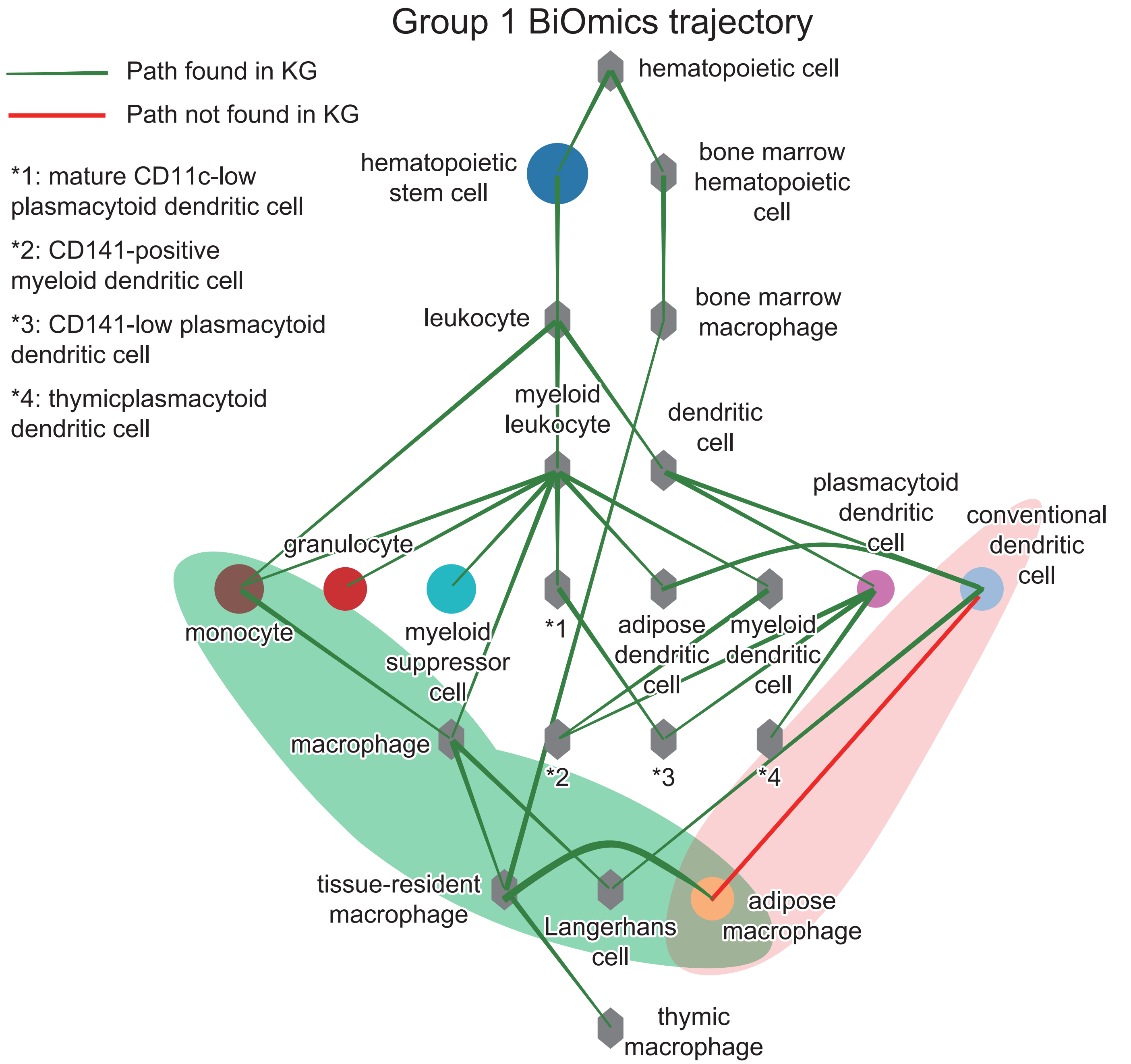


**Supplementary Fig S19. The complete trajectory graph about cell group in bone marrow (Granulocyte, HSC, MSC, PDC, CDC, Monocyte and Adipose macrophage) after BRICK completion the PAGA graph with Knowledge graph. The colorful circle nodes represents the cell types from dataset and the other gray hexagon nodes represents the cell types from knowledge graph. The green path represents verified path and the red path represents unverified path.**

FigureS20

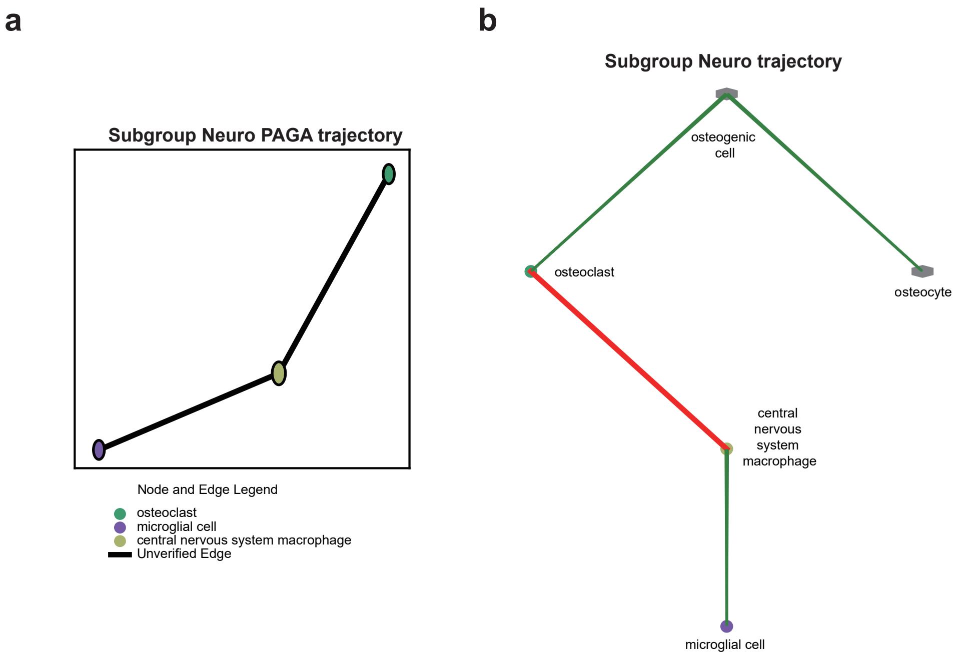


**Supplementary Fig S20. BiOmics analysis and interpretation on Neuro subgroup. a** PAGA trajectory graph on Neuro subgroup. **b.** The BiOmics completion trajectory graph on Neuro subgroup.

FigureS21

Generated Answer: The provided cell developmental trajectory graph reveals a highly interconnected lineage map, illustrating the differentiation pathways of hematopoietic cells into diverse immune cell types. The graph's integration of RNAseq data and a biological knowledge graph allows for an enriched representation of both experimentally observed and computationally inferred trajectories, marked by green and red edges, respectively. At its core, the hematopoietic stem cell (HSC) serves as the progenitor, giving rise to leukocytes, which then branch into various specialized immune lineages, including monocytes, granulocytes, dendritic cells, and macrophages, among others. This trajectory mirrors the hierarchical and tightly regulated process of hematopoiesis, orchestrated by key transcription factors such as GATA2, PU.1, and RUNX1, which guide lineage commitment and differentiation through gene expression programs. For instance, the differentiation of monocytes into macrophages and tissue-resident macrophages is likely governed by the macrophage colony-stimulating factor (M-CSF) signaling pathway and transcription factors like C/EBPα and MafB, which regulate genes essential for phagocytosis and tissue-specific functionality. Similarly, the development of dendritic cells, including plasmacytoid and conventional subsets, involves the IRF family of transcription factors, such as IRF8 and IRF4, which modulate antigen presentation and immune activation pathways.

The graph also highlights the plasticity within the leukocyte lineage, as evidenced by the multiple pathways originating from myeloid leukocytes, leading to granulocytes, macrophages, myeloid suppressor cells, and dendritic cells. This plasticity reflects the dynamic interplay of cytokine signaling (e.g., GM-CSF and IL-6) and epigenetic modifications that enable cells to respond to environmental cues and pathological conditions. Notably, the red edge linking conventional dendritic cells to adipose macrophages suggests a computationally inferred, potentially novel pathway that could represent a transdifferentiation event or a shared precursor cell. Such a connection warrants further experimental validation and exploration of the molecular mechanisms, possibly involving reprogramming factors or metabolic shifts associated with adipose tissue environments.

The presence of gray nodes representing knowledge graph-derived data enhances the interpretability of the graph by linking established biological insights to experimental findings. For example, tissue-resident macrophages are shown to differentiate into adipose macrophages or bone marrow macrophages, reflecting their adaptative roles in distinct tissue niches. This adaptation is likely mediated by local microenvironmental signals such as hypoxia, growth factors, and lipid metabolites, which influence macrophage polarization and function through pathways like HIF-1α and PPARγ signaling. Moreover, the development of Langerhans cells from macrophages and conventional dendritic cells underscores the importance of TGF-β signaling and local epithelial cues in shaping the unique antigen surveillance capabilities of these cells in the skin.

Overall, this graph captures the intricacies of hematopoiesis and immune cell differentiation, emphasizing the interplay between transcriptional regulation, signaling pathways, and environmental factors. The integration of inferred pathways (red edges) alongside experimentally validated ones (green edges) not only provides a comprehensive view of known developmental routes but also offers insights into potentially novel or underexplored transitions. Understanding these pathways at the molecular level could inform therapeutic strategies targeting immune cell function in diseases such as cancer, autoimmune disorders, and tissue repair.

**Supplementary Fig S21. BiOmics interpretation context of trajectory analysis on mouse WBC dataset.**

FigureS22

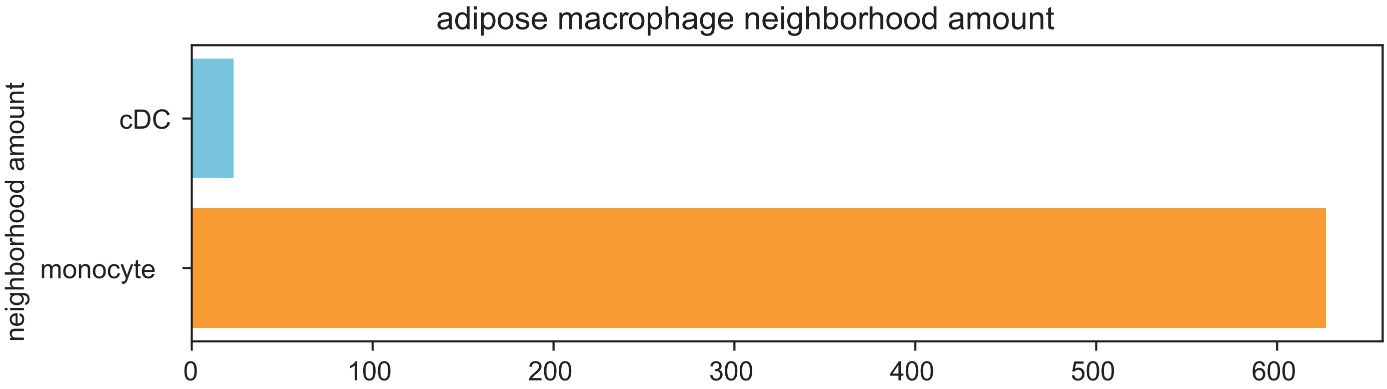


**Supplementary Fig S22. Neighbor count of adipose macrophage comparing CDC and monocyte by KNN graph.**

FigureS23

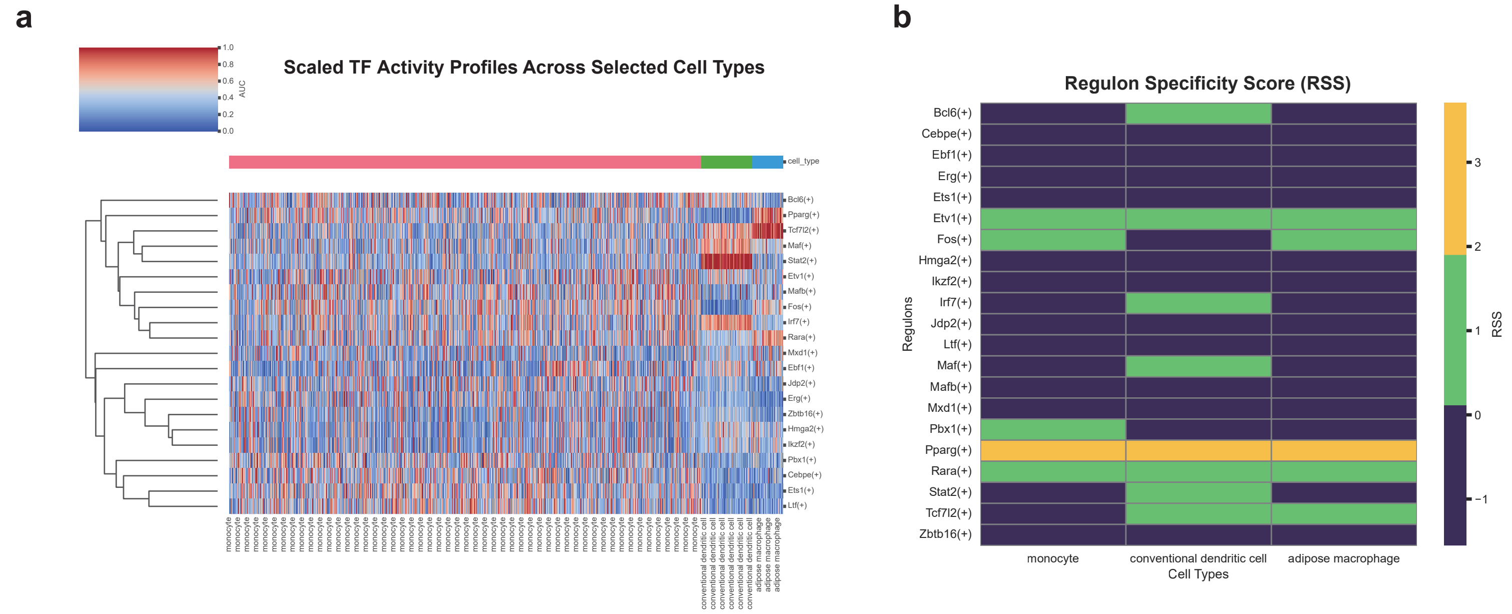


**Supplementary Fig S23. pySCENIC analysis on mouse WBC dataset. a** Scaled TF activity profiles across selected cell types. **b.** Regulon Specificity Score for each regulon across selected cell types.

FigureS24

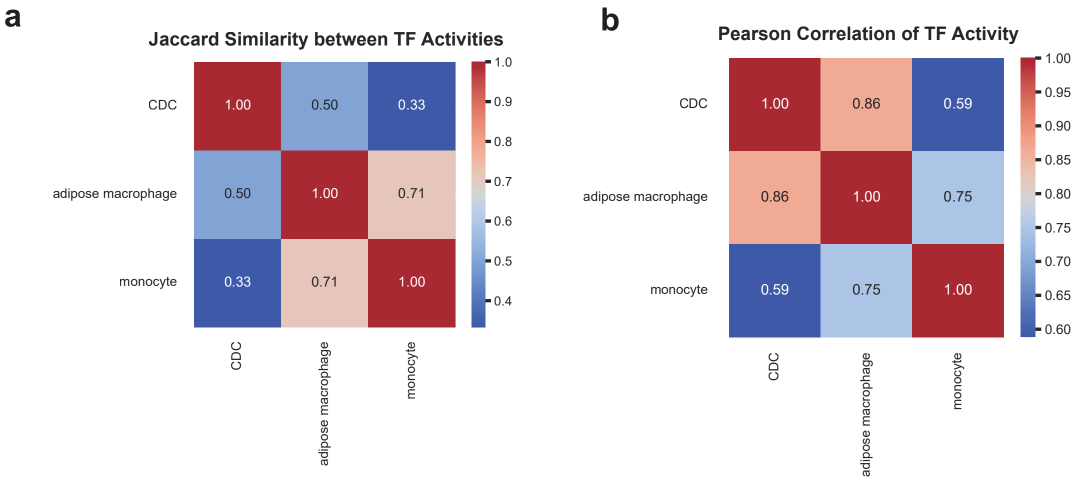


**Supplementary Fig S24. TF activities similiarity between select cell types. a.** Jaccard similiarity, **b.** pearson correlation.

FigureS25

### Analysis of the TF (Fos) and Its Regulon in Monocyte to Adipose Macrophage Differentiation

#### 1. Overview of the Transcription Factor Fos
The transcription factor Fos is a member of the AP-1 (Activator Protein-1) family, which plays a crucial role in various cellular processes, including cell proliferation, differentiation, and stress response. Fos forms heterodimers with Jun proteins to bind to the DNA at specific promoter regions, thereby regulating gene expression. In the context of monocyte to adipose macrophage differentiation, Fos has been implicated in modulating the inflammatory and metabolic functions of macrophages.

#### 2. Verified Genes Regulated by Fos
The verified genes list contains experimentally validated targets of Fos. These genes are known to be regulated by Fos and play significant roles in the development and function of adipose macrophages. Here are some key genes and their functions:

- **Cd163**: A scavenger receptor involved in the clearance of hemoglobin-haptoglobin complexes and other ligands. It is highly expressed in anti-inflammatory M2 macrophages.
- **Dhx58**: An RNA helicase that plays a role in antiviral defense and innate immune responses.
- **Rflnb**: Involved in the regulation of lipid metabolism and inflammation.
- **Lpo**: Lipoxygenase, which is involved in the synthesis of eicosanoids, important mediators of inflammation.
- **Pparg**: Peroxisome proliferator-activated receptor gamma, a key regulator of adipogenesis and anti-inflammatory responses in macrophages.
- **Nos2**: Inducible nitric oxide synthase, which produces nitric oxide, a key mediator of inflammatory responses.
- **Ifng**: Interferon-gamma, a cytokine that promotes the pro-inflammatory M1 phenotype in macrophages.
- **Cxcl10**: Chemokine (C-X-C motif) ligand 10, which recruits immune cells to sites of inflammation.
- **Il1r2**: Interleukin-1 receptor type 2, which acts as a decoy receptor for IL-1, modulating the inflammatory response.
- **Il15ra**: Interleukin-15 receptor subunit alpha, involved in the activation and survival of natural killer (NK) cells and T cells.

These genes collectively suggest that Fos regulates a wide range of processes, including inflammation, lipid metabolism, and immune responses, which are critical for the transition from monocytes to adipose macrophages.

#### 3. Potential Genes in the Regulon
The regulon list includes additional genes that are predicted to be regulated by Fos but have not yet been experimentally verified. Some of these genes include:

- **Mir22hg**: A long non-coding RNA that may play a role in gene regulation.
- **Edil3**: Endothelial cell-specific molecule 1, involved in angiogenesis and vascular remodeling.
- **Ahnak**: A large protein involved in cell adhesion and signaling.
- **Vcan**: Versican, an extracellular matrix proteoglycan involved in cell adhesion and migration.
- **Fn1**: Fibronectin 1, a glycoprotein that plays a role in cell adhesion and migration.
- **Adgrl3**: Adhesion G protein-coupled receptor L3, involved in cell adhesion and signaling.
- **Rara**: Retinoic acid receptor alpha, a nuclear receptor that regulates gene expression in response to retinoic acid.

While these genes are predicted to be regulated by Fos, further experimental validation is required to confirm their regulatory relationship. Their potential roles in monocyte to adipose macrophage differentiation include:
- **Mir22hg** and **Rara** may influence gene expression and cellular differentiation.
- **Edil3**, **Ahnak**, **Vcan**, and **Fn1** are involved in cell adhesion and extracellular matrix interactions, which are important for macrophage migration and tissue remodeling.
- **Adgrl3** may play a role in cell signaling and adhesion, contributing to the functional specialization of macrophages.

#### 4. Functional Implications of the Fos-Regulon in Monocyte to Adipose Macrophage Differentiation
The Fos-regulon appears to be involved in several key processes during the differentiation of monocytes into adipose macrophages:

- **Inflammation and Immune Response**: Genes like *Ifng*, *Cxcl10*, *Il1r2*, and *Il15ra* are involved in the regulation of inflammatory responses and immune cell recruitment, suggesting that Fos plays a role in modulating the inflammatory state of macrophages.
- **Lipid Metabolism and Homeostasis**: Genes such as *Pparg* and *Lpo* are involved in lipid metabolism and homeostasis, indicating that Fos may regulate the metabolic functions of adipose macrophages.
- **Cell Adhesion and Migration**: Genes like *Vcan*, *Fn1*, and *Adgrl3* are involved in cell adhesion and migration, which are essential for the proper localization and function of macrophages in adipose tissue.
- **Gene Expression and Cellular Differentiation**: *Mir22hg* and *Rara* may influence gene expression and cellular differentiation, potentially affecting the phenotypic switch between pro-inflammatory (M1) and anti-inflammatory (M2) macrophages.

#### 5. Conclusion and Recommendations
The Fos-regulon is a complex network of genes that collectively regulate various aspects of monocyte to adipose macrophage differentiation. The verified genes provide strong evidence for the role of Fos in inflammation, lipid metabolism, and immune responses. The potential genes in the regulon, while not yet experimentally validated, suggest additional layers of regulation that may be important for the functional specialization of adipose macrophages.

To further understand the role of Fos in this process, it is recommended to:
- **Experimentally validate** the regulatory relationship between Fos and the potential genes in the regulon.
- **Perform functional studies** to determine the specific roles of these genes in monocyte to adipose macrophage differentiation.
- **Investigate the interplay** between Fos and other transcription factors that may co-regulate the same set of genes.

By doing so, we can gain a more comprehensive understanding of the molecular mechanisms underlying the development and function of adipose macrophages.

**Supplementary Fig S25. BiOmics interpretation context of GRN analysis on mouse WBC dataset.**

FigureS26

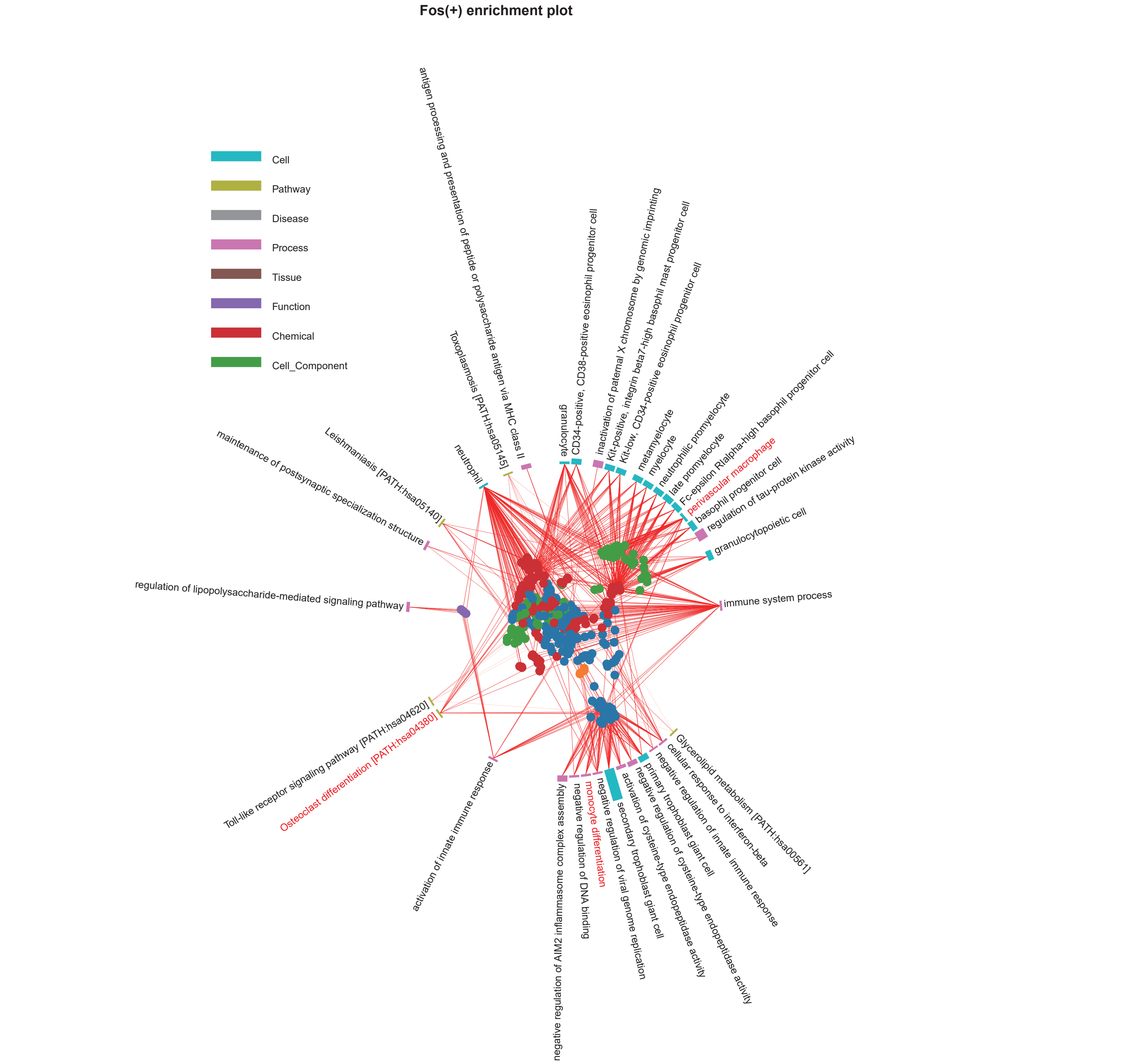


**Supplementary Fig S26. Radial enrich plot of Fos(+) by BiOmics. The colors of the bars distinguish the types of enriched entities, and the heights of the bars represent the number of genes in the enriched entities. Meanwhile, gene sets associated with adjacent bars have higher similarity. In UMAP scatter plot in the center of the circular bar chart, each point represent a gene. The colors of the genes represent the grouping of genes. Genes that are close to each other in the UMAP also indicate a higher functional correlation. Red lines connect each gene to its corresponding enriched term.**

FigureS27

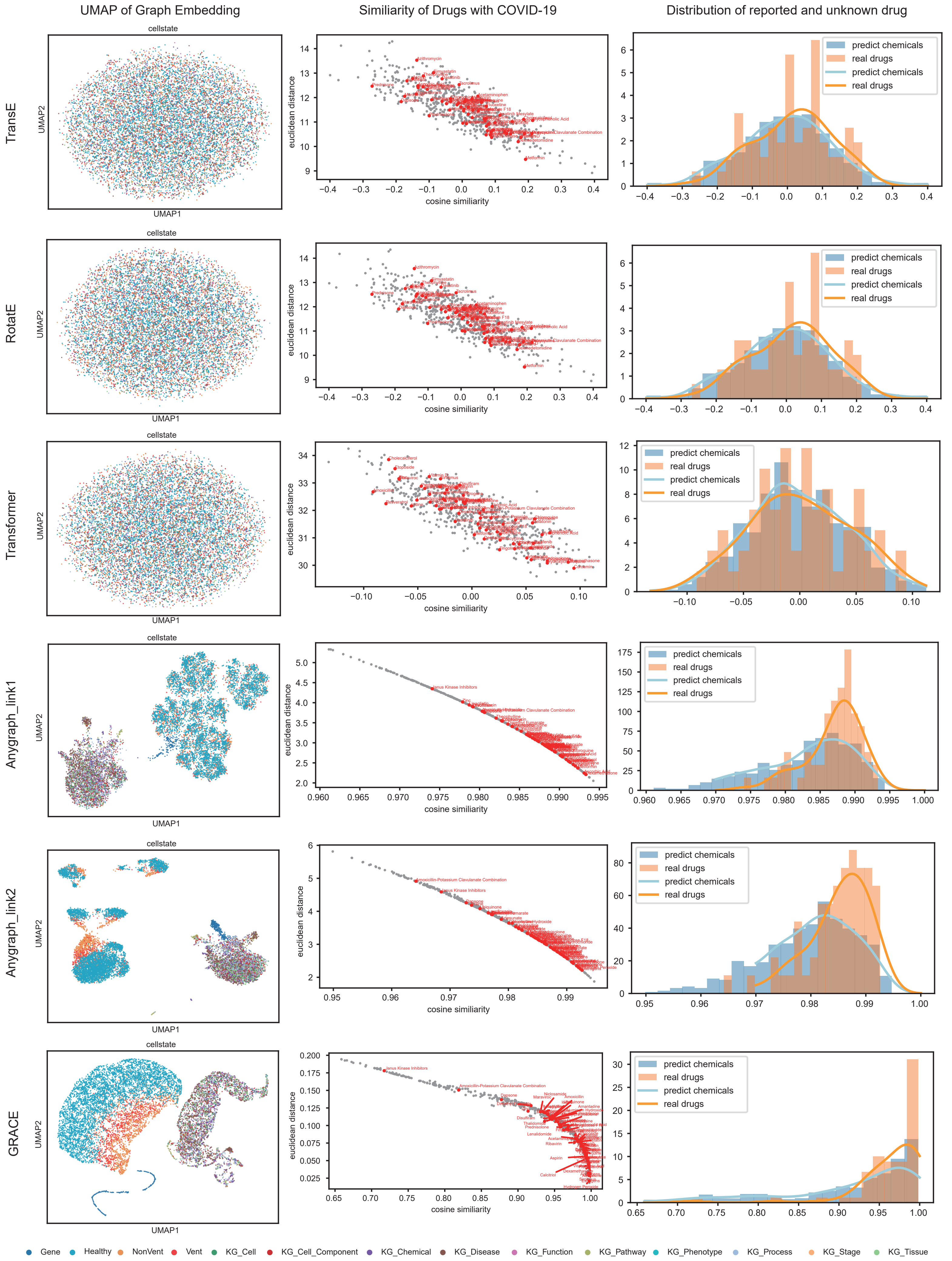


**Supplementary Fig S27. NK cell representative learning for drug repurposing based on different metrics. left column: UMAP of graph embedding of knowledge entities and omics cells and genes. middle columns: scatter plot of distance and similarity between chemicals and COVID-19, x-axis: cosine similiarity, y-axis: euclidean distance, reported COVID-19 chemicals are marked and labeled in red. right columns: hist gram of cosine similarity of reported chemicals and other chemicals. From top to the bottom, the graph embedding metric is TranE, RotatE, Transformer, AnyGraph with first pretrained parameters, AnyGraph with second pretrained parameters, GRACE, respectively.**

FigureS28

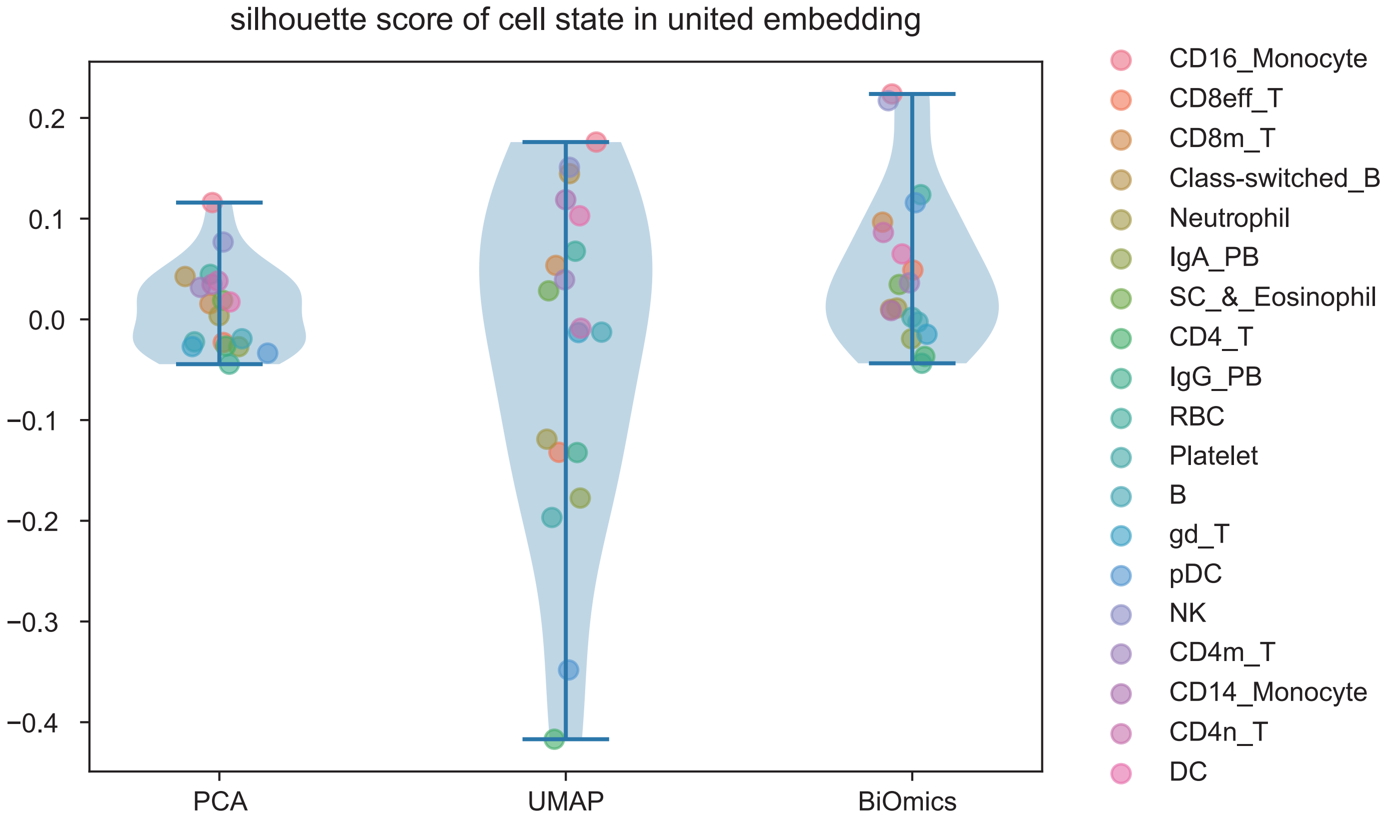


**Supplementary Fig S28. Violine plot sihouette score of cell state in united embedding. jitter indicate differential cell types.**

FigureS29

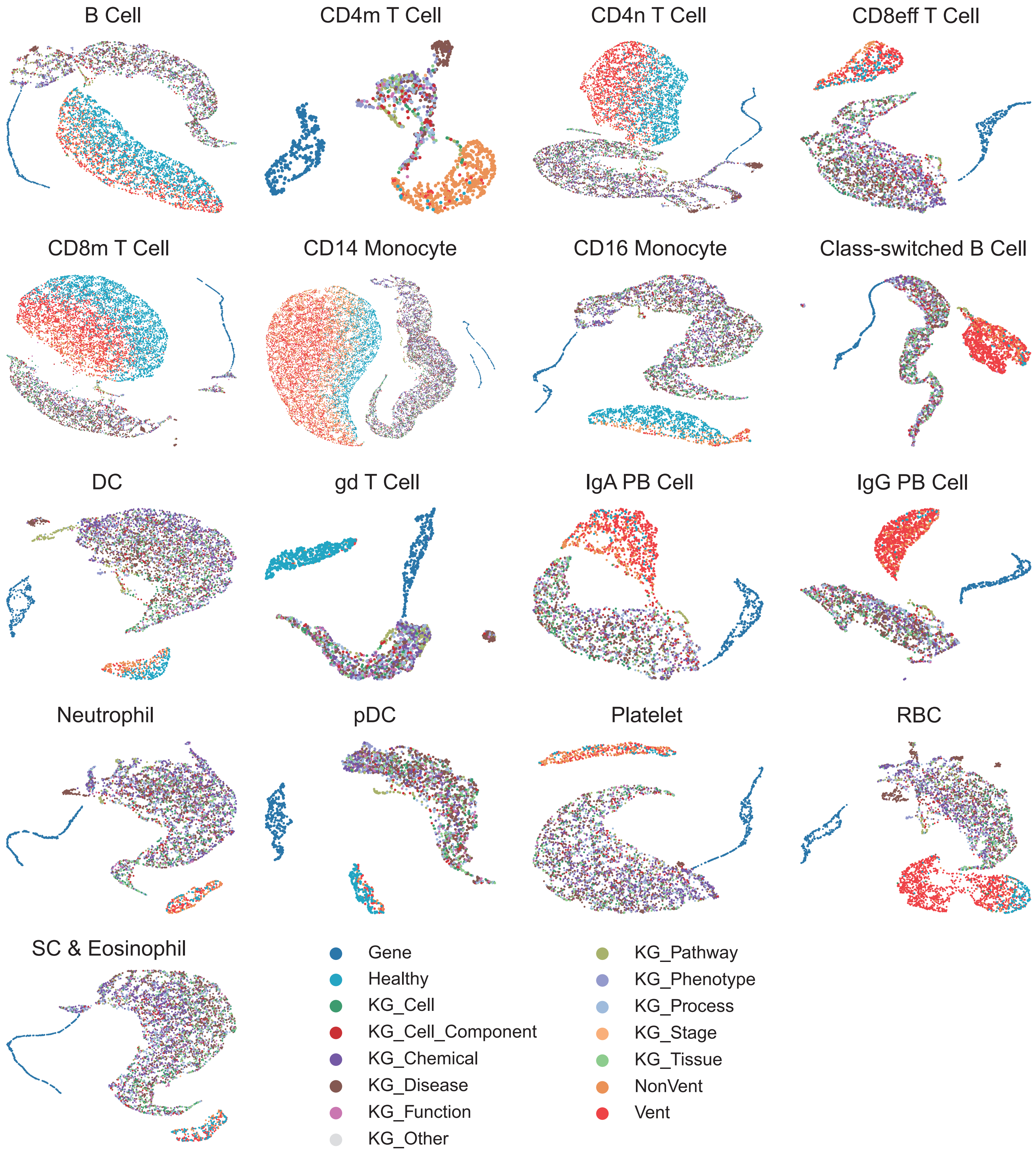


**Supplementary Fig S29. UMAP of graph embedding of knowledge entities and omics cells and genes for differential cell types. Corresponding celltype is labeled on the top of each subplot.**

FigureS30

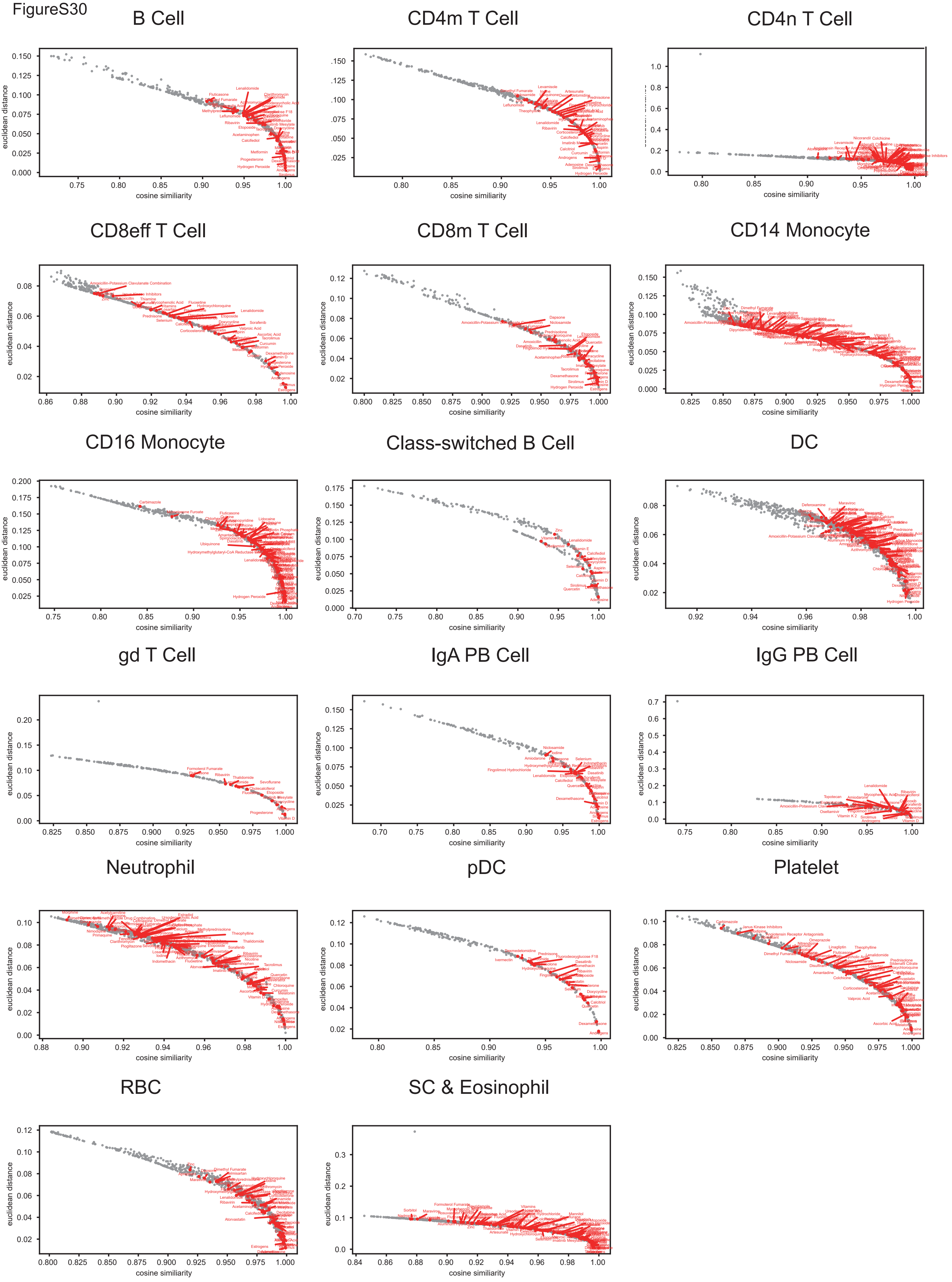


**Supplementary Fig S30. Scatter plot of distance and similiarity between chemicals and COVID-19 for differential cell types. Corresponding celltype is labeled on the top of each subplot.**

FigureS31

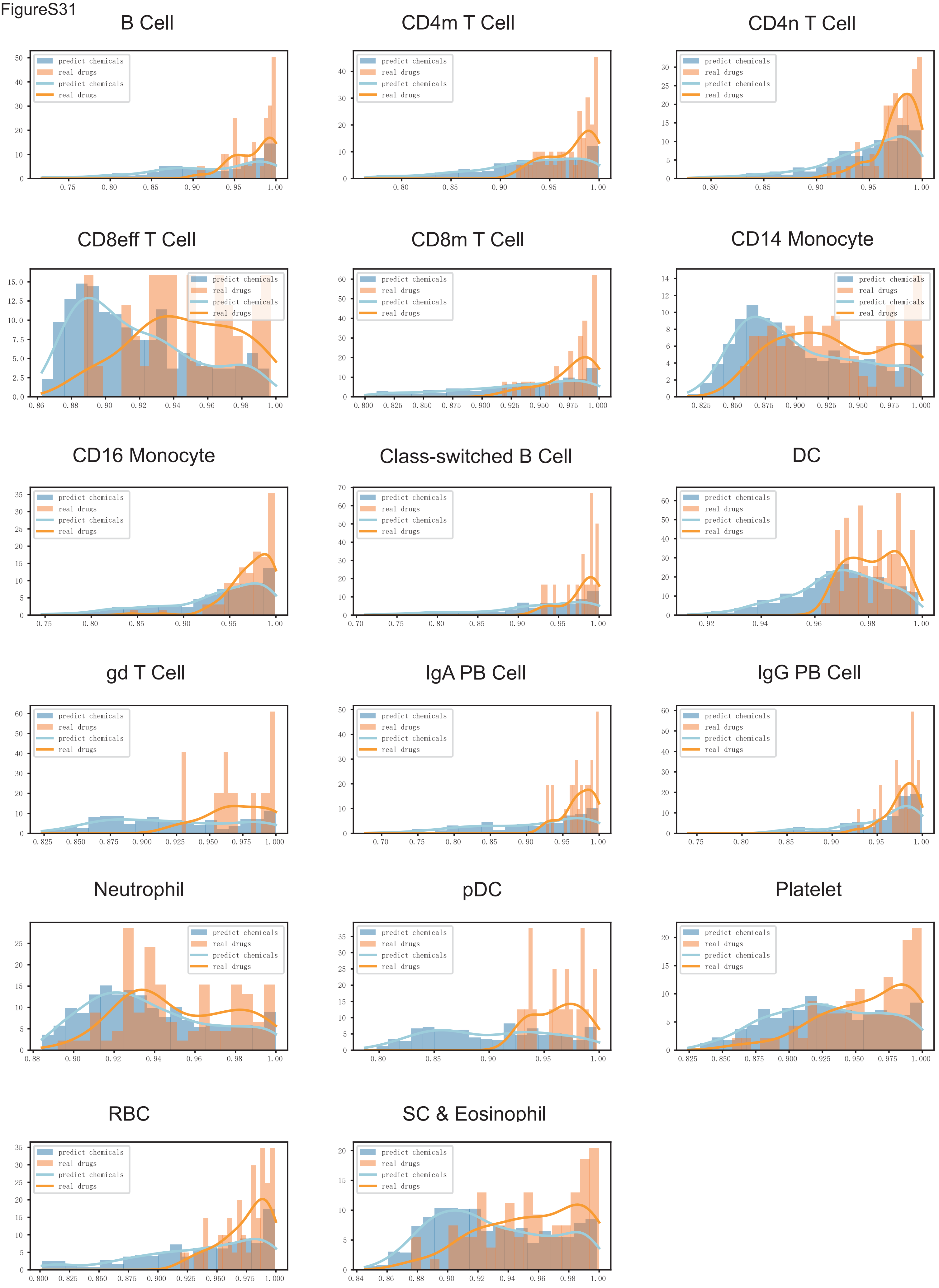


**Supplementary Fig S31. Histgram of cosine similarity of reported chemicals and other chemicals for differential cell types. Corresponding celltype is labeled on the top of each subplot.**

FigureS32

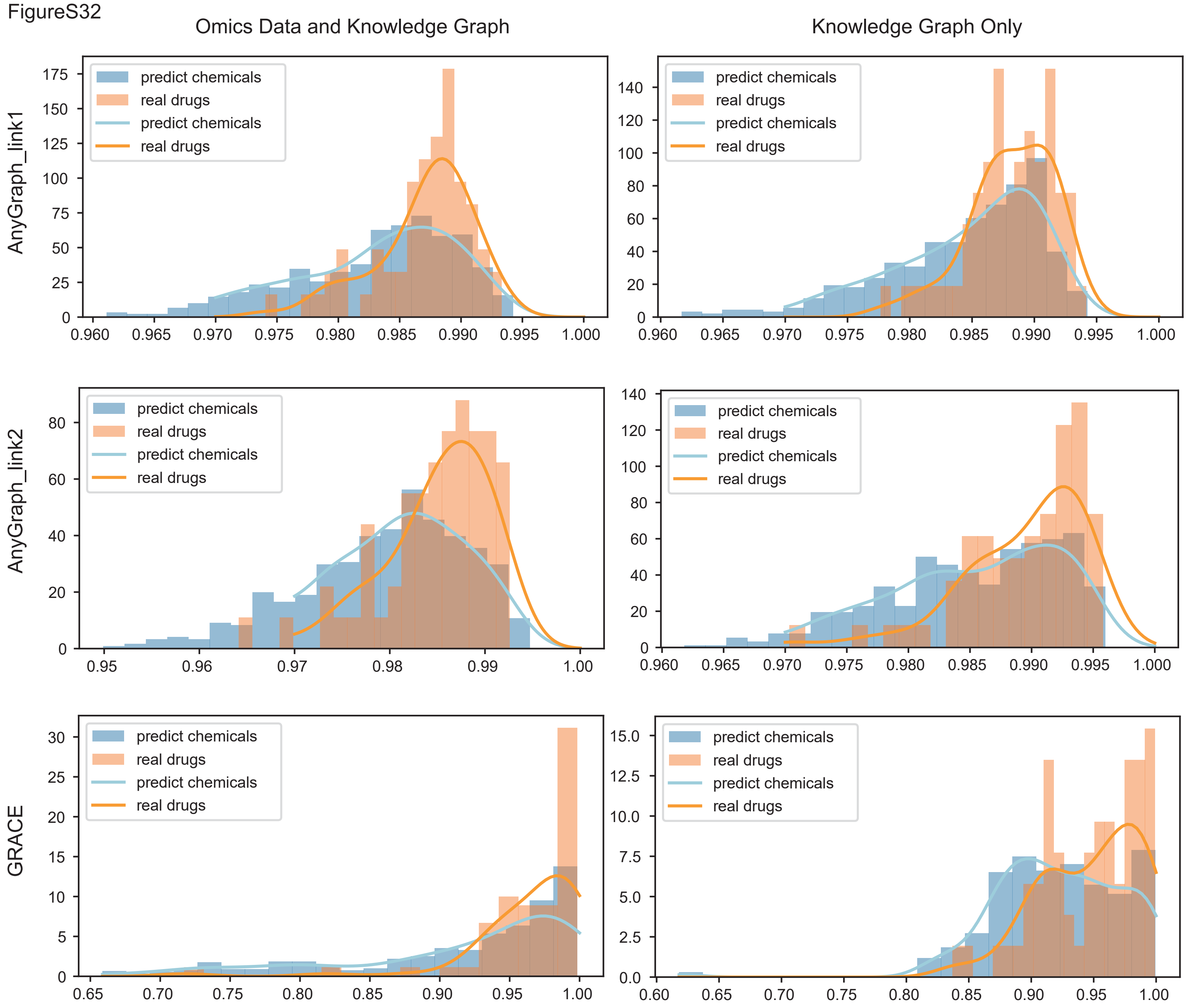


**Supplementary Fig S32. Histgram of cosine similarity of reported chemicals and other chemicals comparing the graph with omics data and knowledge graph only. left column: omics data with knowledge graph, right columns: knowledge graph only, From top to the bottom, the graph embedding metric is AnyGraph with first pretrained parameters, AnyGraph with second pretrained parameters, GRACE, respectively.**

FigureS33

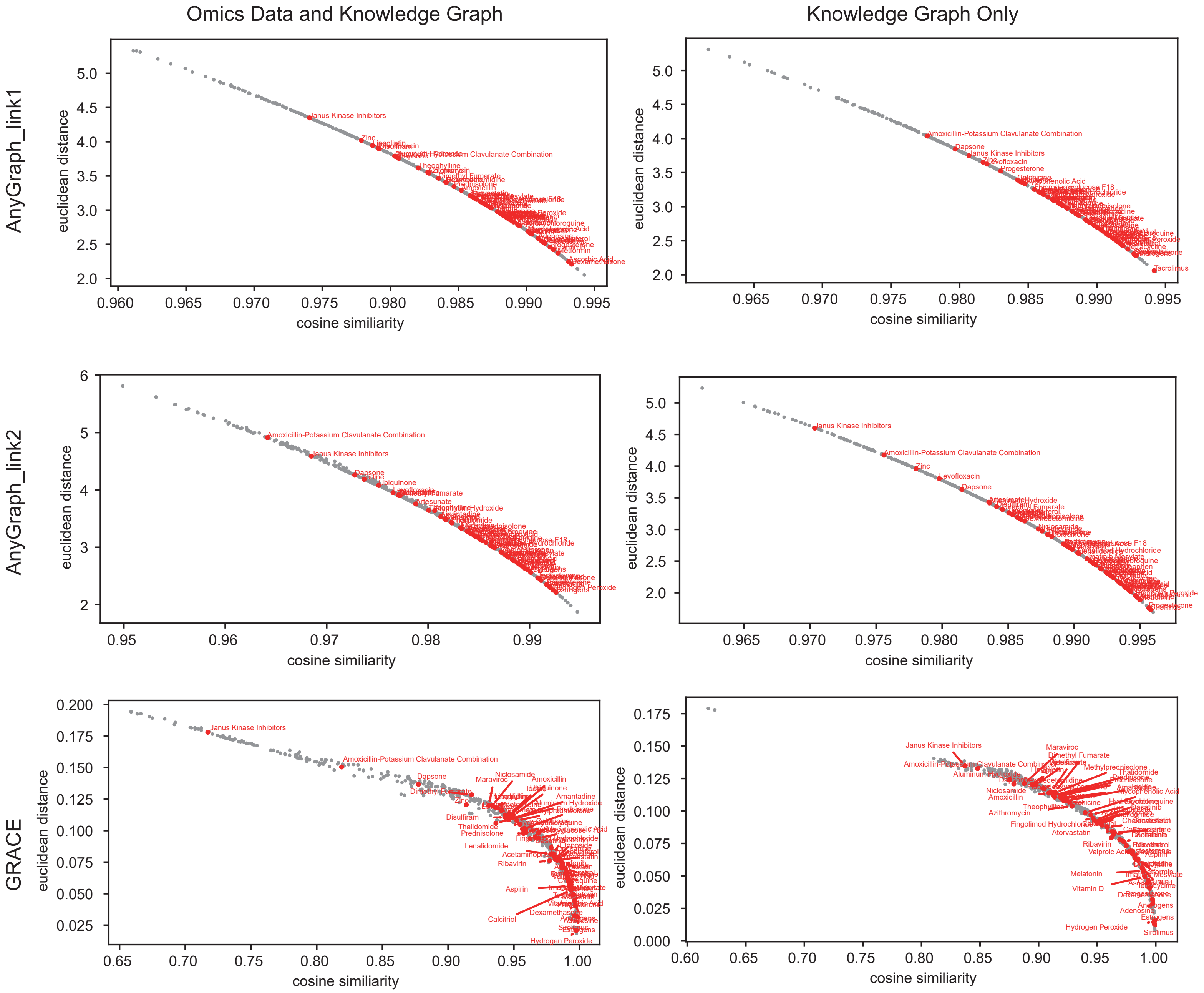


**Supplementary Fig S33. Scatter plot of distance and similiarity between chemicals and COVID-19 comparing the graph with omics data and knowledge graph only. x-axis: cosine similiarity, y-axis: euclidean distance, reported COVID-19 chemicals are marked and labeled in red. left column: omics data with knowledge graph, right columns: knowledge graph only, From top to the bottom, the graph embedding metric is AnyGraph with first pretrained parameters, AnyGraph with second pretrained parameters, GRACE, respectively.**

FigureS34

### Interpretation of Drug Repurposing Results for COVID-19

#### Summary of Evidenced Drugs
Based on the `evidenced_drug` column in Table 1, the following drugs have been identified as having evidence for use in treating COVID-19:
- **Adenosine**
- **Dexamethasone**
- **Calcitriol (Vitamin D)**
- **Aspirin**
- **Imatinib Mesylate**
- **Ribavirin**
- **Chloroquine**
- **Tacrolimus**
- **Lenalidomide**
- **Corticosterone**
- **Cholecalciferol (Vitamin D3)**
- **Calcifediol (Vitamin D2)**
- **Tetracycline**
- **Atorvastatin**
- **Fingolimod Hydrochloride**
- **Simvastatin**
- **Etoposide**
- **Sorafenib**
- **Hydroxychloroquine**

These drugs have shown potential or have been used in clinical settings for the treatment of COVID-19. They are already known to have some level of efficacy or are being studied for their potential benefits.

#### Candidate Drugs Without Evidence
The following candidate drugs, ranked by cosine similarity, do not have evidence for use in treating COVID-19 but show high similarity and may be worth further investigation:

1. **Retinoic Acid (Cosine Similarity: 0.993763)**
   - **Rationale**: Retinoic acid is a metabolite of vitamin A and has been shown to have anti-inflammatory and immunomodulatory effects. Given that COVID-19 often involves a hyperinflammatory response, retinoic acid could potentially help modulate this response and reduce the severity of the disease.

2. **Paclitaxel (Cosine Similarity: 0.990273)**
   - **Rationale**: Paclitaxel is an antineoplastic agent that stabilizes microtubules and inhibits cell division. While it is primarily used in cancer treatment, its mechanism of action might also have implications for viral replication and host cell interaction, making it a potential candidate for further study in the context of COVID-19.

3. **Cisplatin (Cosine Similarity: 0.992347)**
   - **Rationale**: Cisplatin is another antineoplastic agent that forms intra- and inter-strand cross-links in DNA, leading to apoptosis. Its potential to disrupt viral replication or host cell processes could make it a candidate for further investigation in the context of COVID-19.

4. **Fluorouracil (Cosine Similarity: 0.986223)**
   - **Rationale**: Fluorouracil is an antimetabolite that interferes with DNA and RNA synthesis. It is used in cancer treatment and could potentially inhibit viral replication, making it a candidate for further study in the context of COVID-19.

5. **Cyclosporine (Cosine Similarity: 0.979992)**
   - **Rationale**: Cyclosporine is an immunosuppressive agent that inhibits T-cell activation. Given the role of the immune system in the pathogenesis of severe COVID-19, cyclosporine could potentially help modulate the immune response and reduce the severity of the disease.

6. **Bortezomib (Cosine Similarity: 0.976663)**

**Supplementary Fig S34. BiOmics interpretation context of drug repurpose analysis based on NK cell.**
